# Every cog and wheel: Identifying biocomplexity at the genomic and phenotypic level in a population complex of Chinook salmon

**DOI:** 10.1101/2021.03.26.437213

**Authors:** Shannon J. O’Leary, Tasha Q. Thompson, Mariah H. Meek

## Abstract

Genomic diversity is the fundamental building block of biodiversity and the necessary ingredient for adaptation. Our rapidly increasing ability to quantify functional, compositional, and structural genomic diversity of populations forces the question of how to balance conservation goals – should the focus be on important functional diversity and key life history traits or on maximizing genomic diversity as a whole? Specifically, the intra-specific diversity (biocomplexity) comprised of phenotypic and genetic variation can determine the ability of a population to respond to changing environmental conditions. Here, we explore the biocomplexity of California’s Central Valley Chinook salmon (*Oncorhynchus tshawytscha*) population complex at the genomic level. Notably, despite apparent gene flow among individuals with the same migration (life history) phenotypes inhabiting different tributaries, each group is characterized by a surprising component of unique genomic diversity. Our results emphasize the importance of formulating conservation goals focused on maintaining biocomplexity at both the phenotypic and genotypic level. Doing so will maintain the species’ adaptive potential and increase the probability of persistence of the population complex despite changing environmental pressures.

## Introduction

> *“The last word in ignorance is the man who says of an animal or plant, “What good is it?”[…] If the biota, in the course of aeons, has built something we like but do not understand, then who but a fool would discard seemingly useless parts? To keep every cog and wheel is the first precaution of intelligent tinkering.”*
>
> — Aldo Leopold

Genetic diversity represents the fundamental building block of biodiversity – from genes to species, to communities and ecosystems. The genetic diversity contained within populations forms the cogs and wheels comprising the adaptive potential of a population. Protecting this diversity is a central task of conservation biology. However, identifying and quantifying biodiversity is a complex undertaking with distinct frameworks for enumerating compositional, structural, and functional diversity (Petchey & Gaston 2002; Duelli & Obrist 2003; Péru & Dolédec 2010). Functional genetic diversity includes both sequence polymorphisms and differences in gene expression, which together shape the phenotypic diversity comprised of differences in morphology, physiology, and life history characteristics present within a population. Under current environmental conditions, it is expected that some components of functional diversity will be selectively neutral, but future shifts in conditions and evolutionary pressures may result in certain phenotypes and genotypes that appeared unimportant under previous conditions becoming critical to the persistence of a species or population (Messer et al. 2016). Therefore, in order to maintain critical evolutionary and ecological processes that sustain biodiversity across scales, sound conservation and management strategies require a fundamental understanding of both past and present patterns of biodiversity to enable the protection of diversity that will form the building blocks for future adaptation (CBD 2011; Hoelzel et al. 2019; Mable 2019).

Increasingly, the importance of intra-specific diversity (biocomplexity) has been recognized as a determining factor for the stability and resilience of biological systems (Hilborn et al. 2003; Des Roches et al. 2021). For example, at an ecosystem level, the number of trophic levels and number of species at each level determines the stability of the food web, while at a species level, diversity of life history strategies can be critical for maintaining a temporally stable population through risk partitioning (i.e., the portfolio effect (Hilborn et al. 2003; Schindler et al. 2010)). While biocomplexity is frequently viewed as a horizontal measure within a given level of biodiversity, it cannot always be neatly confined to a single hierarchical level. For example, while intra-specific diversity at a population-level is comprised of phenotypic diversity, there is a genetic component dictating some phenotypic differences. Similarly, species interactions not only shape an ecosystem but simultaneously act as evolutionary pressures determining the fitness of individual phenotypes. Therefore, because selection acts on standing variation, forward-looking conservation strategies must stress the importance of maintaining adequate levels of genetic diversity within a population to maintain adaptive potential in changing environmental conditions (Mimura et al. 2017; Mable 2019; Hoban et al. 2020; Des Roches et al. 2021).

Chinook salmon, *Oncorhynchus tshawytscha*, in the Central Valley are emerging as a case study to understanding the importance of biocomplexity for the persistence of a population complex facing multiple external threats, including habitat fragmentation, overexploitation, and climate change. Here, life history diversity resulting from different migration phenotypes creates a “portfolio of stocks” buffering against spatiotemporally variable environmental conditions and anthropogenic impacts, which in turn increases resilience and promotes interannual stability (Carlson & Satterthwaite 2011; Griffiths et al. 2014). Chinook are anadromous with a distinct life history spanning both freshwater and marine ecosystems. Eggs are laid and hatched in the tributaries, where juveniles rear for a period of time before migrating out to the ocean. There, they spend one to several years growing in the ocean before migrating back to their natal river to spawn and die, providing an important source of oceanic nutrients to ecosystems and supporting recreational, commercial, and heritage fisheries (Quinn 2018). The tributaries of the Sacramento-San Joaquin River system contain four distinct run types (migration phenotypes) named for the time of year adults enter freshwater systems to spawn: Winter (endemic to the Central Valley), Spring, Fall, and Late-Fall; the same tributary may support multiple runs (Williams 2006). Early-migrating runs (Winter, Spring) make the trade-off of migrating earlier at a smaller size, leaving behind the nutrient-rich oceanic habitats to access spawning sites higher in the watershed that remain cool over the summer, where they complete maturation in a fasted state while relying on fat stores, spawn, and die (Quinn et al. 2016). By contrast, late-migrating individuals (Fall, Late-Fall) remain in ocean until relatively mature before making their spawning run.

This asynchronicity in run timing of adults stabilizes the population complex overall, as temporal and spatial variation in challenges to population components will vary. For example, while environmental conditions within a given year may be poor for early migrating adults, in that same year they may be optimal for their late-migrating counterparts in the same tributary, thus buffering the population complex overall. However, this buffering ability is threatened when one run-type is consistently negatively impacted across tributaries. Specifically, dams and other anthropogenic factors have disproportionately affected historical early migrator habitat in much of the Central Valley. As a result both Spring and Winter run are listed under the Endangered Species Act (National Marine Fisheries Service 2005) and the dramatic declines in their abundance and distribution has resulted in a loss of biocomplexity across run types, thus lowering the portfolio effect and making the population complex as a whole more vulnerable (Carlson & Satterthwaite 2011).

These demographic changes are also likely to result in an erosion of genetic diversity and consequently adaptive potential due to low effective sizes, which may otherwise have proved important for persistence under future environmental conditions. Measuring the genetic diversity of a population has long been used as a proxy to quantify the “future adaptive potential” of populations (Reed & Frankham 2001, 2003). However, despite a plethora of tools to quantify genetic diversity, consensus on which part of the genetic diversity to focus on for conservation remains elusive. By focusing on the structural biodiversity at a genomic level, tools such as diagnostic single nucleotide polymorphism (SNP) panels for Chinook can be used as intrinsic markers to identify (hierarchical sets of) demographic groups, quantify biocomplexity across a species’ range, and describe changes in the portfolio of stocks (Meek et al. 2016). Recent technological advances now also allow for the direct interrogation of functional biodiversity at the genomic level. For example, recent studies have identified a single chromosomal region (GREB1L to ROCK1) underlying adult migration timing in Chinook and other salmonids (Prince et al. 2017; Narum et al. 2018; Thompson et al. 2019, Thompson et al. 2020). In addition, the increasing availability of genomic data sets holds promise for identifying genes associated with polygenic traits (Ouborg et al. 2010;

Sinclair-Waters et al. 2020). Finally, focusing on the compositional biodiversity at a genomic level will lead to an increased understanding of the differences in genetic diversity and unique components among populations within a complex.

The increased availability of data quantifying functional, compositional, and structural genomic diversity of populations forces questions about what is important to consider for conservation. There is a need for balancing goals focused on preserving important functional diversity and key life history traits with goals aimed at preserving adaptive potential that may be more cryptic, especially when putatively neutral and adaptive markers display divergent signals (Waples & Lindley 2018). This is highlighted by recent studies that identified a major effect locus underlying adult run timing in Chinook salmon (Prince et al. 2017; Narum et al. 2018; Thompson et al. 2019; Ford et al. 2020). The large effect size of this locus (e.g., explaining >85% of the variance in freshwater entry timing; (Thompson et al. 2020) combined with widespread declines and extirpations of spring-run populations and the finding that spring-run alleles are not preserved in the absence of the phenotype (Thompson et al. 2019; Ford et al. 2020) has led to debate on the extent to which conservation policy should address this single locus (Langin 2018; Ford et al.

2020). The debate is additionally complicated because it is poorly understood to what extent adaptive variation beyond this locus may be impacted by loss of the spring-run life history. For example, spring- and fall-run fish historically utilized largely disparate spatio-temporal habitats (e.g., spring-run typically spawned earlier and higher in watersheds; Quinn et al. 2016), which could facilitate the development of important local adaptative differences between the runs. However, human activities have homogenized habitat and substantially increased interbreeding between spring- and fall-run Chinook in many locations (Ford et al. 2020), and it is unclear whether local adaptation has been preserved. Despite this, the remaining wild spring-run populations in California’s Central Valley still access very distinct habitat from their fall-run counterparts. Furthermore, a great deal of habitat heterogeneity exists *within* the spring-run. Thus, the Central Valley provides an opportunity to examine the extent to which additional unique genetic variation may accumulate between runs and across runs in a population complex where spatio-temporal separation between runs is relatively intact.

Here, we aim to understand the genomic biocomplexity, contained within and among the four Central Valley Chinook salmon run types. To achieve this, we leverage a previously published dataset (Meek et al. 2019). The exceeding value of this data set lies in the fact that it encompasses genomic sequences from all the major populations of the critically endangered and threatened Central Valley Chinook salmon, including winter run, which is on the verge of extinction. The analysis of Meek et al. (2019) demonstrated greater population diversity and structure across the Central Valley than had been previously described but lacked the genomic resources and analytical methods needed to elucidate the genomic biocomplexity contained within and among populations. This understanding is vital for informed conservation planning that will protect the full portfolio of diversity and better ensure long term persistence and resilience. In this paper, we resolve this problem by using the recently completed Chinook salmon genome (Christensen et al. 2018) and a novel microhaplotype analysis (Willis et al. 2017) to identify > 10,000 multi-allelic loci distributed throughout the coding and non-coding parts of the Chinook genome. Our results not only provide new insight about this important and highly imperiled species, but also demonstrates the power of applying new advances to existing dataset to gain vital biological understanding, without the need for resampling imperiled populations.

## Materials and Methods

### Sample collection and sequencing

As reported in Meek et al. 2019, fin clips from all four run types (Fall, Late-Fall, Winter, Spring; Figure 1) were obtained from adult Chinook Salmon from each major tributary in the Central Valley during their spawning migrations. Genomic DNA was extracted and digested using *SbfI* to construct RAD libraries following Miller et al. (Miller et al. 2012). Fifteen libraries consisting of 30 – 47 individuals each were sequenced on single Illumina HiSeq 2000 lanes (100bp, single end).

**Figure 1:**
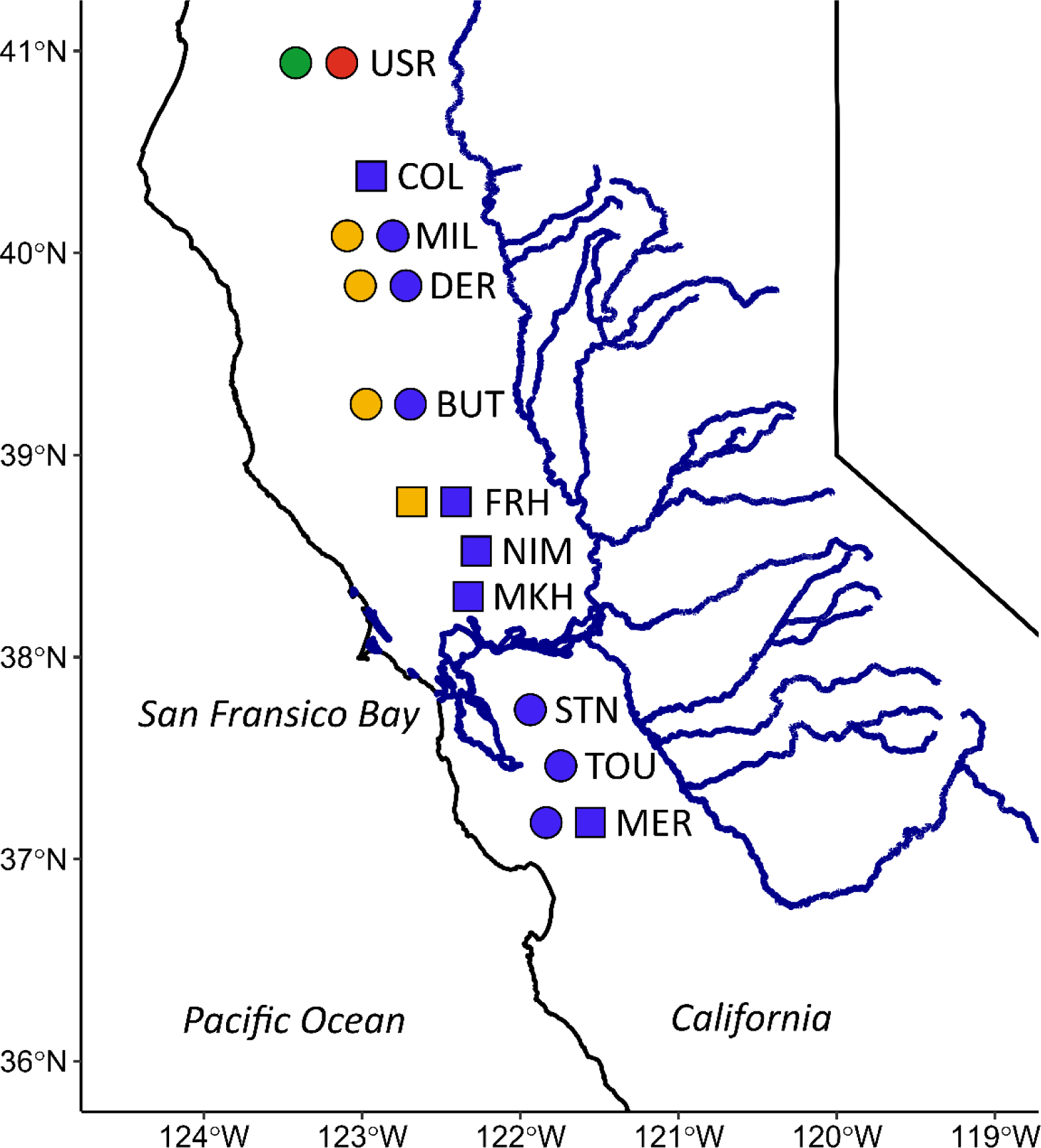
Tributaries of the California’s Central Valley sampled for this study. Hatchery individuals are represented by squares and wild populations by circles. Colors represent the sampled run type at each location (Spring = green, Fall = blue, Late-Fall = red, Winter = yellow). Abbreviations for tributaries used throughout: USR (Upper Sacramento River), COL (Coleman Hatchery/Battle Creek), MIL (Mill Creek), DER (Deer Creek), BUT (Butte Creek), FRH (Feather River Hatchery), NIM (Nimbus Hatchery/American River), MKH (Mokelumne River Hatchery), STN (Stanislaus River), TOU (Tuolumne River), MER (Merced River).

**Figure 2:**
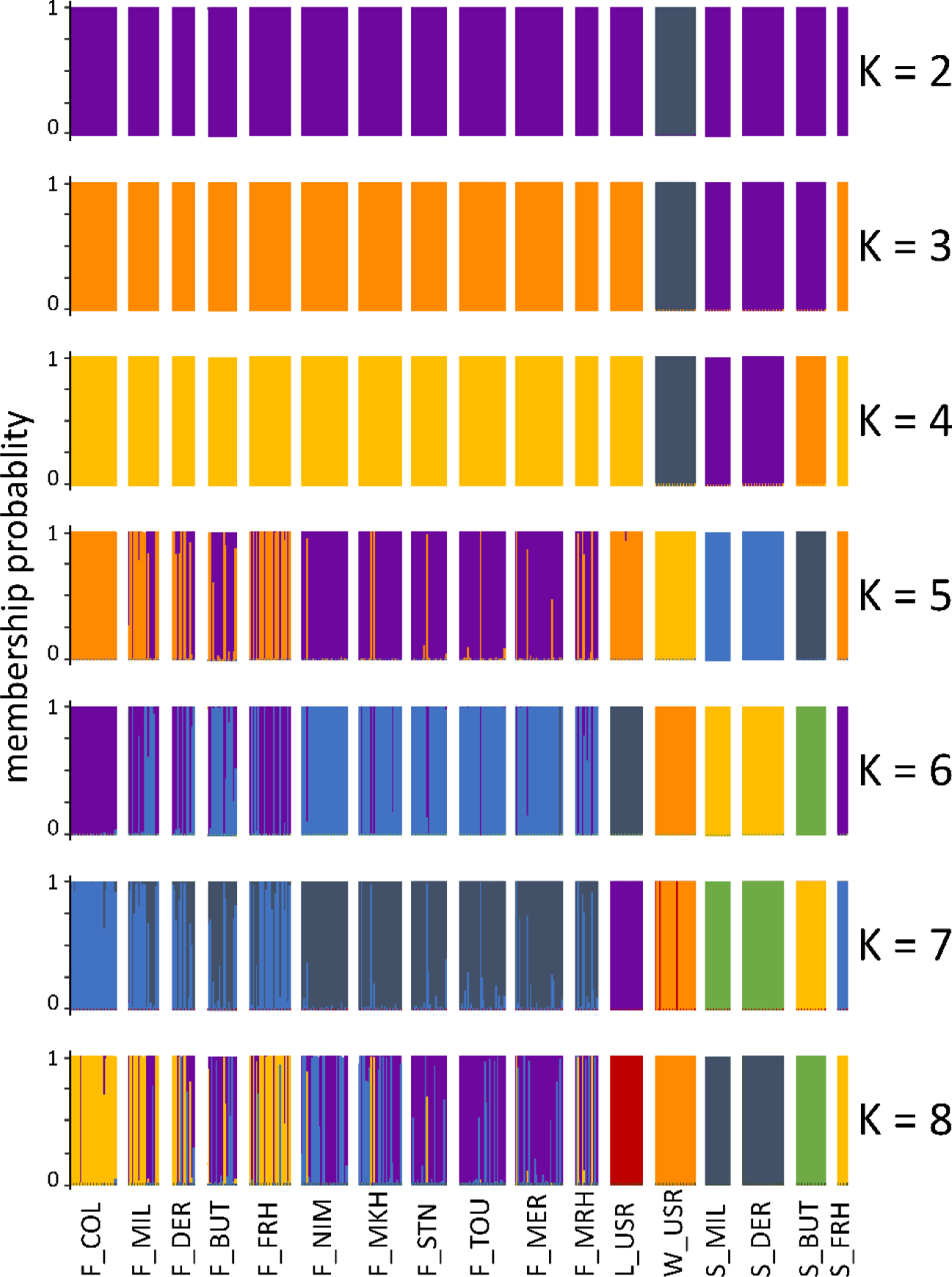
Membership probability of each individual to clusters identified using k-means hierarchical clustering for K = 2 – 8.

### Genotyping

Raw sequences from Meek et al. 2019 were demultiplexed using *process_radtags* (Catchen et al. 2011) and quality trimmed using *fastp* (Chen et al. 2018). Quality trimmed reads were mapped to a Chinook reference genome (Christensen et al. 2018) using *BWA-mem* (Li & Durbin 2009). Reads with a mapping quality < 5 were removed and filtered bam files were concatenated into a single bam file to query coverage per interval using *bedtools* (Quinlan 2014). Mapping intervals > 25bp and < 500bp with coverage > 50 were extracted to create a reduced-representation reference consisting of the recovered RAD-tags, reads were re-mapped to the reduced-representation reference and SNPs were called using *freebayes;* the maximum allowed gap (-E) was set to three and minimum mapping and base quality set to five, otherwise default parameters were used (Garrison & Marth 2012).

The raw data set was rigorously filtered to remove low quality genotypes, loci, and samples following principles set forth in O’Leary et al. (2018). In short, genotypes with < 5 reads or quality < 20 were coded as missing, retained loci had a mean minimum depth of 15, and were called in at least 50% of individuals of a given library, 85% of individuals of a run type, or 90% of individuals overall. Loci were further filtered for allele balance, mapping quality ratio of reference vs alternate allele, depth/quality ratio, and excess heterozygosity to remove paralogs and other technical artifacts. Finally, *rad_haplotyper* (Willis et al. 2017) was used to collapse SNPs on the same RAD-tag into haplotypes. Detailed processing steps and threshold values are contained in Supplementary Material 1. In general, biallelic SNPs contain less information per locus compared to multi-allelic loci such as microsatellites (Morin et al. 2009) and the necessity of thinning SNPs to ensure loci are independent observations (Kaeuffer et al. 2007) further reduces the information content of a data set as the power of a data set resides in the number of independent alleles rather than the number of loci (Kalinowski 2002). Haplotyping a data set rather than thinning preserves the information content of all SNPs in the data set, resolves physical linkage artefacts, and results in more inferential power per locus (Willis et al. 2017; Baetscher et al. 2018). Therefore, this new dataset has much more power to identify genomic diversity and biocomplexity.

### Assessment of population structure and differentiation

Population structure was explored using two methods, a clustering analysis based on genetic similarity and an assessment of population differentiation among individuals grouped *a priori* based on run type and tributary. In the first method, individuals were clustered into K = 1 – 10 groups using k-means clustering based on the PCA-transformed genotype matrix (i.e. no assumptions regarding Hardy-Weinberg or linkage disequilibrium) followed by a discriminant analysis of principle components (DAPC) to determine the membership probabilities of each sample to each inferred cluster as implemented in *adegenet* (Jombart et al. 2010). To ensure sufficient variance was retained to discriminate among groups but not overfit the data, the optimum number of principle components to retain was determined using a stratified cross-validation of DAPC.

In the second method, Weir & Cockerham’s unbiased estimator of *F*ST (Weir & Cockerham 1984) was used to calculate population differentiation among individuals grouped *a priori* by run type within tributary. To test for genetic heterogeneity in the data set, global *F*ST was calculated across all groups, then, pairwise *F*ST was calculated as a *post hoc* test for pairwise differences among groups. Significance was determined using 95% confidence intervals around each estimate generated by resampling loci 1,000 times using *assigner* (Gosselin et al. 2016).

### Estimates of effective population size

Estimates of effective population size, *N*e, for each tributary/run group were determined using *LinkNe* (Hollenbeck et al. 2016), an extension of the linkage disequilibrium method (LD) designed for data sets with loci of known linkage relationships. Only SNPs placed on a chromosome, with a minor allele frequency > 0.01, and the first SNP per RAD-tag were used. Genomic distances (bp) were converted to recombination rates (cM) using the size of the female Chinook linkage map (3,118 cM, (Mckinney et al. 2016)) and the length of the genome (2.4 Gbp, (Christensen et al. 2018)) resulting in an estimate of approx. 770kb equivalent to 1cM.

### Assessment of Genomic Diversity

All measures of genomic diversity were made for individuals grouped by run type within each tributary, with wild and hatchery individuals treated as separate groups. Four types of parameters were assessed, (1) measures of heterozygosity, (2) measures of allelic diversity, (3) sequence-based parameters, and (4) measures of unique variation.

For the first three sets of parameter types, significant heterogeneity was determined using a Friedman’s rank sum test followed by a *post hoc* Wilcoxon signed-rang test to test for significant pairwise differences between groups; *p*-values were corrected for multiples comparisons assuming an FDR of 0.05 (Benjamini & Hochberg 1995). The observed heterozygosity (Ho) was measured as the proportion of heterozygote genotypes per locus (Nei 1987) and the expected heterozygosity (gene diversity) (Hs) as the proportion of heterozygous genotypes expected under Hardy-Weinberg Equilibrium (Nei 1987). To account for differences in sample size, allelic richness was measured as rarefied allele counts. The evenness of allelic diversity at a given locus was calculated as the ratio of the number of abundant to the number of more rare genotypes using the ration of the Stoddart & Taylor index (diversity index weighted for more abundant alleles) and Shannon-Wiener index (diversity weighted for more rare alleles) as implemented in *poppr* (Kamvar et al. 2014); lower values indicate prevalence of more rare alleles and uneven distributions of allele frequencies. The nucleotide diversity statistic π (Nei 1987) was calculated as the sum of the number of pairwise differences between haplotypes of a given nucleotide over the number of comparisons made; this parameter is biased toward alleles segregating at intermediate rates and will underestimate genetic diversity when many rare alleles are present.

Finally, patterns of unique diversity were assessed by comparing (1) the number of fixed loci (i.e. not polymorphic within a group), (2) the number of loci with singletons and the number of singletons per individual by run/tributary group, (3) the number of private polymorphisms, and (4) the number of private alleles. Private polymorphisms are defined as loci where more than one allele is found only in a single group (all other groups are fixed for a single allele). By contrast, private alleles are alleles found in only a single group, though other groups exhibit more than one allele at that locus. To compare whether identified loci are randomly distributed across chromosomes, null distributions were generated by shuffling chromosome designations across loci 1,000 times to determine whether the observed values fall outside the null distribution.

## Results

### Genotyping

The final filtered data set consisted of 386 individuals genotyped for 12,983 multi-allelic loci (hereafter loci) with a total of 30,037 alleles (Table 1).

**Table 1:** Sample sizes and effective population size Ne for all run/tributary groups. Negative point estimates occur when LD attributed to sampling variation is larger than LD attributed to drift. This can either be interpreted as an “infinite” population size (drift is negligible) or due to sample size being too small for an accurate estimate (LD attributed to sampling variation is a function of sample size).

| Run | Tributary/Hatchery | Sample Size | $N_e$ (95% CI) |
| --- | --- | --- | --- |
| Fall | Coleman Hatchery | 30 | 615 (594; 637) |
|  | Mill Creek | 20 | -6,801 (-56,952; -3,618) |
|  | Deer Creek | 15 | -3,939 (-9,323; -2,497) |
|  | Butte Creek | 21 | 3,802 (2,708; 6,378) |
|  | Feather River Hatchery | 27 | 5,172 (3,823; 7,988) |
|  | Nimbus Hatchery | 30 | 7,432 (5,147 – 13,351) |
|  | Mokelumne River Hatchery | 28 | 4,106 (3,207; 5,704) |
|  | Stanislaus River | 23 | 2,567 (2,040; 3,460) |
|  | Tuolumne River | 30 | 6,353 (4,383; 11,538) |
|  | Merced River Hatchery | 15 | 1,125 (940; 1,399) |
|  | Merced River | 31 | 11,436 (6,768; 36,804) |
| Late-Fall | Upper Sacramento River | 21 | 3,114 (2,398; 4,435) |
| Winter | Upper Sacramento River | 26 | 174 (171; 177) |
| Spring | Mill Creek | 16 | 85 (84; 86) |
|  | Deer Creek | 27 | 188 (185; 191) |
|  | Butte Creek | 19 | 561 (529; 597) |
|  | Feather River Hatchery | 7 | -94 (-98; -91) |

### Assessment of population structure and differentiation

Minimum AIC was observed for K = 4 (Figure S1); cross-validation indicated that the optimum number of principle components to retain was 20 – 150 (retaining 11.6 – 75.1% of variance; Table S1). The mean optimum success of assignment declined from 100% (K = 2 – 3) to 96% (K = 7) and then dropped to 89% for K = 8 (Table S1).

Figure 1 summarizes membership plots for K = 2 – 8. For K = 2 Winter run individuals form a single cluster set against individuals from all other run types and tributaries. Similarly, for K = 3, Winter run form a single cluster, a second cluster consists of Spring run individuals from Butte, Deer, and Mill Creek, while Spring Feather River Hatchery and Late-Fall individuals form the final cluster along with all Fall groups. Overall, Late-Fall individuals do not emerge as their own distinct cluster until K = 6, while Spring Feather River individuals continue to cluster with Fall individuals from Coleman and Feather River Hatchery for K = 7 – 8. Overall, Spring run individuals form at most three clusters, with Deer and Mill Creek individuals always clustering together. In general, Fall run individuals are assigned to two or three clusters, here, Coleman Hatchery individuals generally form one cluster along with most Feather River Hatchery individuals (both Fall and Spring) and some Deer and Mill Creek individuals; increasing K to 8 results in Coleman Hatchery individuals starting to form a more distinct cluster of their own.

For individuals grouped by run type within tributaries, global *F*ST = 0.0319 [0.0309 – 0.0328] indicates significant heterogeneity among groups. Pairwise comparisons of Winter individuals and all other run/tributary groups exhibit the highest observed pairwise *F*ST-values (0.14 - 0.161). By contrast, all non-significant comparisons, with CIs including zero, were comparisons among Fall run tributary populations. Late-Fall Upper Sacramento River individuals are significantly different from all other run/tributary groups. For Late-Fall and Winter individuals pairwise *F*ST is higher (0.016) than for Late-Fall/Fall comparisons (0.001 - 0.01), despite inhabiting the same tributary. Spring Feather River Hatchery individuals stand out as having lower pairwise *F*ST-values in comparison to Fall (0.003 - 0.013) and Late-Fall (0.016) tributaries than to other Spring tributaries (0.018 - 0.036). Similarly, for all Spring/Fall tributary comparisons the lowest observed value is for Spring/Fall Feather River Hatchery individuals (0.003). Further details are summarized in Table S2/Figure S2.

### Estimates of effective population size Ne

*N*e estimates for Fall run ranged from 1,125 to 7,432, with the exception of Coleman Hatchery, which was the lowest (*N*e = 615, CI = 594 – 637) and Merced River, which was the highest (11,436 CI = 6,768; 36,804; Table 1). Estimates for spring groups were lower, ranging from 85 (CI = 84 – 86; Mill Creek) to 561 (CI = 529 – 597; Butte Creek), while Late-Fall had an *N*e = 3,114 (CI = 2,398 – 4,435) and Winter run *Ne=*174 (CI = 171 – 177, Table 1). For Fall Deer and Mill Creek and Spring Feather River Hatchery estimates were negative. This occurs when the LD attributed to sampling variation is larger than the LD attributed to drift (i.e. samples sizes too low to accurately estimate *Ne)*. Here, the groups with negative estimates are among the lowest sample sizes present in the data set (Table 1).

### Assessment of Genomic Diversity

#### MEASURES OF HETEROZYGOSITY

The mean observed heterozygosity is lowest for Winter run individuals from the Upper Sacramento River (mean = 0.1272), followed by Spring individuals from Butte Creek (mean = 0.1587) and highest for Fall individuals from Coleman and Feather River Hatcheries (0.1713 and 0.1741, respectively (Table S3, Figure 3A). Similarly, mean expected heterozygosity is lowest for Upper Sacramento River winter run individuals (mean = 0.1285), and highest for Fall Feather River Hatchery and Mill Creek individuals (mean = 0.1718 and 0.1714, respectively, Table S4, Figure 3B). For both observed and expected heterozygosity, Spring tributaries exhibit a wider range of distributions compared to Fall tributaries, despite the smaller number of Spring tributaries in the data set (Figure 3A, B). Late-Fall values fall into the range of distributions observed among Fall tributaries (Figure 3A, B).

**Figure 3:**
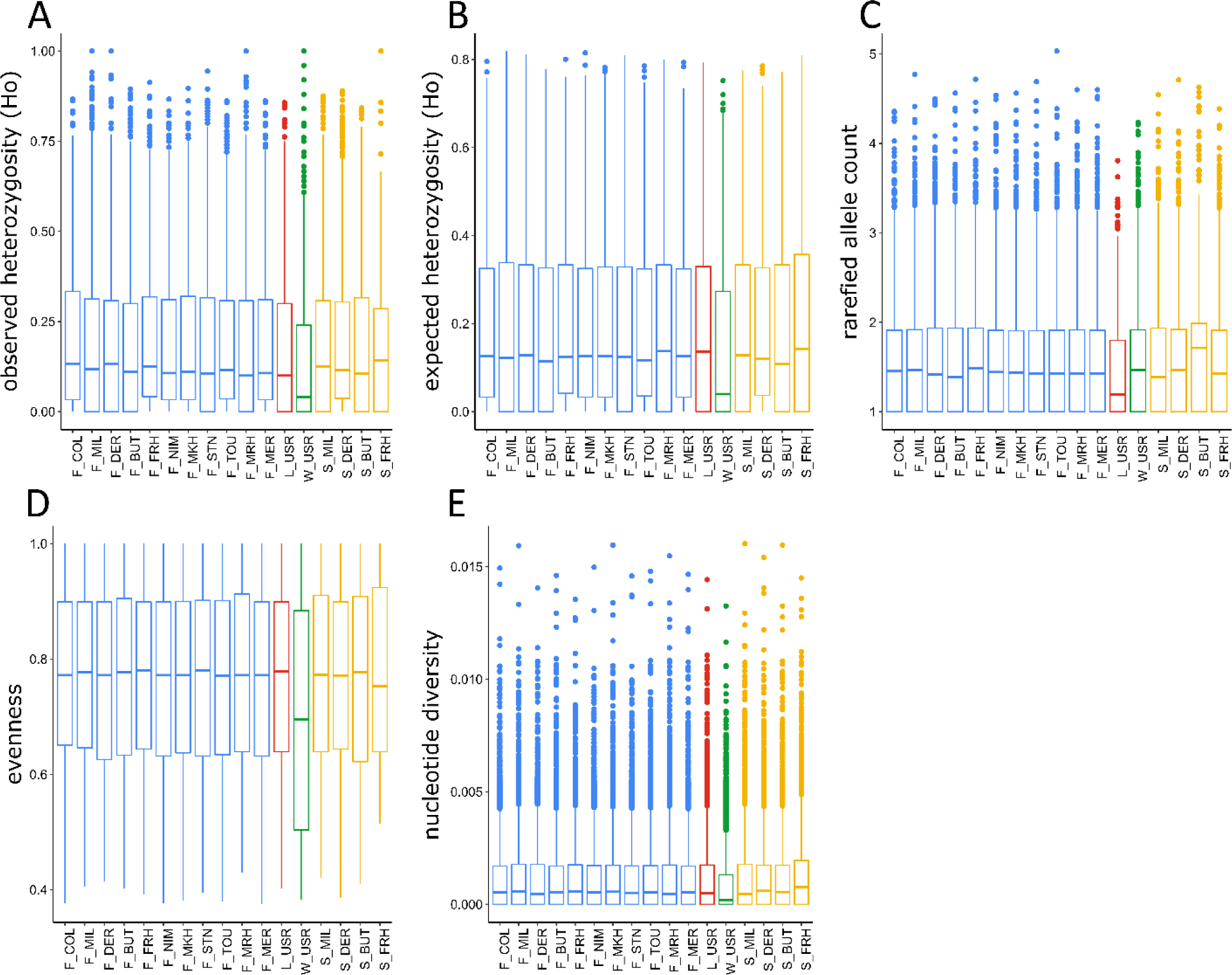
Assessment of the distribution of genomic diversity for individuals grouped by run type within tributaries using heterozygosity-based parameters (A. observed heterozygosity Ho, B. expected heterozygosity Hs), measures of allele diversity (C. Allelic richness, D. Evenness), and sequence-based parameters (E. observed nucleotide diversity). Fall individuals are in blue, Late-Fall in red, Winter depicted in green, and Spring in yellow.

### ALLELIC RICHNESS

The mean values of rarefied allele counts are comparable across tributary/run groups, ranging from 1.51 - 1.52 alleles per locus for all run/tributary groups except Late-Fall Upper Sacramento River (1.36) and Spring Deer Creek individuals (1.48), which exhibit significantly lower mean values (Table S5, Table S6). Despite similar mean values, most pairwise comparisons are significant (Table S5), indicating that even though there is a relatively consistent global number of alleles per locus the patterns of which loci are variable are consistently different across run/tributary groups. For example, Spring Butte Creek individuals exhibit a pattern of rarefied allele counts significantly different from all other locations, and, despite a mean value comparable to most other groups, also exhibit the highest median value (1.71; Figure 3C). Overall, Spring tributaries exhibit more variation among tributaries compared to Fall tributaries (median = 1.38 - 1.71; all pairwise comparisons are significant, Table S5, S6). Distributions are more similar to each other across Fall tributaries, with Butte Creek individuals exhibiting the lowest (1.38) and Feather River Hatchery individuals the highest (1.48) median values (Figure 3C). Notably, individuals from different run types from the same tributary may exhibit quite different patterns. For example, Fall individuals from Mill Creek and the Feather River Hatchery exhibit higher allele counts compared to Spring individuals from the same tributary. By contrast, Fall individuals from Deer and Butte Creek have lower allele counts compared to their Spring counter parts. The strongest contrast is Butte Creek, where Spring individuals exhibit the highest allele counts overall while Fall individuals exhibit the lowest allelic richness among all Fall tributaries (Figure 3C).

#### EVENNESS OF ALLELIC DIVERSITY

Overall, Winter individuals from Upper Sacramento River exhibit significantly lower evenness of allelic richness across loci, indicating that many loci are characterized by rare alleles (Figure 3D). By contrast, Fall Feather River individuals exhibit the highest median evenness (0.78) compared to all other run/tributary groups (Table S7). Within Fall tributaries, mean levels of evenness are all approx. 0.76 though there are some significant differences in their overall distributions (Figure 3D). Again, Spring tributaries exhibit a wider range of mean and median values for evenness compared to Fall (Figure 3D).

#### NUCLEOTIDE DIVERSITY

Winter Upper Sacramento River and Spring Butte Creek individuals exhibit lower nucleotide diversity compared to all other locations (Table S8, Figure 3E). By contrast, Fall Feather River Hatchery individuals exhibit the highest nucleotide diversity (Table S8, Figure 3E).

#### FIXED LOCI

Late-Fall Upper Sacramento individuals and Spring Butte Creek individuals exhibit the highest number of fixed loci (6,416 and 6,210, respectively); for all other run/tributary groups 3,500 - 5,000 fixed loci were identified (Figure 4A). The number of fixed loci in Spring tributaries is generally larger compared to Fall tributaries with the exception of Spring Feather River. By far the two largest intersects are loci fixed exclusively in a single location, Late-Fall Upper Sacramento individuals (460) and Spring Butte Creek individuals (317). All other intersects are < 115 loci. Apart from Spring Feather River hatchery individuals (27), Spring tributaries have more loci fixed in a given tributary (74 - 317) compared to Fall tributaries where 10 - 61 loci are fixed among individuals from a single tributary. Notably, among intersects of loci fixed in two locations, the three largest intersects are all a combination of Late-Fall Upper Sacramento River and a wild Spring population (42 – 115 loci). Overall, about a third of intersects of two run/tributary combinations are loci fixed among Late-Fall Upper Sacramento River individuals and a second location. There is no observed pattern of loci more likely to be fixed among tributaries in geographic proximity (Figure 4A).

**Figure 4:**
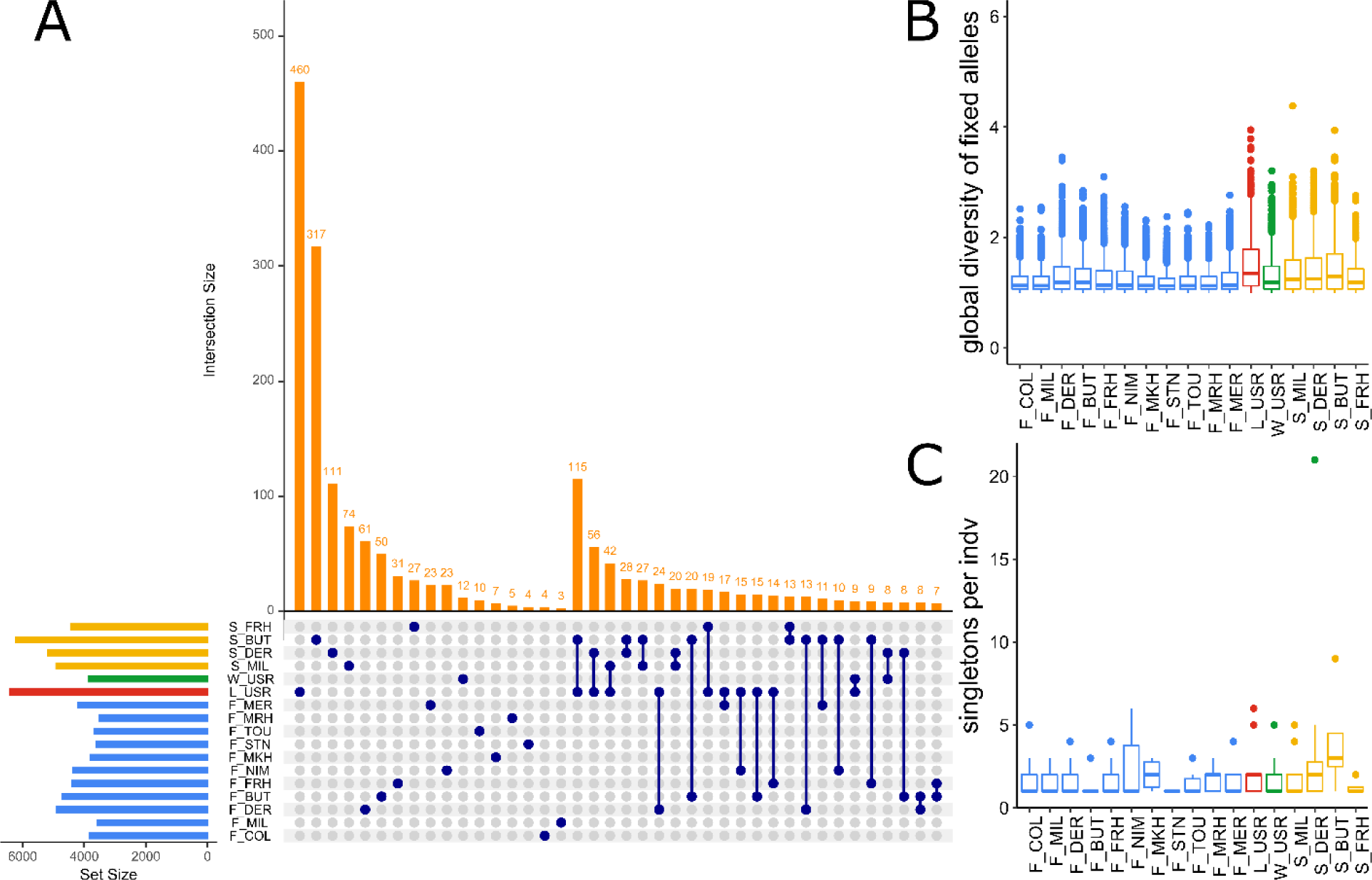
Assessment of fixed loci and singletons. A. Comparisons of fixed loci across run/tributary groups. The set size (horizontal bars) indicate the total number of fixed loci in a given group, the intersect size (vertical orange bars) the number of loci fixed in a single group (single blue dot) or in two (blue dots connected by line). B. Distribution of global allelic richness of loci fixed in a given group. C. Distribution of the number of singletons per individual for each run/tributary group. Fall individuals are in blue, Late- Fall in red, Winter depicted in green, and Spring in yellow.

Loci fixed in Late-Fall Upper Sacramento and Spring Butte Creek individuals also exhibit the highest global allele diversity (mean = 1.46 and 1.41, respectively; Table S9), i.e., loci that are fixed in these groups are more variable when alleles are tabulated across individuals from all runs/tributaries (Figure 4B). By contrast, the global diversity of fixed alleles is lowest in Fall tributary groups, and overall levels are more similar across tributaries (median = 1.13-1.14 with the exception of Fall Deer Creek and Fall Butte Creek at 1.19) compared to spring tributaries where those distributions of global diversity are higher and more variable (median = 1.28 - 1.41; Table S9). The distribution of global diversity among Spring and Fall tributaries varies with some run/tributaries exhibiting much tighter ranges than others (Figure 4B). In general, the proportion of loci that are fixed for a run/tributary group is consistent across

Three hundred forty-seven (1.2% of the total) loci exhibit at least one singleton. A comparison of individuals grouped by run/tributary demonstrates that Spring Butte Creek and Deer Creek individuals (mean = 4.0 and 3.0, respectively) and Late-Fall Upper Sacramento River individuals (mean = 2.23), exhibit the highest mean number of singletons per individual. In general, Fall tributaries exhibit a lower mean number of singletons per individual compared to other run/tributary groups, indicating that they are comparatively less characterized by rare alleles (Figure 4C; Table S12).

#### PRIVATE POLYMORPHISMS

Winter run individuals from Upper Sacramento River have the highest number of loci (N = 153) with private polymorphisms; all other groups have < 100 loci with private polymorphisms (Figure 5). Spring individuals from Butte Creek exhibit the lowest number of loci exclusively polymorphic among individuals of a group (N = 18). Across Late-Fall individuals, 70 loci with private polymorphisms were identified. This number is higher than observed for eight of eleven Fall groups. Notably, individuals from hatcheries fall along the entire range of private polymorphisms; Fall Nimbus Hatchery individuals are on the low end (N = 33) and Spring Feather River Hatchery individuals on the high end (N = 84). Comparing individuals (Figure 5, Table S13). While there are chromosomes with significantly more/less than expected numbers of loci with private polymorphisms no consistent non-random patterns stand out (Table S14). Of the six run/tributary groups with the highest number of chromosomes exhibiting less than the expected number of loci with private polymorphisms, four are Fall run hatcheries (Table S15).

**Figure 5:**
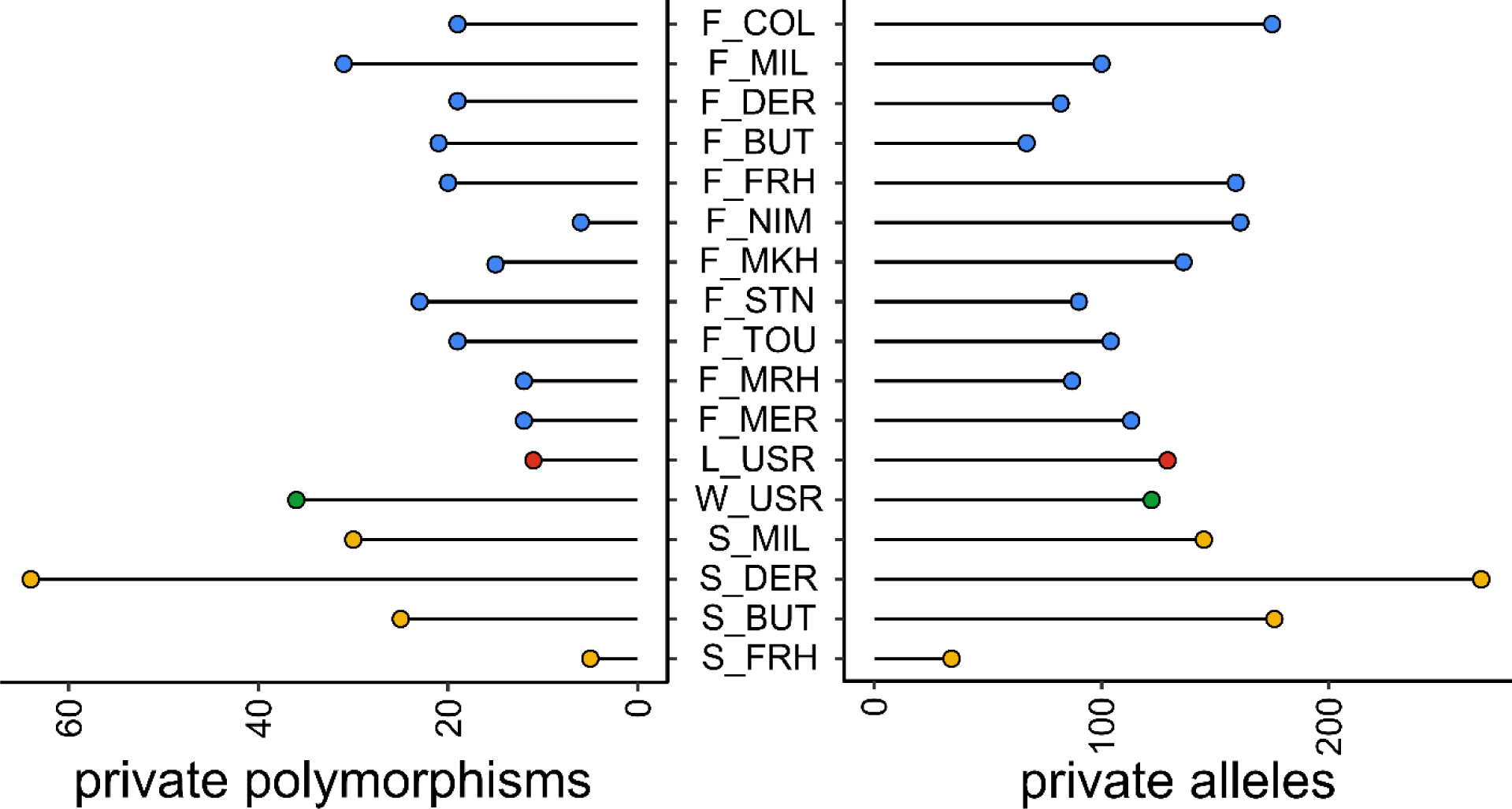
Assessment of unique genomic diversity. Left axis indicates the number of private polymorphisms (loci only variable in a single group), right axis indicates the number of private alleles (alleles found only in a single group) for individuals grouped by run and tributary. Fall individuals are in blue, Late-Fall in red, Winter depicted in green, and Spring in yellow.

#### PRIVATE ALLELES

While most run/tributary groups exhibit 80 - 180 private alleles, Spring Deer Creek individuals exhibit the highest number of private alleles (267) while Fall Butte Creek and Spring Feather River Hatchery individuals exhibit the lowest number of private alleles (34; Figure 5). There is no distinct pattern of hatchery individuals exhibiting more/less private alleles compared to wild individuals, or different run types having consistently higher/lower number of private alleles compared to others; though all groups do exhibit private alleles (Figure 5). Most commonly, private alleles are only carried in 1 - 3 individuals. Winter Upper Sacramento individuals exhibit the highest mean number of individuals carrying private alleles (N = 4.07) compared to other run/tributary groups, followed by Spring Butte Creek individuals (N = 1.34), for all other groups private alleles are found in a mean of 1.01 - 1.17 individuals. Similarly, these two groups have the most “common” private alleles, carried in 25 and ten individuals, respectively (Table S16). Comparing the chromosomal positions of loci with private alleles in

## Discussion

The patterns of genetic diversity observed in populations are the product of past evolutionary and ecological processes. Intra-specific diversity determines the standing variation upon which evolutionary forces act, and levels of phenotypic and genetic variation determine the ability of a population to respond to changes in environmental conditions, the resilience and persistence of a species, and the stability of the ecosystems they inhabit (Prieto et al. 2015; Siefert et al. 2015). Therefore, assessing the biocomplexity of a population complex can help predict the range of possible responses to changing conditions.

Overlooking the importance of biocomplexity at a genetic and phenotypic level and failing to preserve the adaptive potential of populations could have irreversible consequences for the health and sustainability of populations. Here, we present a first ever fine-scale assessment of the biocomplexity contained at the genomic level in California’s Central Valley Chinook salmon population complex. We leverage advancements for genomic analysis to increase our understanding of the hidden diversity contained within a species that is becoming increasingly threatened and at risk of extinction. During this in-depth assessment, we found significant differentiation among and within run types and the tributaries they inhabit, effective population sizes severely below suggested critical thresholds to avoid erosion of genetic diversity and loss in fitness, and corresponding low levels of genetic diversity in populations that have experienced recent demographic declines. Surprisingly, despite apparent gene flow among individuals of the same run type across tributaries, we found each run/tributary group was indeed characterized by a distinct component of unique genomic diversity. This diversity is very likely important to the overall genetic health of the populations, and population complex as a whole, and vital to consider in conservation efforts. Overall, our results emphasize the importance of not only maintaining life history (phenotypic) diversity within and among groups, but also maintaining the genetic diversity of each run and tributary to enhance the portfolio effect, maintain adaptive potential, and ensure the long-term persistence of Chinook salmon in the Central Valley.

Populations become increasingly vulnerable to environmental, demographic, and genetic stochastic effects as their size decreases. Thus, monitoring *N*e of individual components of a population complex is critical for identifying differences in the vulnerability of different groups to genetic issues, evaluating population viability, and predicting evolutionary trajectories to guide conservation managers in their decision-making. Here, individuals from the early migrating run types (Winter, Spring) for each tributary exhibit the lowest *N*e values, well below recommended targets of *N*e = 1,000 to maintain adaptive potential (Frankham et al. 2014). Additionally, we record the lowest estimate of effective population size for the Winter run (Ne = 174) to date. These results further highlight the extreme and alarming risk of extinction facing Winter run.

Similarly, we estimate dangerously low effective population sizes within Spring run populations, with Mill and Deer Creek values being similar to those of Winter run (Ne = 85 and 188, respectively) and Butte Creek being just above 500. There is fine-scale structuring within Spring run. Among wild populations Butte Creek is significantly distinct from Mill and Deer Creeks, which are geographically closer and environmentally more similar to each other. Further, all three wild Spring tributaries are significantly distinct from Feather River Hatchery Spring run. Notably, Spring Feather River Hatchery individuals group more closely with Fall individuals from Coleman and Feather River Hatcheries, pointing towards introgression between runs due to hatchery practices (California-HSRG 2012).

Contrary to the current practice of managing Late-Fall and Fall runs as a single ESU, both results presented here using multi-allelic haplotyped loci and the initial assessment of the sequence data using biallelic SNPs (Meek et al. 2019b) add to the increasing evidence that Late-Fall is genetically distinct from Fall. While our study shows evidence of some distinctions between Fall individuals from more northern tributaries, the differentiation is weak, indicating a loss of biocomplexity with increasing homogenization of Fall run individuals (Williamson & May 2005). In contrast to the early migrating runs (Spring and Winter), *N*e is robust in the late-migrating Fall and Late-Fall groups and at levels sufficient to maintain adaptive potential (Allendorf et al. 2010). The exception to this is Coleman Hatchery Fall individuals, which had an *N*e an order of magnitude lower than other Fall run populations. Also, of note, the *N*e from the Merced River Hatchery Fall run population was also orders of magnitude lower than the *N*e for the putatively wild fish caught in the Merced River. Both of these populations are concerning from a conservation perspective, as maintaining sufficiently large *N*e is a major concern for stocking programs, especially when introgression with wild populations is likely (Ryman & Laikre 1991).

The erosion of genetic diversity due to both genetic drift and inbreeding are inversely proportional to *N*e. Both processes lead to a decreased level of heterozygosity as the rate of alleles being lost due to drift increases and individuals become increasingly likely to mate with individuals with similar genotypes.

Accordingly, if drift is the primary force shaping the genetic diversity within the declining early migrating populations, we would expect to see low levels of heterozygosity and comparatively higher numbers of fixed alleles. Indeed, Upper Sacramento Winter run individuals exhibit the lowest levels of heterozygosity and other measures of genetic diversity compared to all other groups, and the highest number of fixed loci is found in the Spring run groups. Notably, among Spring run groups we find much wider distributions across all measures of diversity – this underscores the stochasticity of genetic drift. Not only is the effect stronger in smaller populations (resulting in an accelerated loss of diversity), but how each component is affected can differ. Thus, groups may diverge from each other by chance alone. In addition, each tributary experiences a different selection regime, driven by environmental differences, resulting in an increased impact of decoupled demographic and environmental stochastic events affecting each population.

The assessment of private polymorphisms (loci only variable in a single group) and private alleles (alleles only found in a single group) reveals that within each tributary, each run exhibits unique components of genetic diversity. Notably, despite Winter run individuals from the Upper Sacramento River having the lowest level of diversity when comparing measures related to heterozygosity and allelic diversity, they exhibit the highest level of private polymorphisms, though they exhibit less diversity. The diversity that is present is unique compared to all the other runs. This is different from Butte Creek Spring individuals which exhibit levels of heterozygosity and allelic diversity similar to Winter run individuals, but also harbor a low number of private polymorphisms, indicating that there are differences in the demographic and evolutionary forces that have shaped genetic diversity in these groups. As a result, despite Spring and Winter run types sharing a similar early migration phenotype and evolutionary history at the GREB1L locus, the genome-wide intra-specific diversity is unique within each group, suggesting differences in standing variation for selective pressures to act upon. Additionally, our new analysis reveals clear, important distinctions in the unique diversity harbored by Late-Fall individuals from the Upper Sacramento compared to Fall run groups, despite Fall and Late-Fall individuals being managed as a single ESU and sharing GREB1L genotypes. The Late-Fall population has lower overall allele counts compared to the Fall populations and the number of private polymorphisms is higher for the Late-Fall compared to almost all (8/11) of the Fall run groups, indicating differences in processes shaping these run types and their evolutionary trajectories.

Finally, while all groups exhibit 80 – 180 private alleles, surprisingly it is groups other than Fall run that tend to be characterized by rare alleles. Since rare alleles are expected to be lost first during bottlenecks, the expectation would be that Fall groups (which have relatively large *Ne*) would instead have a larger number of rare alleles than the groups with smaller *Ne* if genetic drift were the main process affecting allele frequencies. Other processes that could produce these patterns include increased homogenization resulting in an exchange of alleles among tributaries, historical hatchery practices, and/or selection. By contrast, spatially and temporally heterogeneous environments promote and maintain polymorphisms and high levels of standing genetic variation (Svardal et al. 2015; Gulisija & Kim 2015; Bertram & Masel 2019). Indeed, Late-Fall and Spring Deer Creek individuals exhibit the highest numbers of singletons, and Winter and Spring Butte Creek individuals exhibit the highest mean number of individuals carrying private alleles. Even Spring Feather River Hatchery individuals, which have introgressed with Fall individuals in the past, carry private polymorphisms. Overall, these patterns support the conclusion that processes such as balancing selection are occurring to maintain diversity, despite declining population sizes.

This presence of unique diversity among and within individual components of the Central Valley population complex underscores the importance of a management strategy that seeks to maintain a robust portfolio at both a phenotypic and genotypic level. While it is important to acknowledge that the (neutral) genetic diversity of a population is not always correlated with functional diversity (Reed & Frankham 2001), the variation of genotypes within and among individuals has been demonstrated to be a suitable proxy to predict fitness of individuals and the ability of populations and ecosystems to respond to changes in environmental conditions (Vazquez-Dominguez et al. 1999; Reed & Frankham 2003; Reusch et al. 2005; Hoffman et al. 2014). Furthermore, examples from translocation and genetic rescue efforts have demonstrated that heterozygosity and genetic diversity can be more efficient predictors of success than the ability to match (neutral) genotypes as closely as possible to individuals already present in the population (Coleman et al. 2013; Scott et al. 2020). Thus, while the locus that affects run timing is undoubtedly important and of critical conservation concern (Ford et al. 2020), the evolution and accumulation of additional diversity facilitated by differences in run-timing in specific habitats may also be of critical importance for population success (Allendorf et al. 2010). Losing early-run populations therefore runs the risk of losing both early-run alleles (i.e., the ability to recover the early-run phenotype) and the more cryptic yet likely important unique components of genetic diversity harbored among and within migration phenotypes.

This loss of diversity and increasing genetic homogenization may be a more important factor driving the loss of the portfolio than demographic synchronization itself (Dedrick & Baskett 2018; Des Roches et al. 2021). Because of their complicated life history, environmental pressures differ widely across salmon life stages such that the genotypes and phenotypes that confer higher survival probability at one life stage do not necessarily translate into the genotypes and phenotypes best matching conditions during a different life stage. Additionally, climate change will impact environmental conditions in individual tributaries differently, again necessitating genomic diversity across the Central Valley to allow adaptation to changing conditions (Yates et al. 2008). Important phenotypic traits, including growth, temperature tolerance, and stress responses, are likely polygenic traits, controlled by many loci of small effects, and populations characterized by the presence of a large proportion of polygenic phenotypes are more likely to adapt to new conditions and therefore increase population viability with rapidly changing and fluctuating environmental conditions (Kardos & Luikart 2019).

Provided sufficient standing genetic diversity exists and is preserved in the Central Valley, the intraspecific variation present in Central Valley Chinook may allow adaption to changing conditions (Hairston et al. 2005; Richardson et al. 2014; Messer et al. 2016). Even though anthropogenic impacts significantly alter the composition and structure of both neutral and functional diversity at a genetic level, the conservation of intraspecific genetic diversity is frequently overlooked (Laikre et al. 2010; Des Roches et al. 2021) despite serving as the fundamental building block of biodiversity. Indeed, finding unique diversity within each run/tributary group comprising the population complex of Chinook salmon in California’s Central Valley underscores the importance of monitoring intraspecific genomic diversity at multiple levels (across and within locations and life history phenotypes) to inform conservation and management policies that counteract genetic homogenization and conserve biocomplexity at a genomic level. Additionally, our results highlight the necessity of managing population complexes as a whole, with a focus on maintaining biocomplexity at multiple scales, as an important factor determining the resilience to changing environmental pressures.

## Acknowledgements

The authors would like to thank the members of the Meek Lab for helpful discussions and Melinda Baerwald for helpful feedback on the manuscript. Additionally, we would like to thank Pascale Goertler for important discussions related to the project.

## Data Availability

Data for this study are available from Dryad (https://doi.org/10.5061/dryad.tht76hdvt). Supplementary Material 1 and 2 contain fully reproducible code supporting the analysis; research compendium available at https://github.com/sjoleary/ONC_GenDiv.

## Supplementary material

*Note: supplementary files will not render properly during conversion to pdf for review – html files that can be opened in any browser can be downloaded from the research compendium at* https://github.com/sjoleary/ONC_GenDiv.

Supplementary Material 1 (OLeary_SupMat_Genotyping.html): Data acquisition and processing (Genotyping).

Supplementary Material 2 (OLeary_SupMat_Genotyping.html): Standalone extended methods and results describing the assessment of population differentiation and patterns of genomic diversity. Contains R code used for data analysis and supplementary figures and tables cited in the manuscript.

