## Supplementary material for "Every cog and wheel: Identifying biocomplexity at the genomic and phenotypic level in a population complex of Chinook salmon": Supplementary Material 1.html (from another version of this preprint: version 1, 2021-03-28)

Data acquisition and processing: Genotyping

### Map reads to reference genome

#### Reference genome

Downloaded from NCBI (https://www.ncbi.nlm.nih.gov/assembly/GCF\_002872995.1/) (Christensen et al. ⊕2018Christensen, Kris A., Jong S. Leong, Dionne Sakhrani, Carlo A. Biagi, David R. Minkley, Ruth E. Withler, Eric B. Rondeau, Ben F. Koop, and Robert H. Devlin. 2018. “Chinook salmon (Oncorhynchus tshawytscha) genome and transcriptome.” Edited by James P. Meador. *PLoS ONE* 13 (4). Public Library of Science: e0195461. doi:10.1371/journal.pone.0195461.), consisting of 34 chromosomes and 13,995 unplaced sequences.

Unzip and rename (needs to be indexed etc. for read mapping and SNP calling).

```
$HOME/CHINOOK/data/REF/CHRIST2018

# unzip genome file
gunzip GCF_002872995.1_Otsh_v1.0_genomic.fna.gz

# rename
mv GCF_002872995.1_Otsh_v1.0_genomic.fna Otsh_v1.0.fasta
```

**Genome assembly stats**:

- Total sequence length: 2,425,713,975
- Total ungapped length: 2,359,046,630
- Gaps between scaffolds: 1,916
- Number of scaffolds: 15,946
- Scaffold N50: 1,728,323
- Scaffold L50: 336
- Number of contigs: 69,485
- Contig N50: 133,169
- Contig L50: 4,051
- Total number of chromosomes and plasmids: 35
- Number of component sequences (WGS or clone): 15,946

#### Map reads

Any renaming of files needs to happen before `fastq`-files are mapped. The population designation (before underscore) is used by `freebayes` to call SNPs so it is important that population designation make biological sense. Here, all individuals are being grouped into a single group `ONC_`.

Transfer copy of `reference.fasta` (rename genome `fasta` file) and associated files into library directories with demultiplexed and quality trimmed sequences, then run `dDocent` from within each `data/SEQ` directory to map reads to reference genome (`map.slurm`).

`dDocent` wraps `BWA` (Li and Durbin ⊕2009Li, Heng, and Richard Durbin. 2009. “Fast and accurate short read alignment with Burrows-Wheeler transform.” *Bioinformatics* 25 (14): 1754–60. doi:10.1093/bioinformatics/btp324.) to map reads; parameters used: `bwa mem -L 20,5 -t 35 -a -M -T 10 -A 1 -B 9 -O 9 -R`; `dDocent` filters reads that are unmapped or ambiguously mapped (MAPQ > 1).

```
# navigate to starting directory
cd $HOME/CHINOOK/data/SEQ/

for i in BMAG*; do

    cd $HOME/CHINOOK/data/SEQ/$i

    ## PREP FILES ----

    # copy of configuration file
    cp $HOME/CHINOOK/scr/map.config .

    # copy of reference and associated files
    cp $HOME/CHINOOK/data/REF/CHRIST2018/reference.* .

    # make sure path correct to find conda
    export PATH=$PATH:$HOME/miniconda3/bin

    ## RUN dDOCENT ----

    # open bioconda environment
    source activate $HOME/miniconda3/envs/ddocent_env

        # run dDocent to quality trim & map reads to reference genome
        dDocent map.config

    # close bioconda environment
    source deactivates

    rm $i/cat-RRG.bam*

done
```

To run on HPCC need to use `SLURM` scripts to queue job, load modules, and activate `dDocent Bioconda environment` which contains all the necessary programs/software for the SNP calling pipeline. Move `slurm` output to scratch folder to be able to refer to run length/resources needed; remove unnecessary intermediate files.

Remove reads that map with mapping quality < 5, create a `bamlist.list` file containing all names (paths) for each bam file, then concatenate all filtered `bam`-files into single `bam`-file and use `bedtools` to query coverage per interval.

```
# navigate to project directory
cd $HOME/CHINOOK/

# load modules
module purge
module load GCC/6.4.0-2.28 OpenMPI/2.1.2
module load SAMtools/1.9

# create concatenated bam file
samtools merge --threads 40 -b data/SEQ/bamlist.list -f data/SNP_CALLING/cat-RRG.bam
samtools index data/SNP_CALLING/cat-RRG.bam

# get coverage of concatenated file
module purge
module load GCC/4.9.3-2.25  OpenMPI/1.10.2
module load BEDTools/2.26.0

bedtools genomecov -ibam data/SNP_CALLING/cat-RRG.bam -bga > results/all_lib_filtered.coverage
```

#### Compare mapping intervals to identify putative loci

Removing reads with low mapping quality and intervals with low coverage will increase computational efficiency by removing extraneous reads and regions that are not consistently recovered.

Table 1a: Number of features (intervals) for each coverage bin.

| cov\_bin | n |
| --- | --- |
| >50 | 7104978 |
| <50 | 30474257 |
| 0 | 5955562 |
| Retain onl | y intervals with coverage > 50. |

During coverage extraction intervals can be split up because reads do not stack up cleanly, i.e. mapping intervals will split into multiple adjacent intervals. Use `bedtools` to merge adjacent intervals (up to 15bp gap).

```
# load modules
module purge
module load GCC/4.9.3-2.25  OpenMPI/1.10.2
module load BEDTools/2.26.0

# go to project directory
cd $HOME/CHINOOK

# merge adjacent intervals
bedtools merge -d 15 -i results/concat_bam_intervals.bed > results/merged_mapping_intervals.bed
```

Compare depth distribution for merged mapping intervals.

Fig 1: Distribution of mapping interval length for merged intervals. Red dashed line indicates 300bp length fragments. Filtered reads are expected to be between 75-100bp in length.

Table 1b: Number of mapping fragments with length > 500bp

| length > 500 | n |
| --- | --- |
| TRUE | 88 |

Table 1c: Number of mapping fragments with length > 25bp

| length > 25 | n |
| --- | --- |
| TRUE | 169485 |

Retain mapping intervals between 25 and 500 bp length; name contigs as `Contig_1` … Contig\_169485.

#### Extract contigs with reads mapped & map reads

For downstream applications like haplotyping it is necessary to have individual contigs that reads have mapped to and SNPs have been called on as opposed to the entire genome (with contiguous scaffolds per chromosome). Extracting “loci” (contigs) that are relevant needs to occur at this stage; this will also increase computational efficiency during mapping/SNP calling. Use identified mapping intervals to create a `reference.fasta` file that contains sequences equivalent to the intervals that have reads mapped to them.

```
# extract intervals with SNPs
module purge
module load GCCcore/6.4.0
module load BEDTools/2.27.1

# extract intervals in filtered bed file
bedtools getfasta -name -fi data/REF/CHRIST2018/Otsh_v1.0.fasta -bed data/REF/CHRIST2018/map_intervals.bed -fo data/REF/CHRIST2018/reference.fasta
```

Copy new `reference.fasta` into each sequencing folder and map all the `fastq`-files.

```
# loop over sequencing files
for i in $HOME/CHINOOK/data/SEQ/BMAG*; do

    echo mapping $i ... `date`

    cd $i

        rm cat-RRG.bam*
        rm reference.fasta*
        rm namelist
        rm dDocent.runs
        rm bamlist.list
        rm -r logfiles
        rm mapped.bed

    ## PREP FILES ----

    # copy of configuration file
    cp $HOME/CHINOOK/scr/map.config .

    # copy of reference and associated files
    cp $HOME/CHINOOK/data/REF/CHRIST2018/reference.* .

    ## RUN dDOCENT ----

    # make sure path correct to find conda
    export PATH=$PATH:$HOME/miniconda3/bin

    # open bioconda environment
    source activate $HOME/miniconda3/envs/ddocent_env

        # run dDocent to quality trim & map reads to reference genome
        dDocent map.config

    # close bioconda environment
    source deactivate

    rm $i/cat-RRG.bam*

done
```

#### QA/QC read mapping

**Query reads counts and mapping quality**

During the mapping stage, `dDocent` calls `BWA` to map reads from the individuals in the folder to reference and create a `-RG.bam`-file (retaining only unambiuously mapped reads) and `-RG.bam.bai` (index file) for each individual. The second column of a BAM file contains FLAGs with binary encoded information on mapping quality, pairedness etc. that can be used to compare the mapping efficiency to the reduced representation reference.

Determine the number of reads in each `fastq` file (trimmed/untrimmed) and the number of mapped reads in the corresponding bam files, then filter reads with mapping quality < 5. Determine the coverage for mapping intervals for the un/trimmed `*.bam`-files.

```
# navigate to project directory
cd $HOME/CHINOOK

# create results file
echo "SAMPLE TOTAL_UNTRIMMED TOTAL_TRIMMED MAPPED MAPQ5 MAPQ10 MAPQ60" > $HOME/CHINOOK/results/BMAG_mapping.comp

for d in data/SEQ/BMAG*; do

    # navigate to directory
    cd $d

    # keep track of directory being processed
    echo $d -----

    # loop over bam files in directory
    while read i; do

        # keep track of file being processed
        echo $i

        # MAPPING QUALITY STATS ----

        # load modules
        module purge
        module load GCC/6.4.0-2.28 OpenMPI/2.1.2
        module load SAMtools/1.9

        # untrimmed reads
        TOT_UNTR=$(zcat $i.F.fq.gz | echo $((`wc -l`/4)))

        # trimmed reads
        TOT_TRIM=$(zcat $i.R1.fq.gz | echo $((`wc -l`/4)))

        # mapped reads
        MAP=$(samtools view $i-RG.bam -F 4 -c)

        # reads with MAPQ>5
        MAPQ5=$(samtools view $i-RG.bam -q 5 -c)

        # reads with MAPQ>10
        MAPQ10=$(samtools view $i-RG.bam -q 10 -c)

        # reads with MAPQ=60
        MAPQ60=$(samtools view $i-RG.bam -q 60 -c)

        # add to file
        echo "$i $TOT_UNTR $TOT_TRIM $MAP $MAPQ5 $MAPQ10 $MAPQ60" >> $HOME/CHINOOK/results/BMAG_mapping.comp

        # filter to remove reads with MAPQ < 5
        samtools view -bSq 5 $i-RG.bam > $HOME/CHINOOK/data/SNP_CALLING/$i.-RG.bam

    # dDocent creates file with sample ids
    done < namelist

    # return to project directory
    cd $HOME/CHINOOK/

done
```

**Assess proportion of reads mapped per individual**

Fig 2: Distribution of proportion of reads in fastq file mapped per individual (grouped by library). Red dashed line indicates mean proportion of reads mapped overall.

After quality trimming, individuals (N = 561) have a mean number of 2895547 +/- 2450681 (min = 138258, max = 22337503) quality trimmed sequenced reads. On average, 62 +/- 12 % of reads are mapped to the genome.

**Assess mapping quality**

Fig 3: Distribution of proportion of reads quality trimmed, and proportion of quality trimmed reads mapped per individual overall and for varying threshold values of mapping quality. MAPQ ranges from 1 (ambiguously mapped) to 60; dDocent removes unmapped/ambiguously mapped reads from bam files.

### SNP calling

Transfer copy of reference genome as `reference.fasta` along with symlinks for all `fq` and `bam`-files into the `SNP_Calling` folder to call variants across all individuals using `freebayes` (Garrison and Marth ⊕2012Garrison, Erik, and Gabor Marth. 2012. “Haplotype-based variant detection from short-read sequencing.” *Plos One* 11 (3): e0151651. doi:arXiv:1207.3907 [q-bio.GN].). `freebayes` is a bayesian haplotype-based variant detecter and calls variants based on alignments, i.e. based on the sequence of (short) reads aligned to a reference to determine the most-likely combination of genotypes for a set of individuals at each postion in the alignment to a reference.

Execute `dDocent` from within `SNP_Calling`-folder to call variants across all individuals - need to submit `SLURM`-script to run on MSU’s `hpcc`.

File `TotalrawSNPs.vcf` contains all raw SNP/INDEL calls. Do not need to keep links of `fq.gz`-, `bam`-, `.bam.bai`-files after SNPs have been called. Copy `TotalRaSNPs.vcf` to `VCF`-directory for SNP filtering.

### SNP filtering

#### Raw data

**Assess individuals & populations sampled**

Generate a list of all individuals included in the VCF-file to be filtered.

Use `raw.ind` file to write text files of individuals in each library and grouped by runs/locations.

Table 2a: Number of samples per location and run type.

| LOCATION | F | L | S | W |
| --- | --- | --- | --- | --- |
| BTC | 2 | 2 | NA | NA |
| BUT | 29 | 2 | 32 | NA |
| COL | 30 | 2 | NA | NA |
| DER | 31 | NA | 46 | NA |
| FRH | 29 | NA | 30 | NA |
| MER | 34 | NA | NA | NA |
| MIL | 35 | NA | 40 | NA |
| MKH | 30 | NA | NA | NA |
| MRH | 29 | NA | NA | NA |
| NIM | 30 | NA | NA | NA |
| STN | 30 | NA | NA | NA |
| TOU | 35 | NA | NA | NA |
| USR | 2 | 35 | NA | 29 |

Table 2b: Number of samples library.

| LIB | n |
| --- | --- |
| BMAG001 | 31 |
| BMAG002 | 33 |
| BMAG006 | 27 |
| BMAG007 | 34 |
| BMAG008 | 32 |
| BMAG009 | 35 |
| BMAG010 | 35 |
| BMAG013 | 35 |
| BMAG017 | 36 |
| BMAG020 | 45 |
| BMAG021 | 45 |
| BMAG022 | 45 |
| BMAG023 | 46 |
| BMAG024 | 43 |
| BMAG025 | 42 |

**Compare Individual & SNP stats**

Use `VCFtools` to create stats files for depth, missing data, heterozygosity, and site quality for the raw data.

```
vcftools --vcf data/SNP_CALLING/TotalRawSNPs.vcf --out data/VCF/ONC_raw --depth
vcftools --vcf data/SNP_CALLING/TotalRawSNPs.vcf --out data/VCF/ONC_raw --site-mean-depth
vcftools --vcf data/SNP_CALLING/TotalRawSNPs.vcf --out data/VCF/ONC_raw --site-quality
vcftools --vcf data/SNP_CALLING/TotalRawSNPs.vcf --out data/VCF/ONC_raw --missing-indv
vcftools --vcf data/SNP_CALLING/TotalRawSNPs.vcf --out data/VCF/ONC_raw --missing-site
vcftools --vcf data/SNP_CALLING/TotalRawSNPs.vcf --out data/VCF/ONC_raw --het
```

Compare patterns of missing data, read depth, and quality of SNP calls across individuals and loci.

Fig 4: Distribution and relationships of missing data, read depth, and SNP quality across loci and individuals.

The raw data set contains 574 individuals and 2004505 loci.

#### Choose threshold values for quality score, coverage, missing data, and mapping/variant calling artifacts

**FILTER 0: Decompose indels to retain only SNPs**

Identify low quality individuals to remove from the data set; defined as individuals with a mean coverage of < 3 reads across all loci, > 75% missing data (thresholds based on initial exploratory filtering to maximize number of indv/loci retained in data set). In addition, remove all individuals from library BMAG002, BMAG008, and BMAG009 due to a library effect.

A total of 118 were flagged as low quality based on missing data, depth, or heterozygosity.

Remove LQ individuals, decompose indels, and retain only biallelic SNPs.

Compare data set -

Data set contains 456 individuals and 1738126 loci after removing low quality individuals and decomposing indels set into SNPs.

**FILTER 1: Confidence in SNP call**

The QUAL column of a VCF file is a phred-based score indicating the probability that the variant shown in the ALT column is wrong. Given the Phred quality score (Q), and the probability that a base is incorrectly called (P), Q = -10(Log10P). A quality score of 20 indicates, a 1 in 100 chance that the SNP site has been called incorrectly (i.e. 99% probability that correct call).

Code genotypes with genotype call quality < 20 or genotype depth < 5 as missing and remove loci with quality < 20, mean depth < 10 and/or > 50% missing data.

Import singletons and genotype depth file to create list of loci to exclude based on depth.

Fig 5: Distribution of number of singletons per contig.

Data set contains `nrow(singletons)` singletons of those `nrow(doubletons)` are called as a homozygote in one individual (doubletons). Overall, there are `nrow(contigs_total)` remaining contigs, `nrow(contigs_singletons)` (14.27%) contain singletons.

Fig 6: Distribution of the number of singletons per individual

Compare the genotype depth to ensure that there is sufficient depth to recover both alleles (i.e. are doubletons true homozygotes) and ensure quality of singleton gentoypes.

Fig 7: Distribution genotype depth per singleton/doubleton.

```
##  5% 25% 50% 75% 95% 99%
##   6  10  18  45 112 158
```

Identify low quality singletons (potential artifacts resulting from PCR or sequencing error) by flagging singletons with depth < 10 reads and doubletons with depth < 20 reads.

Table 3a: Number of singletons/doubletons with depth < 10 reads.

| SINGLETON/DOUBLETON | n |
| --- | --- |
| D | 161 |
| S | 8805 |

Table 3b: Number of singletons/doubletons with depth < 20 reads.

| SINGLETON/DOUBLETON | n |
| --- | --- |
| D | 320 |
| S | 17560 |

Filter loci with quality score < 20 and singletons/doubletons with low read depth (<10/20 reads).

```
# filter LQ SNP calls
vcftools --vcf data/VCF/temp/ONC.F1a.recode.vcf --out data/VCF/temp/ONC.F1 --exclude-positions data/VCF/LQ_F0.loci --recode --recode-INFO-all

# query stats
vcftools --vcf data/VCF/temp/ONC.F1.recode.vcf --out data/VCF/ONC.F1 --depth
vcftools --vcf data/VCF/temp/ONC.F1.recode.vcf --out data/VCF/ONC.F1 --site-mean-depth
vcftools --vcf data/VCF/temp/ONC.F1.recode.vcf --out data/VCF/ONC.F1 --missing-indv
vcftools --vcf data/VCF/temp/ONC.F1.recode.vcf --out data/VCF/ONC.F1 --missing-site
vcftools --vcf data/VCF/temp/ONC.F1.recode.vcf --out data/VCF/ONC.F1 --het
```

Compare stats post-filtering.

Data set contains 456 individuals and 103281 SNP sites.

**FILTER 2: Genotype call rate and allowed missing data per indv**

Flag loci that were not called in > 50% of individuals in a given library.

Fig 8: Distribution of missing data per locus for individuals grouped by library

Table 4: Number of SNPs called in < 50% of individuals of a given library.

| LIB | n |
| --- | --- |
| BMAG001 | 738 |
| BMAG006 | 1,039 |
| BMAG007 | 1,024 |
| BMAG010 | 608 |
| BMAG013 | 1,571 |
| BMAG017 | 2,720 |
| BMAG020 | 2,148 |
| BMAG021 | 1,446 |
| BMAG022 | 1,791 |
| BMAG023 | 2,300 |
| BMAG024 | 3,977 |
| BMAG025 | 3,397 |

A total of 8161 loci were called in less than 50% of individuals in one or more libraries. Loci being inconsistently called among libraries can result in library effects.

Flag loci that were not called in > 50% of individuals at a given run type (this will be checked again at the end of the filtering process).

Fig 9: Distribution of missing data per locus for individuals grouped by library.

Table 5: Number of SNPs called in < 50% of individuals of a given run type.

| POP | n |
| --- | --- |
| FALL | 916 |
| LATE-FALL | 344 |
| SPRING | 1,418 |
| WINTER | 2,296 |

A total of 3581 SNPs were called in less than 50% of individuals of a given run type.

Remove loci that were not consistently called across all libraries and run types (8680 SNPs).

```
vcftools --vcf data/VCF/temp/ONC.F1.recode.vcf --out data/VCF/temp/ONC.F2a --exclude-positions data/VCF/LQ_F2.loci --recode --recode-INFO-all

vcftools --vcf data/VCF/temp/ONC.F2a.recode.vcf --out data/VCF/ONC.F2a --missing-indv
```

Identify individuals with > 75% missing data

Fig 10: Missing data per individual.

Remove flagged individuals (N = 19).

```
vcftools --vcf data/VCF/temp/ONC.F2a.recode.vcf --out data/VCF/temp/ONC.F2 --remove data/VCF/F2_LQ.indv --recode --recode-INFO-all

vcftools --vcf data/VCF/temp/ONC.F2.recode.vcf --out data/VCF/ONC.F2 --depth
vcftools --vcf data/VCF/temp/ONC.F2.recode.vcf --out data/VCF/ONC.F2 --site-mean-depth
vcftools --vcf data/VCF/temp/ONC.F2.recode.vcf --out data/VCF/ONC.F2 --missing-indv
vcftools --vcf data/VCF/temp/ONC.F2.recode.vcf --out data/VCF/ONC.F2 --missing-site
vcftools --vcf data/VCF/temp/ONC.F2.recode.vcf --out data/VCF/ONC.F2 --het
```

Compare data set post-filtering -

Data set contains 437 individuals and 94601 SNP loci.

**FILTER 3: Filter loci and individuals based on depth, variance in depth, and genotype call rate**

Determine mean depth and variance per locus per library.

Compare distribution of depth coverage per locus per library. Identify loci that do not have consistent read depth among libraries (can lead to library effects), and/or across individuals.

Fig 11: Distribution of mean depth per locus per library

Identify loci with large variance in mean depth across libraries and/or individuals by calculating the coefficient of variance (STD/MEAN).

Fig 12: Comparison of mean across all individual to mean weighted by library and coefficient of variance of read depth across individuals and between libraries. If loci have consistent coverage across loci the mean read depth per locus across all individuals and weighted by library should fall on the red diagonal.

Remove loci flagged for high variance in depth across all individuals and between libraries (removes library effects, N = 50), remove loci with mean depth < 15 or genotype call rate < 75%.

```
vcftools --vcf data/VCF/temp/ONC.F2.recode.vcf --out data/VCF/temp/ONC.F3a --exclude-positions data/VCF/LQ_F3.loci --min-meanDP 15 --max-missing 0.75 --recode --recode-INFO-all

vcftools --vcf data/VCF/temp/ONC.F3a.recode.vcf --out data/VCF/ONC.F3a --depth
vcftools --vcf data/VCF/temp/ONC.F3a.recode.vcf --out data/VCF/ONC.F3a --geno-depth
```

Compare mean depth and number of sites called per individual.

Fig 13: Comparison of mean depth and number of sites called per individual.

Individuals with higher mean depth should have less missing data (called for higher number of sites). High variance in depth can make an individual appear to have good coverage, but may have many loci with false homozgygous calls due to low genotype depth that is masked by high coverage loci.

Use genotype depth file to identify individuals with high variance in read depth across loci.

Fig 14: Comparison of distribution depth per locus within individuals.

Flag LQ individuals based on depth (mean depth < 5 and median depth = 0).

Remove flagged individuals (N = 19).

```
vcftools --vcf data/VCF/temp/ONC.F3a.recode.vcf --out data/VCF/temp/ONC.F3 --remove data/VCF/F3_LQ.indv --recode --recode-INFO-all

vcftools --vcf data/VCF/temp/ONC.F3.recode.vcf --out data/VCF/ONC.F3 --depth
vcftools --vcf data/VCF/temp/ONC.F3.recode.vcf --out data/VCF/ONC.F3 --site-mean-depth
vcftools --vcf data/VCF/temp/ONC.F3.recode.vcf --out data/VCF/ONC.F3 --missing-indv
vcftools --vcf data/VCF/temp/ONC.F3.recode.vcf --out data/VCF/ONC.F3 --missing-site
vcftools --vcf data/VCF/temp/ONC.F3.recode.vcf --out data/VCF/ONC.F3 --het
vcftools --vcf data/VCF/temp/ONC.F3.recode.vcf --out data/VCF/ONC.F3 --hardy
```

Compare stats post-filtering:

Fig 15: Comparison of levels of missing data, depth, and heterozygosity per individual and locus.

**FILTER 4: Allele balance**

AB: Allele balance at heterozygous sites: a number between 0 and 1 representing the ratio of reads showing the reference allele to all reads, considering only reads from individuals called as heterozygous.

Allele balance is the ratio of reads for reference allele to all reads, considering only reads from individuals called as heterozygous. Values range from 0 - 1; allele balance (for real loci) should be approx. 0.5.

Fig 16: Distribution of allele balance across all loci.

Compare relationship of allele balance, heterozygosity, and mean depth per locus.

Fig 17: Comparison of allele balance, heterozygosity, and mean depth per locus.

Filter contigs with SNP calls with AB > 0.35, AB > 0.75; retain loci very close to 0 (retain loci that are fixed variants). Remove genotypes if the quality sum of the reference or alternate allele was 0.

```
../../../scr/vcflib/bin/vcffilter -s -f "AB > 0.35 & AB < 0.75 | AB < 0.01 | AB > 0.99" -s -g "QR > 0 | QA > 0" ONC.F3.recode.vcf > ONC.F4.vcf

awk '!/#/' ONC.F4.vcf | wc -l
```

Remaining SNPs: 47,974.

**FILTER 5: mapping quality**

```
cut -f8 ONC.F4.vcf | grep -oe "MQM=[0-9]*" | sed -s 's/MQM=//g' > ONC.F5.MQM

cut -f8 ONC.F4.vcf | grep -oe "MQMR=[0-9]*" | sed -s 's/MQMR=//g' > ONC.F5.MQMR
```

Remove loci based on ratio of mapping quality for reference and alternate allele, i.e. sites that have a high discrepancy between the mapping qualities of two alleles.

Fig 18: Comparison of mapping quality for reference and alternate allele of a given locus.

Filter loci with mapping quality ratio < 0.25 and > 1.75.

Remaining SNPs: 45.755.

**FILTER 6: Maximum depth & Quality**

Identify distribution of depth (based on original data set) to identify loci with excess coverage. Original number of individuals in data set is 574 (INFO flags in filtered data set are are based on original number of individuals in data set). Create file with the original site depth and quality score for each site.

Calculate average depth and standard deviation and compare to locus quality.

Fig 19: Depth and quality distribution for all SNP loci. Red loci are loci with low quality scores compared to their high depth.

Mean depth per locus (across all indivuals) is 14498 and the standard deviation is 8722.1. Filter SNP site with depth > mean depth + 1 standard deviation = 31942 and that have quality scores < 2x the depth at that site and output depth per site.

Compare the distribution of mean depth per site averaged across individuals to determine cut-off value of sites with excessively high depth indicative of paralogs/multicopy loci.

Fig 20: Mean depth per locus. Red dashed line indicates the mean depth across all loci, the dashed blue lines indicate the 95th and 99th quartiles.

Table 6: Number of loci with > 60 reads.

| MEAN\_DEPTH > 75 | n |
| --- | --- |
| FALSE | 45719 |
| TRUE | 162 |

Filter loci maximum mean read depth > 75 to eliminate putative paralogs.

Analyze stats post-filtering:

Fig 21: Comparison of levels of missing data and coverage per locus and individual.

Data set contains 45720 SNP sites and 418 individuals.

**FILTER 7: Missing data and minimum depth per locus**

Filter loci with mean read depth < 15 and genotype call rate < 90%.

```
vcftools --vcf data/VCF/temp/ONC.F6.recode.vcf --out data/VCF/temp/ONC.F7 --min-meanDP 15 --max-missing 0.9 --recode --recode-INFO-all

vcftools --vcf data/VCF/temp/ONC.F7.recode.vcf --out data/VCF/ONC.F7 --depth
vcftools --vcf data/VCF/temp/ONC.F7.recode.vcf --out data/VCF/ONC.F7 --site-mean-depth
vcftools --vcf data/VCF/temp/ONC.F7.recode.vcf --out data/VCF/ONC.F7 --missing-indv
vcftools --vcf data/VCF/temp/ONC.F7.recode.vcf --out data/VCF/ONC.F7 --missing-site
vcftools --vcf data/VCF/temp/ONC.F7.recode.vcf --out data/VCF/ONC.F7 --het
vcftools --vcf data/VCF/temp/ONC.F7.recode.vcf --out data/VCF/ONC.F7 --hardy

# library BMAG002
vcftools --vcf data/VCF/temp/ONC.F7.recode.vcf --out data/VCF/temp/BMAG002 --keep data/VCF/BMAG002.ind --recode --recode-INFO-all

vcftools --vcf data/VCF/temp/BMAG002.recode.vcf --out data/VCF/BMAG002 --hardy

# library BMAG006
vcftools --vcf data/VCF/temp/ONC.F7.recode.vcf --out data/VCF/temp/BMAG006 --keep data/VCF/BMAG006.ind --recode --recode-INFO-all

vcftools --vcf data/VCF/temp/BMAG006.recode.vcf --out data/VCF/BMAG006 --hardy

# library BMAG007
vcftools --vcf data/VCF/temp/ONC.F7.recode.vcf --out data/VCF/temp/BMAG007 --keep data/VCF/BMAG007.ind --recode --recode-INFO-all

vcftools --vcf data/VCF/temp/BMAG007.recode.vcf --out data/VCF/BMAG007 --hardy

# library BMAG020
vcftools --vcf data/VCF/temp/ONC.F7.recode.vcf --out data/VCF/temp/BMAG020 --keep data/VCF/BMAG020.ind --recode --recode-INFO-all

vcftools --vcf data/VCF/temp/BMAG020.recode.vcf --out data/VCF/BMAG020 --hardy

# library BMAG021
vcftools --vcf data/VCF/temp/ONC.F7.recode.vcf --out data/VCF/temp/BMAG021 --keep data/VCF/BMAG021.ind --recode --recode-INFO-all

vcftools --vcf data/VCF/temp/BMAG021.recode.vcf --out data/VCF/BMAG021 --hardy

# library BMAG022
vcftools --vcf data/VCF/temp/ONC.F7.recode.vcf --out data/VCF/temp/BMAG022 --keep data/VCF/BMAG022.ind --recode --recode-INFO-all

vcftools --vcf data/VCF/temp/BMAG022.recode.vcf --out data/VCF/BMAG022 --hardy

# library BMAG023
vcftools --vcf data/VCF/temp/ONC.F7.recode.vcf --out data/VCF/temp/BMAG023 --keep data/VCF/BMAG023.ind --recode --recode-INFO-all

vcftools --vcf data/VCF/temp/BMAG023.recode.vcf --out data/VCF/BMAG023 --hardy

# library BMAG024
vcftools --vcf data/VCF/temp/ONC.F7.recode.vcf --out data/VCF/temp/BMAG024 --keep data/VCF/BMAG024.ind --recode --recode-INFO-all

vcftools --vcf data/VCF/temp/BMAG024.recode.vcf --out data/VCF/BMAG024 --hardy

# library BMAG025
vcftools --vcf data/VCF/temp/ONC.F7.recode.vcf --out data/VCF/temp/BMAG025 --keep data/VCF/BMAG025.ind --recode --recode-INFO-all

vcftools --vcf data/VCF/temp/BMAG025.recode.vcf --out data/VCF/BMAG025 --hardy

# library BMAG001
vcftools --vcf data/VCF/temp/ONC.F7.recode.vcf --out data/VCF/temp/BMAG001 --keep data/VCF/BMAG001.ind --recode --recode-INFO-all

vcftools --vcf data/VCF/temp/BMAG001.recode.vcf --out data/VCF/BMAG001 --hardy

# library BMAG008
vcftools --vcf data/VCF/temp/ONC.F7.recode.vcf --out data/VCF/temp/BMAG008 --keep data/VCF/BMAG008.ind --recode --recode-INFO-all

vcftools --vcf data/VCF/temp/BMAG008.recode.vcf --out data/VCF/BMAG008 --hardy

# library BMAG010
vcftools --vcf data/VCF/temp/ONC.F7.recode.vcf --out data/VCF/temp/BMAG010 --keep data/VCF/BMAG010.ind --recode --recode-INFO-all

vcftools --vcf data/VCF/temp/BMAG010.recode.vcf --out data/VCF/BMAG010 --hardy

# library BMAG013
vcftools --vcf data/VCF/temp/ONC.F7.recode.vcf --out data/VCF/temp/BMAG013 --keep data/VCF/BMAG013.ind --recode --recode-INFO-all

vcftools --vcf data/VCF/temp/BMAG013.recode.vcf --out data/VCF/BMAG013 --hardy

# library BMAG009
vcftools --vcf data/VCF/temp/ONC.F7.recode.vcf --out data/VCF/temp/BMAG009 --keep data/VCF/BMAG009.ind --recode --recode-INFO-all

vcftools --vcf data/VCF/temp/BMAG009.recode.vcf --out data/VCF/BMAG009 --hardy

# library BMAG017
vcftools --vcf data/VCF/temp/ONC.F7.recode.vcf --out data/VCF/temp/BMAG017 --keep data/VCF/BMAG017.ind --recode --recode-INFO-all

vcftools --vcf data/VCF/temp/BMAG017.recode.vcf --out data/VCF/BMAG017 --hardy
```

Compare data set after filtering -

Data set contains 418 individuals and 36633 loci.

**FILTER 8: Excess heterozygosity**

Identify SNPs with more than 0.5 heterozygosity and significant excess heterozygosity.

Fig 22: Distribution of observed heterozygosity per locus.

Identify loci with heterozygosity > 0.5, then correct p-values for multiple comparisons using Benjamini-Hochberg method. 3410 loci flagged to be removed from the data set.

Table 7a: Number of loci with Ho > 0.75 per library.

| LIB | n |
| --- | --- |
| BMAG001 | 2642 |
| BMAG006 | 2960 |
| BMAG007 | 3043 |
| BMAG010 | 2921 |
| BMAG013 | 2902 |
| BMAG017 | 2926 |
| BMAG020 | 2977 |
| BMAG021 | 2980 |
| BMAG022 | 2940 |
| BMAG023 | 2965 |
| BMAG024 | 3000 |
| BMAG025 | 3010 |

Table 7b: Number of loci with H0 > 0.75 in only one library per library

| LIB | n |
| --- | --- |
| BMAG001 | 1 |
| BMAG006 | 14 |
| BMAG007 | 59 |
| BMAG010 | 33 |
| BMAG013 | 19 |
| BMAG017 | 15 |
| BMAG020 | 11 |
| BMAG021 | 3 |
| BMAG022 | 4 |
| BMAG023 | 13 |
| BMAG024 | 20 |
| BMAG025 | 29 |

Compare patterns by library to determine loci that could cause library effects due to excess heterozygosity. Most loci that are out in one library are out in multiple libraries and overall loci flagged for removal in a library and/or overall).

There is a known library effect for BMAG002 (strongest), BMAG008, and BMAG009 that is due to excess heterozygosity in those individuals (comparison of Fis values for duplicates from BMAG002 and BMAG017 show that Fis values are vastly different; comparing duplicates from other libraries to BMAG017 that is not the case) - using the correct p-values the number of loci out in those libraries is comparable to other libraries, using the uncorrect p-value those numbers are comparatively larger.

Total of 6762 loci are flagged for removal.

```
vcftools --vcf data/VCF/temp/ONC.F7.recode.vcf --out data/VCF/temp/ONC.F8 --exclude-positions data/VCF/hetexcess.loci --recode --recode-INFO-all

vcftools --vcf data/VCF/temp/ONC.F8.recode.vcf --out data/VCF/ONC.F8 --depth
vcftools --vcf data/VCF/temp/ONC.F8.recode.vcf --out data/VCF/ONC.F8 --site-mean-depth
vcftools --vcf data/VCF/temp/ONC.F8.recode.vcf --out data/VCF/ONC.F8 --missing-indv
vcftools --vcf data/VCF/temp/ONC.F8.recode.vcf --out data/VCF/ONC.F8 --missing-site
vcftools --vcf data/VCF/temp/ONC.F8.recode.vcf --out data/VCF/ONC.F8 --het
vcftools --vcf data/VCF/temp/ONC.F8.recode.vcf --out data/VCF/ONC.F8 --geno-depth
```

Compare data set -

Data set contains 33223 SNP sites and 418 individuals.

**FILTER 9: Identify and filter LQ individuals**

Use genotype depth file to identify individuals with high variance in read depth across loci.

Compare distribution of depth within individuals.

Fig 23: Heatmap indicating genotype depth per locus for each individual by run type.

Table 8a: Number of individuals per run type remaining in the data set (duplicate samples excluded from counts).

| RUN | n |
| --- | --- |
| F | 277 |
| L | 27 |
| S | 69 |
| W | 29 |

Compare distribution of depth of individuals grouped by library to identify potential library effects.

Fig 24: Heatmap indicating genotype depth per locus for each individual by library.

Table 8b: Number of individuals per library remaining in the data set.

| LIB | n |
| --- | --- |
| BMAG001 | 29 |
| BMAG006 | 25 |
| BMAG007 | 31 |
| BMAG010 | 34 |
| BMAG013 | 31 |
| BMAG017 | 32 |
| BMAG020 | 45 |
| BMAG021 | 45 |
| BMAG022 | 43 |
| BMAG023 | 43 |
| BMAG024 | 31 |
| BMAG025 | 29 |

Compare distribution of genotype depths (across all individuals).

Fig 25: Comparison of genotype depth across individuals. Genotype depths < 5 are coded as missing.

Determine variance in depth w/in individuals to identify problematic individuals remaining in the data set.

Fig 26: Comparison of mean/median depth, variance in depth, and missing data within individuals.

Compare depth, missing data, and individual heterozygosity levels.

Fig 27: Comparison of depth, missing data, and heterozygosity.

Compare patterns of distribution of Fis (homozygosity of individuals) by library and run type to identify artifacts but not eliminate real biological signal.

Fig 28: Distribution of Fis per individual grouped by library. Dashed red line is the mean level of Fis across all individuals.

Fis can also be a biological signal. Smaller populations could have higher Fis (erosion of genetic diversity) and hybridization of populations could lead to excess heterozygosity.

Fig 29: Distribution of Fis per individual grouped by run type. Dashed red line is the mean level of Fis across all individuals.

Winter run individuals appear to have elevated Fis, which makes biologically sense since they have a much smaller population which can lead to an increase in homozogysity due to drift and/or inbreeding. Spring run individuals having a slight left shift in their Fis values makes sense to since they are potentially hybridizing with Fall run individuals

Fig 30: Distribution of Fis across all individuals.

In general, pervasiveness of paralogs and contamination can lead to a decrease in Fis (excess homozogysity). Biologically, this would occur if there is outbreeding/admixture between two different demes. Excess homozogygosity can be caused by false homozygotes e.g. due to low depth.

Flag low quality individuals with missing data > 0.5, Fis < -0.05, or Fis > 0.4

**FILTER 10: Missing data by sample location**

```
vcftools --vcf data/VCF/temp/ONC.F8.recode.vcf --out data/VCF/temp/ONC.F9 --remove data/VCF/F9_LQ.indv --recode --recode-INFO-all

# output per run ---------------------------------------------------------

# Fall
vcftools --vcf data/VCF/temp/ONC.F9.recode.vcf --out data/VCF/temp/FALL2 --keep data/VCF/F.ind --recode --recode-INFO-all

vcftools --vcf data/VCF/temp/FALL2.recode.vcf --out data/VCF/FALL2 --missing-site

# Late Fall
vcftools --vcf data/VCF/temp/ONC.F9.recode.vcf --out data/VCF/temp/LATE-FALL2 --keep data/VCF/L.ind --recode --recode-INFO-all

vcftools --vcf data/VCF/temp/LATE-FALL2.recode.vcf --out data/VCF/LATE-FALL2 --missing-site

# Spring
vcftools --vcf data/VCF/temp/ONC.F9.recode.vcf --out data/VCF/temp/SPRING2 --keep data/VCF/S.ind --recode --recode-INFO-all

vcftools --vcf data/VCF/temp/SPRING2.recode.vcf --out data/VCF/SPRING2 --missing-site

# Winter
vcftools --vcf data/VCF/temp/ONC.F9.recode.vcf --out data/VCF/temp/WINTER2 --keep data/VCF/W.ind --recode --recode-INFO-all

vcftools --vcf data/VCF/temp/WINTER2.recode.vcf --out data/VCF/WINTER2 --missing-site
```

Compare levels missing data per locus per sample region to avoid non-random patterns of missing data.

Fig 31: Genotype call rate per run type.

Table 9: Number of loci called in < 85% of individuals of a run type.

| POP | n |
| --- | --- |
| LATE-FALL2 | 227 |
| SPRING2 | 824 |
| WINTER2 | 289 |

Identify loci with low genotype call rate in a given sample location (> 15 % missing data) to remove loci from data set. Remove all singleton SNPs.

```
vcftools --vcf data/VCF/temp/ONC.F9.recode.vcf --out data/VCF/temp/ONC.F10 --exclude-positions data/VCF/LQ_F10.loci --recode --recode-INFO-all

vcftools --vcf data/VCF/temp/ONC.F10.recode.vcf --out data/VCF/ONC.F10 --site-quality
vcftools --vcf data/VCF/temp/ONC.F10.recode.vcf --out data/VCF/ONC.F10 --depth
vcftools --vcf data/VCF/temp/ONC.F10.recode.vcf --out data/VCF/ONC.F10 --site-mean-depth
vcftools --vcf data/VCF/temp/ONC.F10.recode.vcf --out data/VCF/ONC.F10 --missing-indv
vcftools --vcf data/VCF/temp/ONC.F10.recode.vcf --out data/VCF/ONC.F10 --missing-site
vcftools --vcf data/VCF/temp/ONC.F10.recode.vcf --out data/VCF/ONC.F10 --het
vcftools --vcf data/VCF/temp/ONC.F10.recode.vcf --out data/VCF/ONC.F10 --geno-depth

vcftools --vcf data/VCF/temp/ONC.F10.recode.vcf --out data/VCF/ONC.F10 --012
```

Compare data set after filtering:

Fig 32: Distribution of missing data, read depth, and heterozygosity for loci and individuals.

#### Final thresholds values for filtered data set:

- minimum sequence quality (minQ): 20
- minimum genotype quality (minGQ): 20
- minimum genotype call rate per locus (geno): 90%
- minimum genotype call rate by population: 85%
- maximum allowed missing data per individual (missInd): 50%
- minimum genotype call rate by library: 50%
- minimum genotype depth (minD): 5
- minimum mean depth per locus (m-minD): 15
- minimum mean depth per locus for loci grouped by library: 10
- mean maximum depth (m-minD): 70
- filtered for allelic balance, mapping quality, depth/quality ratio, and excess heterozygosity.
- only bi-allelic SNPs retained.

Data set contains 31897 SNP sites and 413 individuals.

### Haplotyping

#### Prep for haplotyping

Create list of individuals retained in final vcf file.

```
vcftools --vcf data/VCF/temp/ONC.F10.recode.vcf --out data/HAPLOTYPING/temp/ONC --recode --recode-INFO-all

scr/vcflib/bin/vcfsamplenames data/HAPLOTYPING/temp/ONC.recode.vcf > data/HAPLOTYPING/temp/ONC.individuals
```

Use `ONC.individuals` to create `popmap` as a tab-separated file of individuals and their population designation, with one individual per line (make sure UNIX format). This file is needed to write the genepop file, if not provided the script will run through the process but not write a genepop file and place into same folder `rad_haplotyper.pl` will be run from.

Place all necessary files in `data/HAPLOTYPING/temp` directory (bam, fastq, reference.fasta, vcf-file)

```
ln -s $HOME/CHINOOK/data/SEQ/*/*.fq.gz $HOME/CHINOOK/data/HAPLOTYPING/temp/

ln -s $HOME/CHINOOK/data/SNP_CALLING/*.bam* $HOME/CHINOOK/data/HAPLOTYPING/temp/

# need to rename bam files so sample names as fastq (i.e. remove the fil part)
cd data/HAPLOTYPING/temp/

cp $HOME/CHINOOK/data/SNP_CALLING/mapped.bed $HOME/CHINOOK/data/HAPLOTYPING/temp/

cp $HOME/CHINOOK/data/REF/CHRIST2018/reference.fasta* $HOME/CHINOOK/data/HAPLOTYPING/temp/
```

Make haplotyping process more efficient by filtering `mapped.bed` file to only consist of intervals that contain SNPs in the filtered data set.

#### Run Haplotyper

```
export PATH=$PATH:$HOME/miniconda3/bin

# make sure rad_haplotyper environment is installed
# conda create -n rad_haplotyper_env rad_haplotyper

# open bioconda environment
source activate $HOME/miniconda3/envs/rad_haplotyper_env

    # run rad_haplotyper
    perl rad_haplotyper.pl -v ONC.recode.vcf -b mapped_filtered.bed -r reference.fasta -x 30 -d 10 -z 3 -m .25 -g ONC.gen -p popmap -e --vcfout ONC.haplotyped.vcf

# close bioconda environment
source deactivate
```

Remove intermediate files and copy files for analysis into `data/Haplotyping/`-folder.

```
# delete softlinks to fastq/bam files and unneeded log-files
rm $HOME/CHINOOK/data/HAPLOTYPING/temp/*.fq.gz
rm $HOME/CHINOOK/data/HAPLOTYPING/temp/*.bam*
rm $HOME/CHINOOK/data/HAPLOTYPING/temp/reference.fasta*
rm $HOME/CHINOOK/data/HAPLOTYPING/temp/log.txt
rm $HOME/CHINOOK/data/HAPLOTYPING/temp/hap_loci.txt
rm $HOME/CHINOOK/data/HAPLOTYPING/temp/haplo_recode.log
rm $HOME/CHINOOK/data/HAPLOTYPING/temp/fail.log
rm $HOME/CHINOOK/data/HAPLOTYPING/temp/hap_log.out
rm $HOME/CHINOOK/data/HAPLOTYPING/temp/haplo_dump.out
rm $HOME/CHINOOK/data/HAPLOTYPING/temp/allele_dump.out
rm $HOME/CHINOOK/data/HAPLOTYPING/temp/snp_dump.out
rm $HOME/CHINOOK/data/HAPLOTYPING/temp/rad_haplotyper.pl

# copy genepop files and stats files into main haplotyping directory
cp $HOME/CHINOOK/data/HAPLOTYPING/temp/ind_stats.out $HOME/CHINOOK/data/HAPLOTYPING/
cp $HOME/CHINOOK/data/HAPLOTYPING/temp/stats.out $HOME/CHINOOK/data/HAPLOTYPING/
cp $HOME/CHINOOK/data/HAPLOTYPING/temp/ONC.gen $HOME/CHINOOK/data/HAPLOTYPING/
cp $HOME/CHINOOK/data/HAPLOTYPING/temp/codes.ONC.gen $HOME/CHINOOK/data/HAPLOTYPING/
cp $HOME/CHINOOK/data/HAPLOTYPING/temp/ONC.haplotyped.vcf $HOME/CHINOOK/data/HAPLOTYPING/
```

### Haplotype filtering

#### Overview of haplotyping success

Comparison of the number of loci that were filtered due to excess number of missing data or suspected paralogs during the haplotyping process to those that passed. Haplotyer was run without any threshold values except `-m 0.25`, i.e. loci missing in > 75% individuals are filtered.

The `genepop`-file only contains loci that passed haplotyping stage.

Fig 34: Number of loci that passed haplotyping process.

Filtered loci were almost all removed due to individuals being flagged for that locus for coverage/genotyping error issues (resulting in high missing data).

Fig 35: Haplotyping success and distribution of number of loci flagged as potential paralogs/genotyping error. Loci that did not pass haplotyping process have already been removed.

14654 loci passed haplotyping process; data set contains 413 individuals:

Fig 36: Haplotyping stats per individual.

#### Identify threshold values to filter data set

**Genotype call rate**

Fig 37: Missing data per locus (genotypes flagged as potential paralogs or as affected by genotyping error due to low coverage are flagged as missing.

Table 10: Loci with < 90% genotype call rate.

| Prop\_Haplotyped < 0.9 | n |
| --- | --- |
| FALSE | 14518 |
| TRUE | 136 |

136 loci were flagged as for genotype call rate < 0.9.

**Number of individuals flagged as paralog per locus**

Loci are flagged as possible paralogs for an individuals when more than the expected number of haplotypes based on the SNP genotype call (homozygote, heterozygote) are detected, which can be indicative of paralogs (reads from two different loci being mapped to the same location on the genome).

Cut-offs for loci flagged in

- 1 % of individuals: 4.13
- 5 % of individuals: 20.65

Fig 38: Proportion of individuals flagged as potential paralog.

Table 11: Loci flagged as possible paralogs in > 5 individuals.

| Poss\_Paralog > 5 | n |
| --- | --- |
| FALSE | 14602 |
| TRUE | 52 |

Flag loci that are flagged as potential paralogs in 5 or more individuals. 52 loci were flagged as possible paralogs.

**Number of SNPs & Haplotypes per locus**

Each locus varies in the number of SNPs detected which determines the number of haplotypes expected in that population.

Fig 39: Distribution of number of SNPs per locus. Dark red dashed line is the mean number of SNPs per locus.

Filtering loci based on number of SNPs contained on that locus could bias the data set as loci with high recombination may be removed. On the other hand, assuming an approximate length of 150 bp loci with more than 15 SNPs would mean that 10% of bases are a polymorphisms.

Fig 40: Relationship of number of times a locus is flagged as a potential paralog and number of SNPs/haplotypes per locus.

Assuming that mutation is the only mechanism resulting in new haplotypes, the maximum expected number of haplotypes per locus is number of SNPs N + 1.

Fig 41: Comparison of number of (extra) haplotypes per locus and number of times a locus was flagged as a potentials paralogs or affected by genotyping error/low coverage.

**Number of times locus is flagged as low coverage/genotyping error**

After combining SNPs on the same contig during the haplotyping process, it is possible to observe fewer than the expected number of haplotypes based on the genotype call (heterozygote/homozygote). When this occurs, that genotype is coded as missing. For each locus the number of individuals for which is it flagged as a potential a potential genotyping error or suffering from null alleles due to low coverage detected for a given locus is recorded.

Fig 42: Distribution of number of individuals a locus is flagged as potentially affected by genotyping error/low depth. The red dashed line indicates the mean number of individuals flagged, the blue dashed lines indicates 1% of individuals being flagged.

Table 12: Number of loci that have been flagged as affected by genotyping error/low coverage in > 5 individuals.

| Low\_Cov.Geno\_Err > 5 | n |
| --- | --- |
| FALSE | 14521 |
| TRUE | 133 |

Loci with pronounced patterns of genotyping error likely due to low coverage will have low success rate for genotyping, i.e. they will be caught in missing data filters. Coverage issues are likely genotype specfic; previous filters have targeted loci and individuals with high variance in coverage and suspect genotpes have been coded as missing, i.e. this filter need not be very conservative, regardless, loci that are repeatedly being flagged as problematic should be removed.

**Individuals with low proportion of successfully haplotyped loci**

Loci that are not successfully haplotyped in an individual are coded as missing for that individual. Problematic individual can be identified as having a high proportion of missing data.

Fig 43: Distribution of missing data per individual

Table 13: Number of individuals genotyped for < 50% loci.

| Prop\_Success < 0.5 | n |
| --- | --- |
| FALSE | 399 |
| TRUE | 14 |

Problem individuals can be identified by choosing a cut-off value for the minium proportion of sucessfully haplotyped loci and excluding individuals below that threshold value. Removing loci with low haplotyping success with decrease the amount of missing data. Therefore, final missing data cut-offs should be applied *after* removing those loci from the data set, though it can be helpful to flag potentially problematic individuals based on their haplotyping stats at this point.

**Possible paralogs per individual**

Individuals with high proportion of loci flagged as possible paralogs should be flagged as potential problem individuals.

1% of loci: 146.54

Fig 44: Distribution of no of loci flagged as potetnial paralogs within an individal

Table 14: Number of individuals with > 145 loci (1% of loci) flagged as potential paralogs.

| Poss\_Paralogs > 145 | n |
| --- | --- |
| FALSE | 413 |

Cut-off for individuals with loci flagged paralogs in > 1% of loci is 146.54. The highest number of flagged loci in an individuals is 23, which is equivalent to 0.16% of loci.

**Individuals with high proportion of genotyping error/low coverage issues**

Fig 45: Number of loci flagged as potential genotyping error/low coverage per individual.

Loci that are systematically affected by potential low coverage/genotyping error have been removed from the data set, individuals that are systematically affected will have more missing data which should be adjusted after problematic individuals have been removed from the data set.

#### Filter flagged loci

Load genepop file with haplotyped loci and remove flagged loci from the data set.

```
##
##  Converting data from a Genepop .gen file to a genind object...
##
##
## File description:  ONC.gen
##
## ...done.
```

```
## haplotyped data set
```

```
## /// GENIND OBJECT /////////
##
##  // 413 individuals; 14,654 loci; 32,331 alleles; size: 59.6 Mb
##
##  // Basic content
##    @tab:  413 x 32331 matrix of allele counts
##    @loc.n.all: number of alleles per locus (range: 1-11)
##    @loc.fac: locus factor for the 32331 columns of @tab
##    @all.names: list of allele names for each locus
##    @ploidy: ploidy of each individual  (range: 2-2)
##    @type:  codom
##    @call: read.genepop(file = "data/HAPLOTYPING/ONC.gen", ncode = 3L, quiet = FALSE)
##
##  // Optional content
##    @pop: population of each individual (group size range: 413-413)
```

```
## haplotyped data set after removing flagged loci
```

```
## /// GENIND OBJECT /////////
##
##  // 413 individuals; 14,450 loci; 31,574 alleles; size: 58.3 Mb
##
##  // Basic content
##    @tab:  413 x 31574 matrix of allele counts
##    @loc.n.all: number of alleles per locus (range: 1-9)
##    @loc.fac: locus factor for the 31574 columns of @tab
##    @all.names: list of allele names for each locus
##    @ploidy: ploidy of each individual  (range: 2-2)
##    @type:  codom
##    @call: .local(x = x, i = i, j = j, loc = ..1, drop = drop)
##
##  // Optional content
##    @pop: population of each individual (group size range: 413-413)
```

A total of 4025 SNP-containing loci were removed from the data set after haplotyping (this includes the loci filtered during the haplotyping process itself due to missing data).

Table 15: Samples remaining in data set by run type and river location.

| LOCATION | F | L | W | S |
| --- | --- | --- | --- | --- |
| USR | 2 | 21 | 26 | NA |
| BTC | 2 | 2 | NA | NA |
| COL | 30 | 2 | NA | NA |
| MIL | 20 | NA | NA | 16 |
| DER | 15 | NA | NA | 27 |
| BUT | 21 | 1 | NA | 19 |
| FRH | 27 | NA | NA | 7 |
| NIM | 30 | NA | NA | NA |
| MKH | 30 | NA | NA | NA |
| STN | 23 | NA | NA | NA |
| TOU | 30 | NA | NA | NA |
| MRH | 15 | NA | NA | NA |
| MER | 31 | NA | NA | NA |

### Heterozygosity and HWE

#### Compare patterns of heterozygosity

Generate summary statistics overall and for individuals grouped by run type and location (remove duplicates before calculating).

Compare observed (Ho) and expected (He) heterozygosity for all individuals across all populations.

Fig 46: Observed and expected heterozygosity per locus across all individuals.

Identify loci that are now monomorphic (e.g. because individuals were removed during SNP filtering since no maf filter was applied they stay in the data set). 1437 monomorphic loci removed.

Compare distribution of observed heterozygosity (Ho) in each putative population:

Fig 47: Observed and expected heterozygosity per locus for individuals grouped by run type and river location.

#### Test for deviations from Hardy-Weinberg equilibrium within each group

```
# groups being tested
popNames(tmp)

# run HWE test per group and overall
sep <- seppop(tmp)

HWE <-sep %>%
   lapply(hw.test, B = 100)

HWE[["ALL"]] <- hw.test(tmp, B = 100)

# convert to data.frames
hwe <- list()

for (m in names(HWE)) {
   hwe[[m]] <- as.data.frame(HWE[[m]]) %>%
   rename(pval = Pr.exact) %>%
   rownames_to_column("LOCUS") %>%
   select(LOCUS, pval)
}

hwe <- ldply(hwe, data.frame) %>%
   rename(POP = `.id`) %>%
   mutate(HWE = ifelse(pval <= 0.05, "OUT", "IN"),
          POP = case_when(POP == "F" ~ "Fall",
                          POP == "L" ~ "Late-Fall",
                          POP == "S" ~ "Spring",
                          POP == "W" ~ "Winter",
                          POP == "ALL" ~ "OVERALL"))

write_delim(hwe, "results/by_run.hwe", delim = "\t")
```

Compare number of loci flagged as out of HWE (individuals grouped by run).

Fig 48: Number of loci flagged as out of HWE per run type.

Identify loci that are consistently flagged as out of HWE across run types.

Fig 49: Number of times a locus is flagged as being out of HWE.

No loci removed based on HWE because groups of “randomly mating” individuals are difficult to discern *a priori* (tributaries within run types but some straying). As expected, more loci are flagged as out of HWE when testing across all individuals, and for larger samples encompassing individuals from more different tributaries (Fall, Spring).

### Compare duplicate individuals

Fig 50: Genotyping error frequency per individual.

Identify loci consistently affected by genotyping error.

Fig 51: For discordant loci only: Comparison of genotyping error per locus (freq), number of haplotypes per locus (Haplotypes), number of pairs locus is affected by genotyping error (n), number of times locus is flagged as possible paralog (Poss\_Paralog), proportion of individuals haplotyped in (Prop\_Haplotyped), and number of SNP sites per locus (Sites).

Remove loci affected by genotyping error in > 50% of pairs and one individual per duplicate set.

```
## haplotyped data set after filtering complete
```

```
## /// GENIND OBJECT /////////
##
##  // 397 individuals; 13,000 loci; 30,093 alleles; size: 53.6 Mb
##
##  // Basic content
##    @tab:  397 x 30093 matrix of allele counts
##    @loc.n.all: number of alleles per locus (range: 2-9)
##    @loc.fac: locus factor for the 30093 columns of @tab
##    @all.names: list of allele names for each locus
##    @ploidy: ploidy of each individual  (range: 2-2)
##    @type:  codom
##    @call: .local(x = x, i = i, j = j, loc = ..1, drop = drop)
##
##  // Optional content
##    @pop: population of each individual (group size range: 397-397)
##    @strata: a data frame with 7 columns ( LIB_ID, LIB, SAMPLE_ID, RUN, YEAR, INDV, ... )
```

### Final data set

Write files with filtered data set.

```
## ################################################################################
## ######################### radiator::tidy_genomic_data ##########################
## ################################################################################
## ################################################################################
## ########################### radiator::filter_monomorphic #######################
## ################################################################################
## ######################## filter_monomorphic completed ##########################
## ################################### RESULTS ####################################
## ######################### tidy_genomic_data completed ##########################
```

Write files with individuals and contigs still contained in data set and use to filter vcf file.

Filter vcf file to contain only SNPs (N = 18419) from SNP-containing loci (13000) retained in data set and output in 012 format.

```
# load modules for vcf filtering
module purge
module load GCC/6.4.0-2.28  OpenMPI/2.1.2
module load VCFtools/0.1.15-Perl-5.26.1

# retain only individuals and loci in the filtered/haplotyped data sets
vcftools --vcf data/VCF/temp/ONC.F10.recode.vcf --out data/POPGEN/ONC.haps.filtered --keep data/VCF/filtered.ind --recode --recode-INFO-all

# write 012 output
vcftools --vcf data/POPGEN/ONC.haps.filtered.recode.vcf --out data/POPGEN/ONC.haps.filtered --012

# stat files
vcftools --vcf data/POPGEN/ONC.haps.filtered.recode.vcf --out data/VCF/ONC.haps.fil --depth
vcftools --vcf data/POPGEN/ONC.haps.filtered.recode.vcf --out data/VCF/ONC.haps.fil --site-mean-depth
vcftools --vcf data/POPGEN/ONC.haps.filtered.recode.vcf --out data/VCF/ONC.haps.fil --missing-indv
vcftools --vcf data/POPGEN/ONC.haps.filtered.recode.vcf --out data/VCF/ONC.haps.fil --missing-site
vcftools --vcf data/POPGEN/ONC.haps.filtered.recode.vcf --out data/VCF/ONC.haps.fil --het
vcftools --vcf data/POPGEN/ONC.haps.filtered.recode.vcf --out data/VCF/ONC.haps.fil --hardy
vcftools --vcf data/POPGEN/ONC.haps.filtered.recode.vcf --out data/VCF/ONC.haps.fil --site-quality
vcftools --vcf data/POPGEN/ONC.haps.filtered.recode.vcf --out data/VCF/ONC.haps.fil --geno-depth
```

Compare stats for final filtered data set:

Fig 52: Genotype call rate, missing data, and read depth across loci and individuals for final data set (SNPs retained in haplotyped data set).
