## Supplementary material for "Every cog and wheel: Identifying biocomplexity at the genomic and phenotypic level in a population complex of Chinook salmon": Supplementary Material 2.html (from another version of this preprint: version 1, 2021-03-28)

- 1 Genotyping
- 2 Assessment of population structure and differentiation
  - 2.1 k-mean clustering
  - 2.2 F-statistics
- 3 Estimates of effective population size
- 4 Assessment of genomic diversity
  - 4.1 Calculate measures of genetic diversity based on allele frequency and heterozygosity
  - 4.2 observed heterozygosity
  - 4.3 Expected heterozygosity
  - 4.4 Comparison of allelic richness
  - 4.5 Evenness of allelic diversity
  - 4.6 Nucleotide diversity \(\pi\) (pairwise differences)
  - 4.7 Fixed loci
  - 4.8 Singletons
  - 4.9 Loci variable in only one group of individuals (“private polymorphisms”)
  - 4.10 Private alleles

### 1 Genotyping

⊕Reads were mapped to the Christensen 2018 chinook genome and SNPs were called using freebayes. Filtered data set was haplotyped using rad\_haplotyper. Fst-outlier were identified using FDIST method implemented in Arlequin and bayescan.

```
## 
## FULL DATA SET: all SNP-containing loci/all individuals grouped by run type.
```

```
## /// GENIND OBJECT /////////
## 
##  // 386 individuals; 12,983 loci; 30,037 alleles; size: 52.2 Mb
## 
##  // Basic content
##    @tab:  386 x 30037 matrix of allele counts
##    @loc.n.all: number of alleles per locus (range: 2-9)
##    @loc.fac: locus factor for the 30037 columns of @tab
##    @all.names: list of allele names for each locus
##    @ploidy: ploidy of each individual  (range: 2-2)
##    @type:  codom
##    @call: .local(x = x, i = i, j = j, drop = drop)
## 
##  // Optional content
##    @pop: population of each individual (group size range: 21-270)
##    @strata: a data frame with 10 columns ( LIB, LIB_ID, SAMPLE_ID, YEAR, SOURCE, RUN, ... )
```

Number of samples per tributary and run type.

| LOCATION | Fall | Late-Fall | Winter | Spring |
| --- | --- | --- | --- | --- |
| COL | 30 | - | - | - |
| MIL | 20 | - | - | 16 |
| DER | 15 | - | - | 27 |
| BUT | 21 | - | - | 19 |
| FRH | 27 | - | - | 7 |
| NIM | 30 | - | - | - |
| MKH | 28 | - | - | - |
| STN | 23 | - | - | - |
| TOU | 30 | - | - | - |
| MRH | 15 | - | - | - |
| MER | 31 | - | - | - |
| USR | - | 21 | 26 | - |

\*For details on bioinformatics to genotype the data set see `Demultiplex.html` and `Genotyping.html`.

### 2 Assessment of population structure and differentiation

Population structure was explored using two methods, a clustering analysis based on genetic similarity and an assessment of population differentiation among individuals grouped *a priori* based on run type and tributary.

#### 2.1 k-mean clustering

Individuals were clustered into K = 2 – 10 groups using k-means clustering based on the PCA-transformed genotype matrix (i.e. no assumptions regarding Hardy-Weinberg or linkage disequilibrium) followed by a discriminant analysis of principle components (DAPC) to determine the membership probabilities of each sample to each inferred cluster as implemented in `adegenet` (Jombart, Devillard, and Balloux ⊕2010Jombart, Thibaut, Sébastien Devillard, and François Balloux. 2010. “Discriminant analysis of principal components: a new method for the analysis of genetically structured populations.” *BMC Genetics* 11 (1): 94. https://doi.org/10.1186/1471-2156-11-94.). During DAPC, the genotype matrix is first PCA-transformed, then samples are partitioned into a within- and between-group component to maximize discrimination between groups using linear discriminant analysis, and finally, the membership probability of each sample to each cluster is calculated. To ensure sufficient variance was retained to discriminate among groups but not overfit the data, the optimum number of principle components to retain was determined using a stratified cross-validation of DAPC.

Use k-means clustering to identify clusters based on genetic similarity (data transformed using PCA). For clustering it is appropriate to retain all PCs (all variance in the data set), as overfitting is not an issue.

Figure S1: AIC for data clustered into K = 1 - 40 groups using k-means clustering.

AIC is minimized for K=3.

Perform stratified cross-validation of DAPC using a range of retained PCs, while keeping the number of discriminant functions fixed for K = 2 - 10 to determine the most appropriate number of principle components to retain sufficient variance to discriminate but not overfit the data.

```
## define minimum and maximum K to cluster for ----
min_K <- 2

max_K <- 10

# number of repetitions for x-validation
n_rep <- 30

# range of values to loop over
range <- min_K:max_K

## loop over values for K ----

# list for cross validation plots
xVal_plots <- list()

# data frame for optimum PCs
opt_PC <- data.frame(matrix(ncol = 4, nrow = 0)) %>%
    rename(k_clust = X1,
           optPC = X2,
           variance = X3,
           mean_opt_success = X4)

# list for membership plots
memb_plots <- list()


# run loop
for (k in range){

    # set value for K
    k_clust <- k

    # cluster
    grp <- find.clusters.genind(gen, n.pca = 500, n.clust = k_clust, method = "kmeans")

    # scale allele frq/missing data replaced with mean
    X <- scaleGen(gen, NA.method = "mean")

    xval <- xvalDapc(X, grp$grp,
                     n.pca.max = 400, training.set = 0.9,
                     result = "groupMean", center = TRUE, scale = FALSE,
                     n.pca = NULL, n.rep = n_rep, xval.plot = FALSE)

    # extract assignment success for each xVal step
    PCs <- data.frame(N_PCs = xval[["Cross-Validation Results"]]$n.pca,
                      PROP_SUCCESS = xval[["Cross-Validation Results"]]$success)

    # create crossvalidation plot
    xVal_plots[[k_clust]] <- ggplot(PCs, aes(x = N_PCs, y = PROP_SUCCESS)) +
      stat_density2d(aes(fill = ..density..^0.25),
                     geom = "tile", contour = FALSE, n = 200) +   
      geom_point() +
      scale_fill_continuous(low = "white", high = "dodgerblue4") +
      scale_y_continuous(limits = c(0, 1)) +
      labs(title = glue("Cross validation K = {k_clust}"),
           x = "retained PCs", y = "assignment success") +
      theme_standard +
      theme(legend.position = "none")

    # optimum number of PCs to retain
    retain <- as.numeric(xval$`Number of PCs Achieving Highest Mean Success`)

    variance <- round(xval[["DAPC"]]$var*100, digits = 2)

    opt <- PCs %>%
      filter(N_PCs == retain)

    df <- as.data.frame(x = k_clust) %>%
      mutate(optPC = retain,
             variance = variance,
             mean_opt_success = round(mean(opt$PROP_SUCCESS), digits = 2))

    opt_PC <- opt_PC %>%
        bind_rows(df)

    # extract membership assignment probabilities
    grp_membership <- as.data.frame(xval[["DAPC"]]$posterior) %>%
      rownames_to_column("LIB_ID") %>%
      gather(key = GRP, value = MEMBSHIP, 2:(k_clust+1)) %>%
      left_join(strata) %>%
      unite(RUN_LOC, RUN, LOCATION, sep = "_", remove = FALSE) %>%
      arrange(RUN_LOC, LIB_ID) %>%
      mutate(LIB_ID = factor(LIB_ID, levels = unique(LIB_ID)))

    path <- as.character(glue("results/runs_K{k}.grp.membership"))

    write_delim(grp_membership, path, delim = "\t")

    # create membership plot
    memb_plots[[k_clust]] <- ggplot(grp_membership,
                                    aes(x = LIB_ID, y = MEMBSHIP, fill = GRP, color = GRP)) +
      geom_bar(stat = "identity") +
      facet_grid(. ~ RUN_LOC, scales = "free", space = "free") +
      scale_fill_manual(values = col_nine) +
      scale_color_manual(values = col_nine) +
      labs(x = "INDV", y = "memb. prob") +
      theme_standard +
      theme(axis.text.x = element_blank(),
        strip.text.x = element_text(size = 6, color = "black"),
        strip.text.y = element_text(size = 6, color = "black"),
        legend.position = "none")

}

write_delim(opt_PC, "results/kmeans_all-loci.optPCs", delim = "\t")
```

Retaining too few PCs may result in important variance not being retained and therefore not informing the clustering analysis, while retaining too many PCs results in overfitting, i.e. assignment success decreases.

Proportion of test individuals correctly assigned to their cluster of origin based on number of retained PCs for K = 2 - 9.

Table S1: The optimum number of principle components, %-variance retained and the mean optimum assignment succes for K = 2 - 10 clusters.

| k\_clust | optPC | variance | mean\_opt\_success |
| --- | --- | --- | --- |
| 2 | 50 | 75.10 | 1.00 |
| 3 | 50 | 63.49 | 1.00 |
| 4 | 50 | 36.98 | 0.99 |
| 5 | 100 | 36.98 | 0.98 |
| 6 | 50 | 21.77 | 0.96 |
| 7 | 50 | 21.77 | 0.96 |
| 8 | 100 | 36.98 | 0.91 |
| 10 | 20 | 11.62 | 0.89 |
| 9 | 150 | 50.81 | 0.87 |

Genetic data is scaled and centered, then transformed using a PCA. Retained PCs are then transmitted to a linear discrimant analysis.

Calculate membership probability of each individual to per cluster.

Membership probability of each individual to clusters identified using k-means hierarchical clustering (equiavalent to Figure 2 in manuscript).

Tributaries are grouped within runs by tributary from north to south as Fall\_BUT, Fall\_COL, Fall\_DER, Fall\_FRH, Fall\_MER, Fall\_MIL, Fall\_MKH, Fall\_MRH, Fall\_NIM, Fall\_STN, Fall\_TOU, Late-Fall\_USR, Spring\_BUT, Spring\_DER, Spring\_FRH, Spring\_MIL, and Winter\_USR (individual panels).

#### 2.2 F-statistics

Weir & Cockerham’s unbiased estimator of FST (Weir and Cockerham ⊕1984Weir, B. S., and C. Clark Cockerham. 1984. “Estimating F-Statistics for the Analysis of Population Structure.” *Evolution* 38 (6): 1358. https://doi.org/10.2307/2408641.) was used to calculate population differentiation among individuals grouped *a priori* by run type within tributary.

To test for genetic heterogeneity in the data set, global FST was calculated across all groups.

```
# group genetic data for comparison
setPop(gen) <- ~RUN_LOC

pop <- popNames(gen)

# calculate Fst
tidy <- tidy_genomic_data(data = gen, filename = NULL, parallel = 65)

fst <- assigner::fst_WC84(data = tidy,
                          pop.levels = pop,
                          holdout.samples = NULL,
                          pairwise = TRUE,
                          ci = TRUE,
                          iteration.ci = 1000,
                          quantiles.ci = c(0.025, 0.975),
                          digits = 9,
                          verbose = TRUE)

# write results
write_delim(fst$fst.overall, "results/trib_allLoc.globalfst.ci", delim = "\t")

write_delim(fst[["fst.ranked"]], "results/trib_ranked.perlocus.fst", delim = "\t")

write_delim(fst[["fis.markers"]], "results/trib_perlocus.fis", delim = "\t")

write_delim(fst$pairwise.fst, "results/trib_allLoc.fst.ci", delim = "\t")
```

**Significant heterogeneity** was detected among individuals grouped by run/tributary. Significance was determined using 95% confidence intervals around each estimate generated by re-sampling loci 1,000 times using `assigner` (Gosselin, Anderson, and Bradbury ⊕2016Gosselin, T, Eric C Anderson, and Ian R Bradbury. 2016. “assigner: Assignment Analysis with GBS/RAD Data using R.” *R Package*. https://doi.org/doi : 10.5281/zenodo.51453.).

Global Fst and 95% CI for individuals grouped by run type.

| FST | N\_MARKERS | CI\_LOW | CI\_HIGH |
| --- | --- | --- | --- |
| 0.0318754 | 12886 | 0.0309514 | 0.0328094 |

Pairwise FST was calculated as a *post hoc* test for pairwise differences among groups. Significance was determined using 95% confidence intervals around each estimate generated by re-sampling loci 1,000 times.

Table S2: Pairwise Fst among runs basins.

| POP1 | POP2 | COMP | N\_MARKERS | FST | CI\_LOW | CI\_HIGH |
| --- | --- | --- | --- | --- | --- | --- |
| F\_COL | F\_STN | F-F | 10328 | 0.005 | 0.004 | 0.006 |
| F\_TOU | F\_COL | F-F | 10432 | 0.005 | 0.004 | 0.006 |
| F\_NIM | F\_COL | F-F | 10479 | 0.005 | 0.004 | 0.006 |
| F\_MER | F\_COL | F-F | 10495 | 0.004 | 0.003 | 0.005 |
| F\_MKH | F\_COL | F-F | 10426 | 0.004 | 0.003 | 0.005 |
| F\_MRH | F\_COL | F-F | 10129 | 0.004 | 0.003 | 0.005 |
| F\_BUT | F\_COL | F-F | 10250 | 0.004 | 0.003 | 0.005 |
| F\_FRH | F\_STN | F-F | 10250 | 0.003 | 0.002 | 0.004 |
| F\_FRH | F\_BUT | F-F | 10186 | 0.003 | 0.002 | 0.004 |
| F\_MIL | F\_STN | F-F | 9896 | 0.003 | 0.002 | 0.004 |
| F\_FRH | F\_NIM | F-F | 10433 | 0.003 | 0.002 | 0.003 |
| F\_FRH | F\_TOU | F-F | 10387 | 0.002 | 0.002 | 0.003 |
| F\_FRH | F\_COL | F-F | 10545 | 0.002 | 0.002 | 0.003 |
| F\_MRH | F\_STN | F-F | 9688 | 0.002 | 0.001 | 0.003 |
| F\_DER | F\_STN | F-F | 9784 | 0.002 | 0.001 | 0.003 |
| F\_MIL | F\_COL | F-F | 10249 | 0.002 | 0.001 | 0.003 |
| F\_FRH | F\_MER | F-F | 10417 | 0.002 | 0.001 | 0.003 |
| F\_MKH | F\_FRH | F-F | 10358 | 0.002 | 0.001 | 0.003 |
| F\_MIL | F\_TOU | F-F | 10068 | 0.002 | 0.001 | 0.003 |
| F\_MIL | F\_NIM | F-F | 10146 | 0.002 | 0.001 | 0.003 |
| F\_DER | F\_COL | F-F | 10130 | 0.002 | 0.001 | 0.003 |
| F\_MIL | F\_MER | F-F | 10151 | 0.002 | 0.001 | 0.002 |
| F\_MKH | F\_MIL | F-F | 10034 | 0.002 | 0.001 | 0.002 |
| F\_DER | F\_NIM | F-F | 10047 | 0.002 | 0.001 | 0.002 |
| F\_MRH | F\_BUT | F-F | 9600 | 0.002 | 0.000 | 0.003 |
| F\_BUT | F\_STN | F-F | 9848 | 0.001 | 0.001 | 0.002 |
| F\_FRH | F\_MIL | F-F | 10227 | 0.001 | 0.001 | 0.002 |
| F\_FRH | F\_MRH | F-F | 10026 | 0.001 | 0.000 | 0.002 |
| F\_BUT | F\_NIM | F-F | 10106 | 0.001 | 0.000 | 0.002 |
| F\_FRH | F\_DER | F-F | 10113 | 0.001 | 0.000 | 0.002 |
| F\_DER | F\_TOU | F-F | 9939 | 0.001 | 0.000 | 0.002 |
| F\_MIL | F\_BUT | F-F | 9798 | 0.001 | 0.000 | 0.002 |
| F\_MRH | F\_MIL | F-F | 9594 | 0.001 | 0.000 | 0.002 |
| F\_MKH | F\_STN | F-F | 10061 | 0.001 | 0.000 | 0.002 |
| F\_MRH | F\_TOU | F-F | 9863 | 0.001 | 0.000 | 0.002 |
| F\_NIM | F\_STN | F-F | 10144 | 0.001 | 0.000 | 0.002 |
| F\_BUT | F\_TOU | F-F | 10024 | 0.001 | 0.000 | 0.002 |
| F\_MRH | F\_NIM | F-F | 9948 | 0.001 | 0.000 | 0.002 |
| F\_TOU | F\_STN | F-F | 10075 | 0.001 | 0.000 | 0.002 |
| F\_MER | F\_STN | F-F | 10173 | 0.001 | 0.000 | 0.002 |
| F\_TOU | F\_NIM | F-F | 10306 | 0.001 | 0.000 | 0.001 |
| F\_DER | F\_MER | F-F | 10048 | 0.001 | 0.000 | 0.002 |
| F\_MKH | F\_BUT | F-F | 10005 | 0.001 | 0.000 | 0.001 |
| F\_MRH | F\_MER | F-F | 9956 | 0.001 | 0.000 | 0.001 |
| F\_MER | F\_NIM | F-F | 10336 | 0.001 | 0.000 | 0.001 |
| F\_DER | F\_MIL | F-F | 9678 | 0.001 | 0.000 | 0.002 |
| F\_BUT | F\_MER | F-F | 10116 | 0.000 | 0.000 | 0.001 |
| F\_MKH | F\_DER | F-F | 9934 | 0.000 | 0.000 | 0.001 |
| F\_TOU | F\_MER | F-F | 10282 | 0.000 | 0.000 | 0.001 |
| F\_MKH | F\_MRH | F-F | 9852 | 0.000 | 0.000 | 0.001 |
| F\_MKH | F\_TOU | F-F | 10186 | 0.000 | 0.000 | 0.001 |
| F\_MRH | F\_DER | F-F | 9448 | 0.000 | 0.000 | 0.001 |
| F\_MKH | F\_MER | F-F | 10252 | 0.000 | 0.000 | 0.000 |
| F\_MKH | F\_NIM | F-F | 10259 | 0.000 | 0.000 | 0.000 |
| F\_DER | F\_BUT | F-F | 9667 | 0.000 | 0.000 | 0.001 |
| S\_BUT | S\_FRH | S-S | 8863 | 0.036 | 0.034 | 0.039 |
| S\_MIL | S\_BUT | S-S | 9226 | 0.024 | 0.022 | 0.026 |
| S\_MIL | S\_FRH | S-S | 9009 | 0.021 | 0.018 | 0.023 |
| S\_DER | S\_FRH | S-S | 9710 | 0.018 | 0.016 | 0.021 |
| S\_DER | S\_BUT | S-S | 9764 | 0.018 | 0.017 | 0.020 |
| S\_DER | S\_MIL | S-S | 9838 | 0.004 | 0.003 | 0.005 |
| F\_COL | L\_USR | F-L | 10252 | 0.010 | 0.009 | 0.011 |
| F\_MKH | L\_USR | F-L | 10074 | 0.010 | 0.009 | 0.011 |
| F\_FRH | L\_USR | F-L | 10243 | 0.010 | 0.008 | 0.011 |
| F\_NIM | L\_USR | F-L | 10151 | 0.009 | 0.008 | 0.010 |
| F\_STN | L\_USR | F-L | 9888 | 0.009 | 0.008 | 0.010 |
| F\_TOU | L\_USR | F-L | 10052 | 0.009 | 0.008 | 0.010 |
| F\_BUT | L\_USR | F-L | 9749 | 0.008 | 0.007 | 0.009 |
| F\_MRH | L\_USR | F-L | 9578 | 0.008 | 0.007 | 0.009 |
| F\_MIL | L\_USR | F-L | 9831 | 0.008 | 0.007 | 0.009 |
| F\_MER | L\_USR | F-L | 10159 | 0.008 | 0.007 | 0.009 |
| F\_DER | L\_USR | F-L | 9665 | 0.007 | 0.006 | 0.009 |
| F\_STN | W\_USR | F-W | 9545 | 0.161 | 0.156 | 0.167 |
| F\_MRH | W\_USR | F-W | 9052 | 0.160 | 0.155 | 0.166 |
| F\_NIM | W\_USR | F-W | 9920 | 0.157 | 0.151 | 0.162 |
| F\_BUT | W\_USR | F-W | 9370 | 0.157 | 0.151 | 0.162 |
| F\_MKH | W\_USR | F-W | 9744 | 0.156 | 0.151 | 0.161 |
| F\_TOU | W\_USR | F-W | 9800 | 0.156 | 0.150 | 0.161 |
| F\_DER | W\_USR | F-W | 9191 | 0.154 | 0.149 | 0.160 |
| F\_MIL | W\_USR | F-W | 9385 | 0.154 | 0.149 | 0.159 |
| F\_MER | W\_USR | F-W | 9914 | 0.154 | 0.149 | 0.159 |
| F\_FRH | W\_USR | F-W | 9946 | 0.145 | 0.140 | 0.151 |
| F\_COL | W\_USR | F-W | 9934 | 0.142 | 0.138 | 0.147 |
| F\_NIM | S\_BUT | F-S | 10190 | 0.043 | 0.041 | 0.045 |
| F\_STN | S\_BUT | F-S | 9874 | 0.043 | 0.040 | 0.045 |
| F\_MKH | S\_BUT | F-S | 10075 | 0.042 | 0.040 | 0.045 |
| F\_TOU | S\_BUT | F-S | 10045 | 0.042 | 0.040 | 0.044 |
| F\_MER | S\_BUT | F-S | 10185 | 0.040 | 0.038 | 0.042 |
| F\_MRH | S\_BUT | F-S | 9420 | 0.039 | 0.036 | 0.041 |
| F\_BUT | S\_BUT | F-S | 9720 | 0.039 | 0.037 | 0.041 |
| F\_DER | S\_BUT | F-S | 9561 | 0.037 | 0.035 | 0.040 |
| F\_MIL | S\_BUT | F-S | 9721 | 0.037 | 0.035 | 0.039 |
| F\_COL | S\_BUT | F-S | 10263 | 0.036 | 0.034 | 0.038 |
| F\_FRH | S\_BUT | F-S | 10190 | 0.034 | 0.032 | 0.036 |
| F\_NIM | S\_DER | F-S | 10662 | 0.028 | 0.026 | 0.030 |
| F\_NIM | S\_MIL | F-S | 10235 | 0.028 | 0.026 | 0.030 |
| F\_MKH | S\_MIL | F-S | 10111 | 0.028 | 0.026 | 0.030 |
| F\_MKH | S\_DER | F-S | 10546 | 0.028 | 0.026 | 0.030 |
| F\_TOU | S\_MIL | F-S | 10112 | 0.027 | 0.026 | 0.029 |
| F\_STN | S\_MIL | F-S | 9913 | 0.027 | 0.025 | 0.029 |
| F\_STN | S\_DER | F-S | 10413 | 0.027 | 0.025 | 0.029 |
| F\_TOU | S\_DER | F-S | 10544 | 0.027 | 0.025 | 0.028 |
| F\_MRH | S\_MIL | F-S | 9570 | 0.026 | 0.024 | 0.028 |
| F\_MER | S\_MIL | F-S | 10219 | 0.026 | 0.024 | 0.028 |
| F\_MER | S\_DER | F-S | 10620 | 0.026 | 0.024 | 0.028 |
| F\_BUT | S\_DER | F-S | 10302 | 0.025 | 0.023 | 0.027 |
| F\_BUT | S\_MIL | F-S | 9799 | 0.024 | 0.022 | 0.026 |
| F\_MRH | S\_DER | F-S | 10115 | 0.024 | 0.022 | 0.026 |
| F\_DER | S\_DER | F-S | 10189 | 0.023 | 0.021 | 0.024 |
| F\_MIL | S\_MIL | F-S | 9805 | 0.022 | 0.020 | 0.024 |
| F\_MIL | S\_DER | F-S | 10317 | 0.022 | 0.020 | 0.024 |
| F\_DER | S\_MIL | F-S | 9650 | 0.022 | 0.020 | 0.024 |
| F\_COL | S\_DER | F-S | 10702 | 0.021 | 0.020 | 0.023 |
| F\_COL | S\_MIL | F-S | 10297 | 0.021 | 0.020 | 0.023 |
| F\_FRH | S\_MIL | F-S | 10235 | 0.020 | 0.019 | 0.022 |
| F\_FRH | S\_DER | F-S | 10637 | 0.020 | 0.019 | 0.021 |
| F\_NIM | S\_FRH | F-S | 9790 | 0.013 | 0.011 | 0.015 |
| F\_STN | S\_FRH | F-S | 9382 | 0.012 | 0.010 | 0.014 |
| F\_MKH | S\_FRH | F-S | 9627 | 0.011 | 0.009 | 0.013 |
| F\_BUT | S\_FRH | F-S | 9264 | 0.010 | 0.008 | 0.012 |
| F\_TOU | S\_FRH | F-S | 9654 | 0.010 | 0.008 | 0.012 |
| F\_COL | S\_FRH | F-S | 9885 | 0.010 | 0.008 | 0.012 |
| F\_MER | S\_FRH | F-S | 9762 | 0.009 | 0.007 | 0.011 |
| F\_MRH | S\_FRH | F-S | 8928 | 0.008 | 0.006 | 0.010 |
| F\_DER | S\_FRH | F-S | 9050 | 0.008 | 0.006 | 0.010 |
| F\_MIL | S\_FRH | F-S | 9266 | 0.007 | 0.005 | 0.008 |
| F\_FRH | S\_FRH | F-S | 9754 | 0.003 | 0.002 | 0.005 |
| W\_USR | L\_USR | W-L | 9355 | 0.160 | 0.154 | 0.165 |
| S\_BUT | L\_USR | S-L | 9685 | 0.044 | 0.042 | 0.047 |
| S\_MIL | L\_USR | S-L | 9808 | 0.029 | 0.027 | 0.031 |
| S\_DER | L\_USR | S-L | 10307 | 0.029 | 0.027 | 0.031 |
| S\_FRH | L\_USR | S-L | 9252 | 0.016 | 0.014 | 0.019 |
| W\_USR | S\_BUT | W-S | 8812 | 0.159 | 0.154 | 0.165 |
| W\_USR | S\_FRH | W-S | 8288 | 0.158 | 0.152 | 0.163 |
| W\_USR | S\_MIL | W-S | 8985 | 0.148 | 0.143 | 0.153 |
| W\_USR | S\_DER | W-S | 9717 | 0.140 | 0.136 | 0.145 |

Figure S2: Pairwise Fst of individuals grouped by run/tributary. Fst-values denoted with \* have lower CI > 0

### 3 Estimates of effective population size

Estimates of effective population size Ne were determined using `LinkNe` (Hollenbeck, Portnoy, and Gold ⊕2016Hollenbeck, C M, D S Portnoy, and J R Gold. 2016. “A method for detecting recent changes in contemporary effective population size from linkage disequilibrium at linked and unlinked loci.” *Heredity* 117 (4): 207–16. https://doi.org/10.1038/hdy.2016.30.) an extension of the linkage disequilibrium method (LD) designed for data sets with loci of known linkage relationships. Loci that are linked, for example because they are on the same chromosome have elevate LD because of suppressed recombination which needs to be accounted for to not downwardly bias Ne estimates.

For Ne estimates data sets were generated per tributary/run group consisting of SNPs with a minor allele frequency > 0.01 and only one SNP per contig placed on a chromosome was retained.

**Map file with location of loci on chromosomes**

Create a file in standard `*.map` format indicating which chromosome each locus is on. Information can be used to generate Ne estimates using all loci, only loci on the same chromosome, only loci on different chromosomes; use only loci on Chromosomes (not unplaced scaffolds).

```
## Contig - chromosome matches all chromosome ----

# chromosome and chromosome number
chrom <- read_delim("data/REF/CHRIST2018/chrom.contigs", delim = " ") %>%
                    mutate(CHROM = str_trim(CHROM))

# position of contigs on chromosomes/scaffolds
contigs <- read_delim("data/REF/CHRIST2018/map_intervals.bed", delim = "\t",
                      col_names = c("CHROM", "chromStart", "chromEnd", "CONTIG")) %>%
            left_join(chrom)

loc <- as.data.frame(locNames(gen)) %>%
        rename(CONTIG = `locNames(gen)`) %>%
        left_join(contigs) %>%
        select(CHROMOSOME, CONTIG) %>%
        filter(!is.na(CHROMOSOME)) %>%
        arrange(CHROMOSOME, CONTIG)

write_delim(contigs, "data/POPGEN/contigs_all_chrom.txt", delim = "\t", col_names = FALSE)


## File per chromosome ----

for(p in unique(loc$CHROMOSOME)){

    print <- loc %>%
        filter(CHROMOSOME == p)

    path <- glue("data/POPGEN/contigs_chrom{p}.txt")

    write_delim(print, path, delim = "\t", col_names = FALSE)

}
```

Linked loci (e.g. because on the same chromosome) have elevated LD because of suppressed recombination which needs to be accounted for in order to not downwardly bias Ne estimates (Waples, Larson, and Waples ⊕2016Waples, R K, W A Larson, and R S Waples. 2016. “Estimating contemporary effective population size in non-model species using linkage disequilibrium across thousands of loci.” *Heredity* 117 (4): 233–40. https://doi.org/10.1038/hdy.2016.60.).

**Genepop file of diploid genotypes**

Use SNPs to calculate Ne (easier to deal with allele cut-offs). Filter loci with minor allele frequency < 0.01 then thin the data aset to retain only one SNP per contig (rad-tag).

```
# load modules for vcf filtering
module purge
module load GCC/6.4.0-2.28  OpenMPI/2.1.2
module load VCFtools/0.1.15-Perl-5.26.1

# set maf 0.01
vcftools --vcf data/POPGEN/ONC.haps.filtered.recode.vcf --out data/POPGEN/ONC.haps.maf01 --maf 0.01 --recode --recode-INFO-all

# thin data set
vcftools --vcf data/POPGEN/ONC.haps.maf01.recode.vcf --out data/POPGEN/ONC.haps.maf01.thin --thin 500 --recode --recode-INFO-all
```

Generate a single genepop file per population.

```
## READ IN GENETIC DATA ----

vcf <- read.vcfR("data/POPGEN/ONC.haps.maf01.thin.recode.vcf")

# convert to genind object
gen <- vcfR2genind(vcf)


## REMOVE LOCI NOT ON CHROMOSOMES ----

# chromosome and chromosome number
chrom <- read_delim("data/REF/CHRIST2018/chrom.contigs", delim = " ") %>%
                    mutate(CHROM = str_trim(CHROM))

# position of contigs on chromosomes/scaffolds
contigs <- read_delim("data/REF/CHRIST2018/map_intervals.bed", delim = "\t",
                      col_names = c("CHROM", "chromStart", "chromEnd", "CONTIG")) %>%
            left_join(chrom)

loc <- as.data.frame(locNames(gen)) %>%
        rename(LOCUS = `locNames(gen)`) %>%
        separate(LOCUS, into = c("tmp", "contig", "pos"), remove = FALSE) %>%
        mutate(CONTIG = glue("Contig_{contig}")) %>%
        left_join(contigs) %>%
        select(CHROMOSOME, CONTIG, LOCUS) %>%
        filter(!is.na(CHROMOSOME)) %>%
        arrange(CHROMOSOME, CONTIG)

gen <- gen[loc = loc$LOCUS]


## EXPORT GENEPOP ----

# define populations using defined stratification
setPop(gen) <- ~RUN_LOC

# create list of genind objects
grp_gen <- seppop(gen)

# write genepop for each population
for (p in names(grp_gen)){

    # genind for population
    tmp <- grp_gen[[p]]

    # convert to tidy data set
    tidy <- tidy_genomic_data(data = tmp, filename = NULL) %>%
        dplyr::select(1:6)

    path <- glue("data/POPGEN/ONC_SNP.maf01.{p}")

    header <- glue("CV chinook salmon thinned SNPs maf 0.01 {p}")

    # write genepop file
    write_genepop(data = tidy,
              filename = path,
              genepop.header = header)

}
```

Genomic distances (bp) were converted to recombination rates (cM) using the size of the female Chinook linkage map (3,118 cM, (Mckinney et al. ⊕2016Mckinney, G. J., L. W. Seeb, W. A. Larson, D. Gomez-Uchida, M. T. Limborg, M. S. O. Brieuc, M. V. Everett, K. A. Naish, R. K. Waples, and J. E. Seeb. 2016. “An integrated linkage map reveals candidate genes underlying adaptive variation in Chinook salmon (Oncorhynchus tshawytscha).” *Molecular Ecology Resources* 16 (3): 769–83. https://doi.org/10.1111/1755-0998.12479.)) and the length of the genome (2.4 Gbp, (Christensen et al. ⊕2018Christensen, Kris A., Jong S. Leong, Dionne Sakhrani, Carlo A. Biagi, David R. Minkley, Ruth E. Withler, Eric B. Rondeau, Ben F. Koop, and Robert H. Devlin. 2018. “Chinook salmon (Oncorhynchus tshawytscha) genome and transcriptome.” Edited by James P. Meador. *PLoS ONE* 13 (4): e0195461. https://doi.org/10.1371/journal.pone.0195461.)) resulting in an estimate of approx. 770kb being equivalent to one cM.

Format data frame with three columns (`locus`, `chromosome`, `position`); header is required.

```
map <- loc %>%
    left_join(contigs) %>%
    mutate(POSITION = chromStart/770000) %>%
    select(LOCUS, CHROMOSOME, POSITION) %>%
    rename(locus = LOCUS,
           chromosome = CHROMOSOME,
           position = POSITION)

write_delim(map, "data/POPGEN/ONC_SNP.thinned.maf01.map", delim = "\t")
```

**Run LinkNe**

```
# run out of folder with all files (example for one populations)
perl linkNe.pl -i ONC_SNP.maf01.F_BUT_genepop.gen -map ONC_SNP.thinned.maf01.map -o F_BUT.Ne
```

The largest c-values contain the the contemporary Ne estimates.

Estimates of effective population size for tributary/run groups using LinkNe.

| POP | NE | PARA\_95\_LOW | PARA\_95\_HIGH |
| --- | --- | --- | --- |
| F\_COL | 615 | 594 | 637 |
| F\_MIL | -6801 | -56952 | -3618 |
| F\_DER | -3939 | -9323 | -2497 |
| F\_BUT | 3802 | 2708 | 6378 |
| F\_FRH | 5172 | 3823 | 7988 |
| F\_NIM | 7432 | 5147 | 13351 |
| F\_MKH | 4106 | 3207 | 5704 |
| F\_STN | 2567 | 2040 | 3460 |
| F\_TOU | 6353 | 4383 | 11538 |
| F\_MRH | 1125 | 940 | 1399 |
| F\_MER | 11436 | 6768 | 36804 |
| L\_USR | 3114 | 2398 | 4435 |
| W\_USR | 174 | 171 | 177 |
| S\_MIL | 85 | 84 | 86 |
| S\_DER | 188 | 185 | 191 |
| S\_BUT | 561 | 529 | 597 |
| S\_FRH | -94 | -98 | -91 |

### 4 Assessment of genomic diversity

For all measures of genetic diversity comparisons were made for individuals grouped by run type within each tributary, with wild and hatchery individuals treated as separate groups even if they were sampled in the same tributary. Sample sizes among run/tributary groups are comparable with most in the range of 20 - 30, though only seven Feather River Spring individuals are present.

The following parameters are compared:

- Heterozygosity
  - observed heterozygosity Ho
  - gene diversity (expected heterozygosity) Hs
- Allele diversity
  - Allelic richness (allele counts and rarefied allele counts)
  - Evenness
- Sequence-based parameters
  - Watterson’s estimator (segregating sites): \(\Theta\_{S}\)
  - Nucleotide diversity (\(\pi\))

For each parameter, significant heterogeneity among groups was determined using a **Friedman’s test**, which implements a repeated block design (alleles observed per locus present in each sample). Next, significance of pairwise comparisons was tested using a **Wilcoxon signed rank test**; p-values were adjusted assuming a FDR of 0.05. The use of the signed rank test gives “direction” to the test of significance, due to the block design, i.e. pairwise comparisons show that loci will consistently be lower (higher) across individuals as opposed to a test that does not account for the fact that the same blocks/loci are being observed (e.g. a t-test).

Results of tests of significance for comparisons are presented as a heatmap indicating the test statistic of pairwise comparisons and printed p-values, significantly different comparisons (P < 0.05) is presented. Finally, the distribution of values across loci for individuals grouped by run or run/tributary are presented as boxplots and a table with summary statistics.

- Patterns of unique diversity
  - fixed loci
  - singletons
  - loci only variable in one group (private polymorphisms)
  - true private alleles

To compare whether identified loci are randomly distributed across chromosomes, null distributions were generated by shuffling chromosome designations across loci 1,000 times to determine whether the observed values fall outside the null distribution.

#### 4.1 Calculate measures of genetic diversity based on allele frequency and heterozygosity

```
# define groups to analyze ====
setPop(gen) <- ~RUN_LOC

gen_grp <- seppop(gen)

gen_grp[["ALL"]] <- gen

# calculate diversity stats ====
loc_stats <- list()

for (p in names(gen_grp)) {

locA <- locus_table(gen_grp[[p]], index = "shannon") %>%
  as.data.frame() %>%
  rownames_to_column("LOCUS")

locB <- locus_table(gen_grp[[p]], index = "simpson") %>%
  as.data.frame() %>%
  rownames_to_column("LOCUS")

temp <- left_join(locA, locB)

locC <- locus_table(gen_grp[[p]], index = "invsimpson") %>%
  as.data.frame() %>%
  rownames_to_column("LOCUS")

loc_stats[[p]] <- left_join(temp, locC)

}

loc_stats <- ldply(loc_stats, data.frame) %>%
  select(-Hexp) %>%
  rename(GRP = `.id`,
         SIMPSON_IDX = `X1.D`,
         N_ALLELES = allele,
         SHANNON_IDX = H,
         STODD_TAYLOR_IDX = G,
         EVENNESS = Evenness)

# calculate genetic diversity stats (heterozygosity-based) ====
loc_stats_2 <- list()

for (p in names(gen_grp)) {

dat <- hierfstat:::.genind2hierfstat(gen_grp[[p]])
stats <- basic.stats(dat)

loc_stats_2[[p]] <- stats$perloc %>%
  rownames_to_column("LOCUS")

}

# combine into single data frame ====
loc_stats_2 <- ldply(loc_stats_2, data.frame) %>%
  rename(GRP = `.id`)

loc_stats <- left_join(loc_stats, loc_stats_2) %>%
  select(GRP, LOCUS, N_ALLELES, EVENNESS, Ho, Hs, Ht, Fis, SHANNON_IDX, SIMPSON_IDX, STODD_TAYLOR_IDX) %>%
  filter(LOCUS != "mean")

loc_stats[is.na(loc_stats)] <- NA

write_delim(loc_stats, "results/gendiv.locstats", delim = "\t")
```

#### 4.2 observed heterozygosity

The observed heterozygosity (**Ho**) is measured as the proportion of heterozygote genotypes per locus (Nei ⊕1987Nei, Masatoshi. 1987. *Molecular Evolutionary Genetics*. https://doi.org/10.7312/nei-92038.).

**Compare distribution of observed heterozygosity within and among runs**

⊕Test of significant differences among runs using Friedman’s test.

```
## 
##  Friedman rank sum test
## 
## data:  Ho and GRP and LOCUS
## Friedman chi-squared = 3891.6, df = 16, p-value < 0.00000000000000022
```

Test for significant pairwise differences between tributaries within runs using Wilcoxon signed rank test.

```
# remove loci with Na values
rm <- het %>%
  filter(is.na(Ho))

het <- het %>%
  filter(!LOCUS %in% rm$LOCUS)

# groups to compare
comp <- as.character(unique(het$GRP))

# pairs of comparisons
pairs <- expand.grid(comp, comp) %>%
  filter(!Var1 == Var2) %>%
  rownames_to_column("PAIR") %>%
  split(.$PAIR) %>%
  purrr::map(function(x){
    x %>%
      select(-PAIR) %>%
      gather(key = temp, value = GRP, 1:2) %>%
      select(-temp)
  })

# empty data frame for results
results <- setNames(data.frame(matrix(ncol = 4, nrow = 0)), 
                    c("pop1", "pop2", "stat", "p.value")) %>%
  mutate(pop1 = as.character(pop1),
         pop2 = as.character(pop2),
         stat = as.numeric(stat),
         p.value = as.numeric(p.value))

n <- as.numeric(length(pairs))

# loop over pairs
for(i in 1:length(pairs)){

  p <- i

  pair <- pairs[[p]]$GRP

  temp <- het %>%
    filter(GRP %in% pair) %>%
    mutate(GRP = ordered(GRP, levels = pair),
         LOCUS = as.factor(LOCUS)) %>%
  droplevels()

  wilcox <- wilcoxsign_test(Ho ~ GRP | LOCUS,
                data = temp,
                zero.method = "Pratt")

  df <- data.frame("pop1" = pair[1],
                   "pop2" = pair[2],
                   "stat" = as.numeric(wilcox@statistic@teststatistic),
                   "p-value" = as.numeric(pvalue(wilcox)))

  results <- bind_rows(results, df)

}

results <- results %>%
  mutate(pop1 = ordered(pop1, levels = trib_runs),
         pop2 = ordered(pop2, levels = trib_runs))

write_delim(results, "results/obshet_run_trib.wilcox", delim = "\t")
```

Identify pairwise significantly different distribution between run/trib groups.

Results pairwise comparisons of observed heterozygosity among tributaries within runs using Wilcoxon signed rank test.

Compare distributions.

Distribution of observed heterozygosity of individuals grouped by tributary within each run.

Significantly different pairwise comparisons of observed heterozygosity across loci for individuals grouped by tributaries within runs.

| pop1 | pop2 | stat | p.value | p\_adj |
| --- | --- | --- | --- | --- |
| F\_FRH | W\_USR | 39.1048 | 0.0000 | 0.0000 |
| F\_COL | W\_USR | 38.6791 | 0.0000 | 0.0000 |
| F\_MER | W\_USR | 35.3954 | 0.0000 | 0.0000 |
| F\_NIM | W\_USR | 35.3437 | 0.0000 | 0.0000 |
| S\_DER | W\_USR | 35.3418 | 0.0000 | 0.0000 |
| F\_MKH | W\_USR | 35.1831 | 0.0000 | 0.0000 |
| F\_MIL | W\_USR | 34.3626 | 0.0000 | 0.0000 |
| F\_STN | W\_USR | 34.3477 | 0.0000 | 0.0000 |
| F\_TOU | W\_USR | 34.3074 | 0.0000 | 0.0000 |
| L\_USR | W\_USR | 33.1392 | 0.0000 | 0.0000 |
| F\_BUT | W\_USR | 32.4029 | 0.0000 | 0.0000 |
| F\_DER | W\_USR | 31.6226 | 0.0000 | 0.0000 |
| S\_MIL | W\_USR | 29.6530 | 0.0000 | 0.0000 |
| F\_MRH | W\_USR | 29.3136 | 0.0000 | 0.0000 |
| S\_BUT | W\_USR | 26.4736 | 0.0000 | 0.0000 |
| S\_FRH | W\_USR | 23.0823 | 0.0000 | 0.0000 |
| F\_FRH | S\_BUT | 16.1773 | 0.0000 | 0.0000 |
| F\_COL | S\_BUT | 14.3671 | 0.0000 | 0.0000 |
| F\_MIL | S\_BUT | 11.4354 | 0.0000 | 0.0000 |
| F\_MKH | S\_BUT | 11.3978 | 0.0000 | 0.0000 |
| F\_FRH | F\_BUT | 11.3174 | 0.0000 | 0.0000 |
| F\_NIM | S\_BUT | 11.2935 | 0.0000 | 0.0000 |
| F\_FRH | S\_MIL | 11.2336 | 0.0000 | 0.0000 |
| F\_FRH | S\_FRH | 10.8857 | 0.0000 | 0.0000 |
| F\_FRH | L\_USR | 10.6817 | 0.0000 | 0.0000 |
| F\_FRH | F\_MRH | 10.6776 | 0.0000 | 0.0000 |
| F\_MER | S\_BUT | 10.5184 | 0.0000 | 0.0000 |
| F\_FRH | F\_TOU | 10.3985 | 0.0000 | 0.0000 |
| F\_FRH | F\_MER | 10.0416 | 0.0000 | 0.0000 |
| F\_FRH | F\_DER | 9.5463 | 0.0000 | 0.0000 |
| F\_TOU | S\_BUT | 9.5314 | 0.0000 | 0.0000 |
| F\_COL | F\_BUT | 9.2579 | 0.0000 | 0.0000 |
| F\_FRH | S\_DER | 8.8955 | 0.0000 | 0.0000 |
| F\_FRH | F\_NIM | 8.7359 | 0.0000 | 0.0000 |
| F\_COL | L\_USR | 8.5759 | 0.0000 | 0.0000 |
| F\_FRH | F\_MKH | 8.5154 | 0.0000 | 0.0000 |
| F\_DER | S\_BUT | 8.4337 | 0.0000 | 0.0000 |
| F\_COL | S\_FRH | 8.3184 | 0.0000 | 0.0000 |
| F\_COL | S\_MIL | 8.0711 | 0.0000 | 0.0000 |
| S\_DER | S\_BUT | 7.6913 | 0.0000 | 0.0000 |
| F\_STN | S\_BUT | 7.6438 | 0.0000 | 0.0000 |
| F\_FRH | F\_STN | 7.5029 | 0.0000 | 0.0000 |
| F\_COL | S\_DER | 7.2397 | 0.0000 | 0.0000 |
| F\_COL | F\_MRH | 7.1334 | 0.0000 | 0.0000 |
| F\_MIL | F\_BUT | 7.0241 | 0.0000 | 0.0000 |
| F\_MKH | S\_FRH | 6.8937 | 0.0000 | 0.0000 |
| S\_MIL | S\_BUT | 6.7613 | 0.0000 | 0.0000 |
| F\_MRH | S\_BUT | 6.7527 | 0.0000 | 0.0000 |
| F\_NIM | F\_BUT | 6.4274 | 0.0000 | 0.0000 |
| F\_MKH | S\_MIL | 6.2274 | 0.0000 | 0.0000 |
| F\_BUT | S\_BUT | 6.0031 | 0.0000 | 0.0000 |
| F\_MKH | F\_BUT | 6.0024 | 0.0000 | 0.0000 |
| F\_MIL | L\_USR | 5.9055 | 0.0000 | 0.0000 |
| F\_NIM | S\_FRH | 5.8448 | 0.0000 | 0.0000 |
| F\_NIM | S\_MIL | 5.5633 | 0.0000 | 0.0000 |
| F\_COL | F\_DER | 5.5048 | 0.0000 | 0.0000 |
| F\_COL | F\_STN | 5.4600 | 0.0000 | 0.0000 |
| L\_USR | S\_BUT | 5.2146 | 0.0000 | 0.0000 |
| F\_COL | F\_TOU | 5.1889 | 0.0000 | 0.0000 |
| F\_NIM | L\_USR | 5.1206 | 0.0000 | 0.0000 |
| F\_FRH | F\_MIL | 5.0978 | 0.0000 | 0.0000 |
| F\_FRH | F\_COL | 5.0077 | 0.0000 | 0.0000 |
| F\_MER | S\_MIL | 4.9342 | 0.0000 | 0.0000 |
| F\_MER | S\_FRH | 4.8220 | 0.0000 | 0.0000 |
| F\_NIM | F\_MRH | 4.8102 | 0.0000 | 0.0000 |
| F\_MIL | S\_FRH | 4.8086 | 0.0000 | 0.0000 |
| F\_MIL | S\_DER | 4.7729 | 0.0000 | 0.0000 |
| F\_MKH | L\_USR | 4.6296 | 0.0000 | 0.0000 |
| F\_TOU | S\_FRH | 4.5094 | 0.0000 | 0.0000 |
| F\_MIL | F\_STN | 4.3115 | 0.0000 | 0.0000 |
| F\_NIM | S\_DER | 4.3009 | 0.0000 | 0.0000 |
| F\_COL | F\_MER | 4.1906 | 0.0000 | 0.0000 |
| F\_MER | F\_BUT | 4.1713 | 0.0000 | 0.0000 |
| F\_MKH | F\_MRH | 4.0767 | 0.0000 | 0.0000 |
| F\_MKH | F\_DER | 3.9318 | 0.0001 | 0.0002 |
| F\_TOU | S\_MIL | 3.9263 | 0.0001 | 0.0002 |
| F\_STN | S\_FRH | 3.9094 | 0.0001 | 0.0002 |
| F\_MIL | S\_MIL | 3.8983 | 0.0001 | 0.0002 |
| F\_MER | L\_USR | 3.8340 | 0.0001 | 0.0002 |
| F\_DER | F\_BUT | 3.8071 | 0.0001 | 0.0002 |
| F\_MKH | F\_MER | 3.7969 | 0.0001 | 0.0002 |
| F\_STN | S\_DER | 3.6471 | 0.0003 | 0.0005 |
| F\_COL | F\_MIL | 3.5339 | 0.0004 | 0.0007 |
| F\_MKH | S\_DER | 3.4436 | 0.0006 | 0.0010 |
| L\_USR | S\_FRH | 3.3424 | 0.0008 | 0.0013 |
| F\_NIM | F\_DER | 3.2529 | 0.0011 | 0.0017 |
| F\_MIL | F\_MRH | 3.2057 | 0.0013 | 0.0020 |
| S\_FRH | S\_BUT | 3.2008 | 0.0014 | 0.0022 |
| F\_DER | S\_MIL | 3.0998 | 0.0019 | 0.0029 |
| F\_MER | F\_MRH | 2.9651 | 0.0030 | 0.0045 |
| F\_MKH | F\_TOU | 2.9566 | 0.0031 | 0.0046 |
| F\_MER | S\_DER | 2.9367 | 0.0033 | 0.0049 |
| F\_TOU | F\_BUT | 2.9209 | 0.0035 | 0.0051 |
| F\_MIL | F\_TOU | 2.7212 | 0.0065 | 0.0094 |
| F\_DER | L\_USR | 2.6895 | 0.0072 | 0.0103 |
| S\_DER | S\_FRH | 2.6374 | 0.0084 | 0.0119 |
| F\_STN | S\_MIL | 2.6302 | 0.0085 | 0.0119 |
| F\_STN | L\_USR | 2.6288 | 0.0086 | 0.0119 |
| F\_NIM | F\_MER | 2.5304 | 0.0114 | 0.0157 |
| F\_MRH | S\_MIL | 2.5200 | 0.0117 | 0.0159 |
| F\_NIM | F\_STN | 2.4878 | 0.0129 | 0.0174 |
| F\_MKH | F\_NIM | 2.4789 | 0.0132 | 0.0176 |
| F\_TOU | F\_MRH | 2.3624 | 0.0182 | 0.0240 |
| F\_BUT | S\_FRH | 2.2552 | 0.0241 | 0.0315 |
| F\_BUT | L\_USR | 2.2400 | 0.0251 | 0.0325 |
| F\_STN | F\_MRH | 2.2356 | 0.0254 | 0.0326 |
| F\_TOU | L\_USR | 2.2095 | 0.0271 | 0.0344 |

Table S3: Mean, median, 25th, and 75th quartile of observed heterozygosity across loci for individuals grouped by run type & tributary.

| GRP | mean | median | Q1 | Q3 |
| --- | --- | --- | --- | --- |
| F\_COL | 0.1713 | 0.1000 | 0 | 0.3000 |
| F\_MIL | 0.1699 | 0.1000 | 0 | 0.2941 |
| F\_DER | 0.1674 | 0.0714 | 0 | 0.2857 |
| F\_BUT | 0.1641 | 0.1000 | 0 | 0.2778 |
| F\_FRH | 0.1741 | 0.1111 | 0 | 0.2963 |
| F\_NIM | 0.1688 | 0.1000 | 0 | 0.3000 |
| F\_MKH | 0.1689 | 0.0800 | 0 | 0.2917 |
| F\_STN | 0.1673 | 0.0909 | 0 | 0.2857 |
| F\_TOU | 0.1659 | 0.0833 | 0 | 0.2857 |
| F\_MRH | 0.1658 | 0.0769 | 0 | 0.2857 |
| F\_MER | 0.1670 | 0.1000 | 0 | 0.2903 |
| L\_USR | 0.1654 | 0.0952 | 0 | 0.2857 |
| W\_USR | 0.1272 | 0.0385 | 0 | 0.2308 |
| S\_MIL | 0.1651 | 0.0714 | 0 | 0.2857 |
| S\_DER | 0.1628 | 0.0870 | 0 | 0.2800 |
| S\_BUT | 0.1587 | 0.0588 | 0 | 0.2778 |
| S\_FRH | 0.1695 | 0.1429 | 0 | 0.2857 |

#### 4.3 Expected heterozygosity

Expected heterozygosity **Hs** is calculated as the proportion of heterozygous genotypes expected under Hardy-Weinberg equilibrium (Nei ⊕1987Nei, Masatoshi. 1987. *Molecular Evolutionary Genetics*. https://doi.org/10.7312/nei-92038.).

**Compare distribution of gene diversity within and among runs**

⊕Test of significant differences among runs using Friedman’s test.

```
## 
##  Friedman rank sum test
## 
## data:  Hs and GRP and LOCUS
## Friedman chi-squared = 3421.1, df = 16, p-value < 0.00000000000000022
```

Test for significant pairwise differences between tributaries within runs using Wilcoxon signed rank test.

Results pairwise comparisons of expected heterozygosity among tributaries within runs using Wilcoxon signed rank test.

Compare distributions across groups.

Distribution of expected heterozygosity per locus for individuals grouped by tributary within each run.

Significance of pairwise comparisons of expected heterozygosity for individuals grouped by tributaries within runs.

| pop1 | pop2 | stat | p.value | p\_adj |
| --- | --- | --- | --- | --- |
| F\_FRH | W\_USR | 39.2049 | 0.0000 | 0.0000 |
| S\_DER | W\_USR | 39.1176 | 0.0000 | 0.0000 |
| F\_COL | W\_USR | 38.1169 | 0.0000 | 0.0000 |
| F\_MIL | W\_USR | 36.5044 | 0.0000 | 0.0000 |
| F\_MER | W\_USR | 35.9299 | 0.0000 | 0.0000 |
| F\_NIM | W\_USR | 35.4636 | 0.0000 | 0.0000 |
| F\_TOU | W\_USR | 35.2467 | 0.0000 | 0.0000 |
| F\_STN | W\_USR | 35.0595 | 0.0000 | 0.0000 |
| F\_MKH | W\_USR | 35.0166 | 0.0000 | 0.0000 |
| L\_USR | W\_USR | 34.4389 | 0.0000 | 0.0000 |
| F\_BUT | W\_USR | 33.8632 | 0.0000 | 0.0000 |
| S\_MIL | W\_USR | 33.7289 | 0.0000 | 0.0000 |
| F\_DER | W\_USR | 33.4003 | 0.0000 | 0.0000 |
| F\_MRH | W\_USR | 32.8072 | 0.0000 | 0.0000 |
| S\_BUT | W\_USR | 29.6358 | 0.0000 | 0.0000 |
| S\_FRH | W\_USR | 25.8343 | 0.0000 | 0.0000 |
| F\_FRH | S\_BUT | 10.7653 | 0.0000 | 0.0000 |
| F\_MIL | S\_BUT | 9.3655 | 0.0000 | 0.0000 |
| S\_DER | S\_BUT | 9.2576 | 0.0000 | 0.0000 |
| F\_COL | S\_BUT | 9.1696 | 0.0000 | 0.0000 |
| F\_FRH | F\_MER | 8.8120 | 0.0000 | 0.0000 |
| F\_FRH | F\_NIM | 8.5196 | 0.0000 | 0.0000 |
| F\_FRH | F\_MKH | 8.2886 | 0.0000 | 0.0000 |
| F\_MIL | F\_BUT | 8.0117 | 0.0000 | 0.0000 |
| F\_FRH | F\_BUT | 7.8149 | 0.0000 | 0.0000 |
| S\_MIL | S\_BUT | 7.6811 | 0.0000 | 0.0000 |
| F\_FRH | L\_USR | 7.6379 | 0.0000 | 0.0000 |
| F\_MIL | L\_USR | 7.5210 | 0.0000 | 0.0000 |
| F\_FRH | F\_TOU | 7.4032 | 0.0000 | 0.0000 |
| F\_MIL | F\_STN | 7.1562 | 0.0000 | 0.0000 |
| F\_NIM | S\_BUT | 7.0782 | 0.0000 | 0.0000 |
| F\_MER | S\_BUT | 6.8788 | 0.0000 | 0.0000 |
| F\_MRH | S\_BUT | 6.6184 | 0.0000 | 0.0000 |
| F\_MKH | S\_BUT | 6.4468 | 0.0000 | 0.0000 |
| F\_DER | S\_BUT | 6.3022 | 0.0000 | 0.0000 |
| F\_TOU | S\_BUT | 6.2453 | 0.0000 | 0.0000 |
| F\_FRH | F\_DER | 6.2127 | 0.0000 | 0.0000 |
| F\_FRH | F\_STN | 6.0891 | 0.0000 | 0.0000 |
| F\_FRH | S\_FRH | 5.9649 | 0.0000 | 0.0000 |
| F\_COL | L\_USR | 5.6659 | 0.0000 | 0.0000 |
| F\_FRH | S\_MIL | 5.4279 | 0.0000 | 0.0000 |
| F\_FRH | F\_COL | 5.3574 | 0.0000 | 0.0000 |
| F\_MIL | F\_TOU | 5.0770 | 0.0000 | 0.0000 |
| F\_COL | F\_BUT | 4.7236 | 0.0000 | 0.0000 |
| F\_MIL | F\_MKH | 4.7030 | 0.0000 | 0.0000 |
| F\_MIL | F\_MER | 4.5026 | 0.0000 | 0.0000 |
| S\_DER | F\_TOU | 4.4197 | 0.0000 | 0.0000 |
| F\_MIL | F\_NIM | 4.3035 | 0.0000 | 0.0000 |
| S\_DER | S\_FRH | 4.2387 | 0.0000 | 0.0000 |
| F\_BUT | S\_BUT | 4.2346 | 0.0000 | 0.0000 |
| F\_FRH | F\_MRH | 4.1109 | 0.0000 | 0.0000 |
| S\_DER | F\_MKH | 4.0649 | 0.0000 | 0.0000 |
| F\_MIL | S\_FRH | 3.8499 | 0.0001 | 0.0003 |
| F\_STN | S\_BUT | 3.7820 | 0.0002 | 0.0005 |
| F\_MRH | F\_BUT | 3.7347 | 0.0002 | 0.0005 |
| S\_DER | F\_MER | 3.7025 | 0.0002 | 0.0005 |
| F\_COL | S\_FRH | 3.6908 | 0.0002 | 0.0005 |
| F\_DER | L\_USR | 3.5925 | 0.0003 | 0.0007 |
| S\_DER | F\_BUT | 3.5784 | 0.0003 | 0.0007 |
| S\_DER | F\_NIM | 3.4957 | 0.0005 | 0.0011 |
| F\_COL | F\_STN | 3.4423 | 0.0006 | 0.0013 |
| F\_DER | F\_BUT | 3.3728 | 0.0007 | 0.0015 |
| F\_MRH | F\_DER | 3.2570 | 0.0011 | 0.0024 |
| F\_MRH | L\_USR | 3.1089 | 0.0019 | 0.0040 |
| S\_DER | L\_USR | 2.9818 | 0.0029 | 0.0061 |
| L\_USR | S\_BUT | 2.8755 | 0.0040 | 0.0082 |
| F\_MKH | F\_MER | 2.7790 | 0.0055 | 0.0112 |
| F\_MKH | F\_NIM | 2.7182 | 0.0066 | 0.0132 |
| S\_MIL | F\_BUT | 2.7065 | 0.0068 | 0.0134 |
| F\_BUT | L\_USR | 2.6942 | 0.0071 | 0.0136 |
| F\_MIL | F\_DER | 2.6926 | 0.0071 | 0.0136 |
| F\_NIM | L\_USR | 2.6676 | 0.0076 | 0.0144 |
| S\_DER | F\_DER | 2.6247 | 0.0087 | 0.0162 |
| S\_MIL | L\_USR | 2.5775 | 0.0100 | 0.0184 |
| F\_MKH | L\_USR | 2.5647 | 0.0103 | 0.0187 |
| F\_MER | L\_USR | 2.5351 | 0.0112 | 0.0200 |
| F\_COL | F\_MER | 2.4938 | 0.0126 | 0.0221 |
| S\_DER | F\_STN | 2.4919 | 0.0127 | 0.0221 |
| F\_NIM | F\_BUT | 2.4844 | 0.0130 | 0.0224 |
| F\_MER | F\_BUT | 2.4674 | 0.0136 | 0.0231 |
| S\_FRH | S\_BUT | 2.3866 | 0.0170 | 0.0285 |
| F\_NIM | S\_FRH | 2.2779 | 0.0227 | 0.0376 |
| F\_MKH | F\_BUT | 2.2373 | 0.0253 | 0.0415 |
| S\_DER | S\_MIL | 2.1591 | 0.0308 | 0.0499 |

Table S4: Mean, median, 25th, and 75th quartile of observed heterozygosity per locus for individuals grouped by run type & tributary.

| GRP | mean | median | Q1 | Q3 |
| --- | --- | --- | --- | --- |
| F\_COL | 0.1693 | 0.0966 | 0 | 0.3034 |
| F\_MIL | 0.1714 | 0.1078 | 0 | 0.3125 |
| F\_DER | 0.1685 | 0.0769 | 0 | 0.3022 |
| F\_BUT | 0.1662 | 0.0974 | 0 | 0.2974 |
| F\_FRH | 0.1718 | 0.1068 | 0 | 0.3063 |
| F\_NIM | 0.1678 | 0.0966 | 0 | 0.3034 |
| F\_MKH | 0.1680 | 0.1032 | 0 | 0.3003 |
| F\_STN | 0.1670 | 0.0929 | 0 | 0.3030 |
| F\_TOU | 0.1676 | 0.0998 | 0 | 0.3014 |
| F\_MRH | 0.1695 | 0.0833 | 0 | 0.3077 |
| F\_MER | 0.1676 | 0.0966 | 0 | 0.2960 |
| L\_USR | 0.1668 | 0.0929 | 0 | 0.3000 |
| W\_USR | 0.1285 | 0.0385 | 0 | 0.2369 |
| S\_MIL | 0.1695 | 0.0769 | 0 | 0.3125 |
| S\_DER | 0.1699 | 0.1068 | 0 | 0.3030 |
| S\_BUT | 0.1642 | 0.1023 | 0 | 0.3070 |
| S\_FRH | 0.1695 | 0.1429 | 0 | 0.3571 |

#### 4.4 Comparison of allelic richness

The number of alleles observed per locus is not independent from sample size, especially for small samples sizes; with an increasing number of individuals sampled an increasing number of alleles is expected to be identified as rare alleles are more likely to be observed only for larger sample sizes. Differences in sample size can be adjusted for using rarefaction, i.e. by determining the lowest sample size and then subsampling x individuals from each group and counting alleles and using the mean number as the allele count.

**Compare differences in rarefied allele counts among tributaries within runs**

Calculate allele count per locus corrected for differences in sample size using rarefaction.

```
# by tributary within run
setPop(gen) <- ~RUN_LOC

# calculate allele counts using rarefaction
dat <- genind2hierfstat(gen)

df <- allelic.richness(dat,
                       diploid = TRUE)

df <- as.data.frame(df$Ar) %>%
  rownames_to_column("LOCUS")

colnames(df) <- c("LOCUS", trib_runs)

write_delim(df, "results/trib_run_rarefied.allelecount", delim = "\t")
```

⊕Test of significant differences among runs using Friedman’s test.

```
## 
##  Friedman rank sum test
## 
## data:  AR and pop and LOCUS
## Friedman chi-squared = 4236.7, df = 16, p-value < 0.00000000000000022
```

Test for significant pairwise differences between tributaries within runs using Wilcoxon signed rank test.

Significance of pairwise comparison of rarefied allele counts among runs using Wilcoxon signed rank test. Tile color indicates p-value, label is stat (indicates direction of difference); p-values were adjusted for multiple comparisons using Benjamini-Hochberg.

Compare distributions of values.

Distribution of rarefied allele counts per locus for individuals grouped by run type within tributaries.

Table S5 Significance of pairwise comparisons of allele counts per locus for individuals grouped by run type within tributaries.

| pop1 | pop2 | stat | p.value | p\_adj |
| --- | --- | --- | --- | --- |
| F\_MIL | L\_USR | 47.3157 | 0.0000 | 0.0000 |
| W\_USR | L\_USR | 46.7329 | 0.0000 | 0.0000 |
| F\_MRH | L\_USR | 46.4299 | 0.0000 | 0.0000 |
| F\_FRH | L\_USR | 44.2771 | 0.0000 | 0.0000 |
| F\_STN | L\_USR | 43.5133 | 0.0000 | 0.0000 |
| F\_MKH | L\_USR | 43.0949 | 0.0000 | 0.0000 |
| F\_TOU | L\_USR | 42.7212 | 0.0000 | 0.0000 |
| F\_COL | L\_USR | 42.6344 | 0.0000 | 0.0000 |
| F\_MER | L\_USR | 42.3246 | 0.0000 | 0.0000 |
| F\_NIM | L\_USR | 41.3909 | 0.0000 | 0.0000 |
| S\_FRH | L\_USR | 41.3626 | 0.0000 | 0.0000 |
| F\_BUT | L\_USR | 41.2526 | 0.0000 | 0.0000 |
| S\_MIL | L\_USR | 40.7391 | 0.0000 | 0.0000 |
| F\_DER | L\_USR | 40.5246 | 0.0000 | 0.0000 |
| S\_BUT | L\_USR | 35.6730 | 0.0000 | 0.0000 |
| S\_DER | L\_USR | 34.6154 | 0.0000 | 0.0000 |
| F\_MIL | S\_DER | 14.2654 | 0.0000 | 0.0000 |
| F\_FRH | S\_DER | 13.1611 | 0.0000 | 0.0000 |
| F\_MRH | S\_DER | 12.9707 | 0.0000 | 0.0000 |
| S\_BUT | S\_DER | 11.9349 | 0.0000 | 0.0000 |
| W\_USR | S\_DER | 11.6031 | 0.0000 | 0.0000 |
| F\_TOU | S\_DER | 10.5659 | 0.0000 | 0.0000 |
| S\_MIL | S\_DER | 10.5214 | 0.0000 | 0.0000 |
| F\_BUT | S\_DER | 10.3866 | 0.0000 | 0.0000 |
| F\_DER | S\_DER | 10.3305 | 0.0000 | 0.0000 |
| F\_MIL | F\_STN | 10.2502 | 0.0000 | 0.0000 |
| F\_STN | S\_DER | 10.2064 | 0.0000 | 0.0000 |
| F\_FRH | S\_FRH | 9.6039 | 0.0000 | 0.0000 |
| F\_MIL | F\_TOU | 9.4725 | 0.0000 | 0.0000 |
| F\_COL | S\_DER | 9.3000 | 0.0000 | 0.0000 |
| F\_MIL | F\_COL | 9.2263 | 0.0000 | 0.0000 |
| F\_MKH | S\_DER | 9.1481 | 0.0000 | 0.0000 |
| F\_FRH | F\_MER | 8.6395 | 0.0000 | 0.0000 |
| F\_MIL | S\_FRH | 8.4771 | 0.0000 | 0.0000 |
| F\_FRH | F\_NIM | 8.4581 | 0.0000 | 0.0000 |
| F\_MIL | F\_MKH | 8.1919 | 0.0000 | 0.0000 |
| S\_BUT | S\_FRH | 8.0404 | 0.0000 | 0.0000 |
| F\_NIM | S\_DER | 7.3802 | 0.0000 | 0.0000 |
| F\_MER | S\_DER | 7.1595 | 0.0000 | 0.0000 |
| F\_FRH | F\_MKH | 7.1041 | 0.0000 | 0.0000 |
| F\_MIL | F\_NIM | 6.9014 | 0.0000 | 0.0000 |
| S\_BUT | F\_NIM | 6.8211 | 0.0000 | 0.0000 |
| F\_FRH | F\_STN | 6.8165 | 0.0000 | 0.0000 |
| F\_MRH | S\_FRH | 6.7512 | 0.0000 | 0.0000 |
| S\_BUT | F\_MER | 6.5489 | 0.0000 | 0.0000 |
| S\_BUT | S\_MIL | 6.5482 | 0.0000 | 0.0000 |
| F\_FRH | F\_COL | 6.5399 | 0.0000 | 0.0000 |
| S\_BUT | F\_DER | 6.4537 | 0.0000 | 0.0000 |
| F\_FRH | F\_TOU | 6.4394 | 0.0000 | 0.0000 |
| F\_BUT | S\_FRH | 6.3824 | 0.0000 | 0.0000 |
| S\_BUT | F\_BUT | 6.0927 | 0.0000 | 0.0000 |
| F\_MIL | F\_MER | 6.0609 | 0.0000 | 0.0000 |
| F\_MIL | F\_MRH | 6.0357 | 0.0000 | 0.0000 |
| F\_MIL | S\_MIL | 5.6104 | 0.0000 | 0.0000 |
| F\_DER | S\_FRH | 5.3952 | 0.0000 | 0.0000 |
| F\_DER | F\_NIM | 5.3912 | 0.0000 | 0.0000 |
| F\_BUT | F\_NIM | 5.2145 | 0.0000 | 0.0000 |
| S\_FRH | S\_DER | 5.0936 | 0.0000 | 0.0000 |
| F\_DER | F\_COL | 4.7337 | 0.0000 | 0.0000 |
| S\_BUT | F\_COL | 4.6233 | 0.0000 | 0.0000 |
| F\_DER | F\_STN | 4.5255 | 0.0000 | 0.0000 |
| S\_BUT | F\_MKH | 4.4261 | 0.0000 | 0.0000 |
| W\_USR | F\_MKH | 4.4182 | 0.0000 | 0.0000 |
| W\_USR | F\_COL | 4.3173 | 0.0000 | 0.0000 |
| F\_DER | F\_MKH | 4.1916 | 0.0000 | 0.0000 |
| F\_MRH | F\_NIM | 4.1878 | 0.0000 | 0.0000 |
| F\_DER | F\_MER | 4.1412 | 0.0000 | 0.0000 |
| S\_BUT | F\_STN | 4.0940 | 0.0000 | 0.0000 |
| F\_NIM | S\_FRH | 4.0102 | 0.0001 | 0.0002 |
| W\_USR | F\_TOU | 3.8267 | 0.0001 | 0.0002 |
| W\_USR | F\_STN | 3.8196 | 0.0001 | 0.0002 |
| F\_MIL | F\_BUT | 3.7908 | 0.0002 | 0.0004 |
| F\_BUT | F\_MER | 3.7600 | 0.0002 | 0.0004 |
| F\_DER | F\_TOU | 3.6623 | 0.0002 | 0.0004 |
| W\_USR | S\_FRH | 3.6612 | 0.0003 | 0.0005 |
| F\_MRH | F\_MER | 3.6446 | 0.0003 | 0.0005 |
| F\_MRH | F\_STN | 3.5798 | 0.0003 | 0.0005 |
| F\_BUT | F\_STN | 3.5178 | 0.0004 | 0.0007 |
| S\_BUT | F\_TOU | 3.4181 | 0.0006 | 0.0010 |
| F\_FRH | F\_MRH | 3.3930 | 0.0007 | 0.0012 |
| F\_BUT | F\_MKH | 3.3042 | 0.0010 | 0.0017 |
| F\_BUT | F\_TOU | 3.1003 | 0.0019 | 0.0032 |
| S\_MIL | S\_FRH | 3.0946 | 0.0020 | 0.0033 |
| F\_DER | S\_MIL | 3.0465 | 0.0023 | 0.0037 |
| F\_BUT | F\_COL | 2.9286 | 0.0034 | 0.0054 |
| F\_TOU | S\_FRH | 2.9152 | 0.0036 | 0.0057 |
| F\_COL | F\_STN | 2.8341 | 0.0046 | 0.0072 |
| F\_COL | F\_TOU | 2.8100 | 0.0050 | 0.0077 |
| S\_BUT | F\_FRH | 2.7887 | 0.0053 | 0.0081 |
| F\_STN | S\_FRH | 2.7431 | 0.0061 | 0.0092 |
| F\_DER | F\_BUT | 2.7092 | 0.0067 | 0.0100 |
| W\_USR | F\_NIM | 2.7077 | 0.0068 | 0.0101 |
| F\_FRH | W\_USR | 2.6747 | 0.0075 | 0.0110 |
| F\_COL | S\_FRH | 2.4104 | 0.0159 | 0.0230 |
| F\_NIM | F\_MER | 2.3341 | 0.0196 | 0.0281 |
| S\_MIL | F\_NIM | 2.2804 | 0.0226 | 0.0320 |
| F\_FRH | S\_MIL | 2.2314 | 0.0257 | 0.0360 |
| F\_MKH | S\_FRH | 2.1175 | 0.0342 | 0.0475 |
| S\_MIL | F\_MER | 2.0975 | 0.0360 | 0.0495 |

Table S6: Mean, median, 25th, and 75th quartile of allelic richness across loci for individuals grouped by run type & tributary.

| pop | mean | median | Q1 | Q3 |
| --- | --- | --- | --- | --- |
| F\_COL | 1.51 | 1.45 | 1 | 1.91 |
| F\_MIL | 1.52 | 1.47 | 1 | 1.92 |
| F\_DER | 1.51 | 1.42 | 1 | 1.93 |
| F\_BUT | 1.51 | 1.38 | 1 | 1.93 |
| F\_FRH | 1.52 | 1.48 | 1 | 1.93 |
| F\_NIM | 1.51 | 1.44 | 1 | 1.91 |
| F\_MKH | 1.51 | 1.44 | 1 | 1.91 |
| F\_STN | 1.51 | 1.43 | 1 | 1.90 |
| F\_TOU | 1.51 | 1.43 | 1 | 1.91 |
| F\_MRH | 1.52 | 1.43 | 1 | 1.91 |
| F\_MER | 1.51 | 1.42 | 1 | 1.91 |
| L\_USR | 1.36 | 1.19 | 1 | 1.80 |
| W\_USR | 1.52 | 1.47 | 1 | 1.91 |
| S\_MIL | 1.51 | 1.38 | 1 | 1.93 |
| S\_DER | 1.48 | 1.46 | 1 | 1.92 |
| S\_BUT | 1.51 | 1.71 | 1 | 1.99 |
| S\_FRH | 1.50 | 1.42 | 1 | 1.91 |

#### 4.5 Evenness of allelic diversity

The **Evenness** of allelic diversity at a given locus is calculated as the ratio of the number of abundant genotypes to the number of rarer genotypes calculated using the ratio of `Stoddart & Tayolor index` (diversity index weighted for more abundant alleles) & `Shannon-Wiener index` (diversity index weighted for more rare alleles).

**Compare differences in evenness of allelic diversity among tributaries within runs**

⊕Test of significant differences among runs using Friedman’s test.

```
## 
##  Friedman rank sum test
## 
## data:  EVENNESS and GRP and LOCUS
## Friedman chi-squared = 791.32, df = 16, p-value < 0.00000000000000022
```

Test for significant pairwise differences between runs using Wilcoxon signed rank test.

Significance of pairwise comparisons of evenness of allelic diversity per locus for individuals grouped by run type within tributaries using using Wilcoxon signed rank test. Tile color indicates stat (indicates direction of difference); p-values were adjusted for multiple comparisons using Benjamini-Hochberg.

Significance of pairwise comparisons of evenness of allelic diversity across loci for individuals grouped by run type within tributaries.

| pop1 | pop2 | stat | p.value | p\_adj |
| --- | --- | --- | --- | --- |
| F\_FRH | W\_USR | 21.10 | 0.0000 | 0.0000 |
| S\_FRH | W\_USR | 20.80 | 0.0000 | 0.0000 |
| F\_MIL | W\_USR | 20.36 | 0.0000 | 0.0000 |
| S\_MIL | W\_USR | 20.31 | 0.0000 | 0.0000 |
| F\_COL | W\_USR | 20.18 | 0.0000 | 0.0000 |
| S\_DER | W\_USR | 19.95 | 0.0000 | 0.0000 |
| F\_DER | W\_USR | 19.24 | 0.0000 | 0.0000 |
| F\_MER | W\_USR | 19.24 | 0.0000 | 0.0000 |
| F\_MRH | W\_USR | 19.22 | 0.0000 | 0.0000 |
| F\_NIM | W\_USR | 19.12 | 0.0000 | 0.0000 |
| F\_STN | W\_USR | 19.02 | 0.0000 | 0.0000 |
| F\_MKH | W\_USR | 18.97 | 0.0000 | 0.0000 |
| F\_BUT | W\_USR | 18.78 | 0.0000 | 0.0000 |
| F\_TOU | W\_USR | 18.78 | 0.0000 | 0.0000 |
| L\_USR | W\_USR | 18.59 | 0.0000 | 0.0000 |
| S\_BUT | W\_USR | 17.83 | 0.0000 | 0.0000 |
| F\_FRH | F\_TOU | 4.65 | 0.0000 | 0.0000 |
| S\_FRH | S\_BUT | 4.61 | 0.0000 | 0.0000 |
| F\_FRH | S\_BUT | 4.55 | 0.0000 | 0.0000 |
| F\_FRH | L\_USR | 4.34 | 0.0000 | 0.0000 |
| S\_FRH | F\_TOU | 4.24 | 0.0000 | 0.0000 |
| F\_FRH | F\_BUT | 4.20 | 0.0000 | 0.0000 |
| S\_FRH | L\_USR | 4.19 | 0.0000 | 0.0000 |
| F\_FRH | F\_MKH | 4.04 | 0.0001 | 0.0005 |
| F\_FRH | F\_NIM | 3.99 | 0.0001 | 0.0005 |
| F\_FRH | F\_MER | 3.91 | 0.0001 | 0.0005 |
| S\_FRH | F\_NIM | 3.89 | 0.0001 | 0.0005 |
| S\_FRH | F\_BUT | 3.89 | 0.0001 | 0.0005 |
| S\_FRH | F\_MKH | 3.82 | 0.0001 | 0.0005 |
| S\_FRH | F\_MER | 3.75 | 0.0002 | 0.0009 |
| S\_MIL | S\_BUT | 3.63 | 0.0003 | 0.0013 |
| F\_MIL | S\_BUT | 3.55 | 0.0004 | 0.0016 |
| F\_MIL | F\_BUT | 3.53 | 0.0004 | 0.0016 |
| F\_FRH | F\_STN | 3.50 | 0.0005 | 0.0019 |
| F\_MIL | F\_TOU | 3.47 | 0.0005 | 0.0019 |
| S\_FRH | F\_STN | 3.41 | 0.0007 | 0.0026 |
| S\_FRH | F\_DER | 3.40 | 0.0007 | 0.0026 |
| S\_FRH | S\_DER | 3.34 | 0.0008 | 0.0029 |
| F\_FRH | F\_DER | 3.34 | 0.0009 | 0.0031 |
| S\_FRH | F\_COL | 3.31 | 0.0009 | 0.0031 |
| F\_FRH | F\_COL | 3.30 | 0.0010 | 0.0033 |
| F\_MIL | L\_USR | 3.12 | 0.0018 | 0.0058 |
| F\_MIL | F\_NIM | 3.10 | 0.0020 | 0.0063 |
| F\_FRH | S\_DER | 2.96 | 0.0031 | 0.0096 |
| F\_MIL | F\_MKH | 2.76 | 0.0058 | 0.0175 |
| F\_COL | S\_BUT | 2.72 | 0.0065 | 0.0192 |
| F\_MIL | F\_DER | 2.70 | 0.0068 | 0.0197 |
| F\_MRH | S\_BUT | 2.62 | 0.0088 | 0.0249 |
| F\_MIL | F\_MER | 2.55 | 0.0107 | 0.0297 |
| S\_FRH | F\_MRH | 2.52 | 0.0116 | 0.0316 |
| S\_DER | S\_BUT | 2.51 | 0.0121 | 0.0323 |
| F\_FRH | F\_MRH | 2.39 | 0.0171 | 0.0447 |
| S\_MIL | L\_USR | 2.34 | 0.0191 | 0.0490 |

Compare distributions of values.

Distribution of evenness of allelic diversity per locus for individuals grouped by run type within tributaries.

Table S7: Mean, median, 25th, and 75th quartile of evenness of allelic richness per locus for individuals grouped by run type & tributary.

| GRP | mean | median | Q1 | Q3 |
| --- | --- | --- | --- | --- |
| F\_COL | 0.7645 | 0.7720 | 0.6506 | 0.8990 |
| F\_MIL | 0.7681 | 0.7747 | 0.6461 | 0.8990 |
| F\_DER | 0.7631 | 0.7720 | 0.6257 | 0.8990 |
| F\_BUT | 0.7617 | 0.7747 | 0.6336 | 0.8990 |
| F\_FRH | 0.7697 | 0.7823 | 0.6478 | 0.8990 |
| F\_NIM | 0.7631 | 0.7720 | 0.6321 | 0.8990 |
| F\_MKH | 0.7632 | 0.7703 | 0.6397 | 0.8990 |
| F\_STN | 0.7631 | 0.7786 | 0.6336 | 0.8990 |
| F\_TOU | 0.7620 | 0.7703 | 0.6353 | 0.8988 |
| F\_MRH | 0.7648 | 0.7720 | 0.6397 | 0.8990 |
| F\_MER | 0.7634 | 0.7720 | 0.6321 | 0.8984 |
| L\_USR | 0.7609 | 0.7769 | 0.6397 | 0.8990 |
| W\_USR | 0.6925 | 0.6853 | 0.5040 | 0.8694 |
| S\_MIL | 0.7663 | 0.7734 | 0.6397 | 0.8990 |
| S\_DER | 0.7633 | 0.7684 | 0.6397 | 0.8990 |
| S\_BUT | 0.7578 | 0.7769 | 0.6223 | 0.9085 |
| S\_FRH | 0.7715 | 0.7534 | 0.6397 | 0.9240 |

#### 4.6 Nucleotide diversity \(\pi\) (pairwise differences)

\(\theta\_{T}\) (Nei ⊕1987Nei, Masatoshi. 1987. *Molecular Evolutionary Genetics*. https://doi.org/10.7312/nei-92038.), the **nucleotide diversity** (\(\pi\)) is calculated as the sum of the number of differences between pairs of haplotypes of a given locus, divided by the number of comparisons made. This parameter is biased towards alleles segregating at intermediate rates and will underestimate genetic diversity due to rare alleles.

Create a tidy data set of variant call information (position, alternate alleles) from filtered VCF file used to create haplotypes. Create a data frame of haplotype sequences in the data set(s) using reference contig sequences, VCF information, and `codes.out` file from `rad_haplotyper`.

```
# Read in the reference sequence
fa <- seqinr::read.fasta("data/REF/CHRIST2018/reference.fasta")

# create tibble with locus name and sequence
ref <- tibble::tibble(name = seqinr::getName(fa), seq = toupper(unlist(seqinr::getSequence(fa, as.string = TRUE))))

# Read the haplotype VCF file and put it in tidy format
vcf_tidy <- read.vcfR("data/POPGEN/ONC.haps.filtered.recode.vcf") %>%
    vcfR2tidy()

# Get the data frame with the positions
pos_tbl <- vcf_tidy$fix %>%
  filter(ALT %in% c("A", "T", "C", "G")) %>%
  unite(LOCUS, CHROM, POS, sep = "_", remove = FALSE)


# Make a data frame with all of the haplotype sequences from the 'codes.out' file from rad_haplotyper
hap_index <- read_tsv("data/HAPLOTYPING/codes.ONC.gen", col_names = FALSE) %>%
    filter(X1 %in% locNames(gen_obj)) %>%
    split(.$X1) %>%
    purrr::map(function(x) {
      codes <- str_split(x$X2, ",") %>%
        unlist()

      code_tbl <- tibble(locus = x$X1, code = codes) %>%
        separate(code, c("hap", "code"), sep = ":") %>%
        left_join(ref, by = c("locus" = "name"))

      pos <- pos_tbl %>%
        filter(CHROM == code_tbl$locus[1]) %>%
        pull(POS)

      for (i in 1:nrow(code_tbl)) {

        alleles <- str_split(code_tbl$hap[i], "") %>%
          unlist()

        replace <- tibble(pos = pos, allele = alleles)

        for (j in 1:nrow(replace)) {
          str_sub(code_tbl$seq[i], replace$pos[j], 1) <- replace$allele[j]
        }

      }

      code_tbl

    }) %>%
    bind_rows()
```

Calculate nucleotide diversity for individuals grouped by run type within each tributary.

```
# set pops
setPop(gen) <- ~RUN_LOC

# create tidy data set
gen_tidy <- gen %>%
  genind2df(oneColPerAll = TRUE) %>%
  rownames_to_column(var = "ind") %>%
  gather(allele, code, -pop, -ind) %>%
  separate(allele, c("locus", "allele"), sep = "\\.") %>%
  filter(pop %in% trib_runs) %>%
  droplevels()

# Calculate haplotype related stats
hap_div_tbl <- locNames(gen) %>%
  purrr::map(function(y) {

    # for each locus
    gen_tidy %>%
      filter(locus == y) %>%
      filter(!code == "NA") %>%
      left_join(hap_index, by = c("code", "locus")) %>%
      as.tibble() %>%
      arrange(factor(pop, levels = trib_runs)) %>%

      # for each population per locus
      split(.$pop) %>%
      purrr::map(function(x) {
        y <- do.call(rbind, strsplit(x$seq, ""))
        rownames(y) <- paste(x$ind, x$allele, sep = ".")

        # create DNAbin object for locus
        dna_bin <- as.DNAbin(y)                            

        # calculate Tajima's D and p-values
        taj_test <- tajima.test(dna_bin)

        # create table with results
        tibble(locus = x$locus[1],
               pop = x$pop[1],
               nuc_div = nuc.div(dna_bin)
               
      }) %>%
      bind_rows()
  }) %>%
    bind_rows()

write_delim(hap_div_tbl, "results/trib_run.hapdiv", delim = "\t")
```

⊕Test of significant differences among runs using Friedman’s test. To get an unreplicated complete block design only loci variable in all locations were used for test of heterogeneity among groups.

```
## 
##  Friedman rank sum test
## 
## data:  nuc_div and pop and locus
## Friedman chi-squared = 3466, df = 16, p-value < 0.00000000000000022
```

Test for significance of pairwise comparisons using Wilcoxon signed rank test.

Pairwise comparison of levels of significant differences among individuals grouped by tributary and run type.

Compare distribution of values.

Distribution of nucleotide diversity across all loci for individuals grouped by tributary and runs

Significance of pairwise comparisons of nucleotide diversity among runs within tributaries.

| pop1 | pop2 | stat | p.value | p\_adj |
| --- | --- | --- | --- | --- |
| F\_FRH | W\_USR | 39.1632 | 0.0000 | 0.0000 |
| S\_DER | W\_USR | 38.7345 | 0.0000 | 0.0000 |
| F\_COL | W\_USR | 38.0562 | 0.0000 | 0.0000 |
| F\_MIL | W\_USR | 36.2644 | 0.0000 | 0.0000 |
| F\_MER | W\_USR | 35.8979 | 0.0000 | 0.0000 |
| F\_NIM | W\_USR | 35.4991 | 0.0000 | 0.0000 |
| F\_TOU | W\_USR | 35.3305 | 0.0000 | 0.0000 |
| F\_MKH | W\_USR | 34.9710 | 0.0000 | 0.0000 |
| F\_STN | W\_USR | 34.8902 | 0.0000 | 0.0000 |
| L\_USR | W\_USR | 34.3191 | 0.0000 | 0.0000 |
| F\_BUT | W\_USR | 33.7190 | 0.0000 | 0.0000 |
| S\_MIL | W\_USR | 33.4761 | 0.0000 | 0.0000 |
| F\_DER | W\_USR | 33.3037 | 0.0000 | 0.0000 |
| F\_MRH | W\_USR | 32.6041 | 0.0000 | 0.0000 |
| S\_BUT | W\_USR | 29.3360 | 0.0000 | 0.0000 |
| S\_FRH | W\_USR | 25.7697 | 0.0000 | 0.0000 |
| F\_FRH | S\_BUT | 10.6609 | 0.0000 | 0.0000 |
| F\_MIL | S\_BUT | 9.5362 | 0.0000 | 0.0000 |
| F\_COL | S\_BUT | 9.3455 | 0.0000 | 0.0000 |
| S\_DER | S\_BUT | 9.2383 | 0.0000 | 0.0000 |
| F\_FRH | F\_MER | 8.4649 | 0.0000 | 0.0000 |
| F\_FRH | F\_MKH | 8.0066 | 0.0000 | 0.0000 |
| F\_FRH | F\_NIM | 7.8470 | 0.0000 | 0.0000 |
| F\_MIL | F\_BUT | 7.8298 | 0.0000 | 0.0000 |
| F\_FRH | F\_BUT | 7.8249 | 0.0000 | 0.0000 |
| S\_MIL | S\_BUT | 7.5688 | 0.0000 | 0.0000 |
| F\_FRH | L\_USR | 7.5485 | 0.0000 | 0.0000 |
| F\_FRH | F\_TOU | 7.3375 | 0.0000 | 0.0000 |
| F\_NIM | S\_BUT | 7.2899 | 0.0000 | 0.0000 |
| F\_MIL | L\_USR | 7.1077 | 0.0000 | 0.0000 |
| F\_MER | S\_BUT | 7.0675 | 0.0000 | 0.0000 |
| F\_MIL | F\_STN | 6.9026 | 0.0000 | 0.0000 |
| F\_MKH | S\_BUT | 6.7400 | 0.0000 | 0.0000 |
| F\_FRH | F\_DER | 6.4714 | 0.0000 | 0.0000 |
| F\_MRH | S\_BUT | 6.4262 | 0.0000 | 0.0000 |
| F\_TOU | S\_BUT | 6.4191 | 0.0000 | 0.0000 |
| F\_FRH | S\_FRH | 6.3863 | 0.0000 | 0.0000 |
| F\_DER | S\_BUT | 6.1614 | 0.0000 | 0.0000 |
| F\_FRH | F\_STN | 6.1338 | 0.0000 | 0.0000 |
| F\_COL | L\_USR | 5.6465 | 0.0000 | 0.0000 |
| F\_FRH | S\_MIL | 5.6304 | 0.0000 | 0.0000 |
| F\_FRH | F\_COL | 5.0746 | 0.0000 | 0.0000 |
| F\_MIL | F\_TOU | 4.9915 | 0.0000 | 0.0000 |
| F\_COL | F\_BUT | 4.9600 | 0.0000 | 0.0000 |
| F\_FRH | F\_MRH | 4.4911 | 0.0000 | 0.0000 |
| F\_MIL | F\_MKH | 4.4521 | 0.0000 | 0.0000 |
| F\_COL | S\_FRH | 4.4422 | 0.0000 | 0.0000 |
| S\_DER | S\_FRH | 4.3575 | 0.0000 | 0.0000 |
| F\_MIL | S\_FRH | 4.3237 | 0.0000 | 0.0000 |
| F\_MIL | F\_MER | 4.2719 | 0.0000 | 0.0000 |
| F\_STN | S\_BUT | 4.2075 | 0.0000 | 0.0000 |
| F\_BUT | S\_BUT | 4.1843 | 0.0000 | 0.0000 |
| S\_DER | F\_TOU | 3.9545 | 0.0001 | 0.0003 |
| F\_MIL | F\_NIM | 3.8026 | 0.0001 | 0.0003 |
| F\_COL | F\_STN | 3.4890 | 0.0005 | 0.0012 |
| F\_MRH | F\_BUT | 3.4789 | 0.0005 | 0.0012 |
| S\_DER | F\_BUT | 3.3540 | 0.0008 | 0.0019 |
| S\_DER | F\_MKH | 3.2867 | 0.0010 | 0.0023 |
| F\_DER | F\_BUT | 3.2540 | 0.0011 | 0.0025 |
| F\_DER | L\_USR | 3.2284 | 0.0012 | 0.0027 |
| L\_USR | S\_BUT | 3.1632 | 0.0016 | 0.0035 |
| S\_DER | F\_MER | 3.1512 | 0.0016 | 0.0035 |
| F\_NIM | F\_BUT | 2.8425 | 0.0045 | 0.0097 |
| S\_DER | F\_NIM | 2.8292 | 0.0047 | 0.0100 |
| S\_DER | L\_USR | 2.7650 | 0.0057 | 0.0119 |
| F\_NIM | S\_FRH | 2.7593 | 0.0058 | 0.0120 |
| F\_MIL | F\_DER | 2.7508 | 0.0059 | 0.0120 |
| F\_NIM | L\_USR | 2.7437 | 0.0061 | 0.0122 |
| F\_MKH | L\_USR | 2.7355 | 0.0062 | 0.0122 |
| F\_MRH | F\_DER | 2.6703 | 0.0076 | 0.0148 |
| F\_MER | F\_BUT | 2.6484 | 0.0081 | 0.0155 |
| F\_MKH | F\_MER | 2.5498 | 0.0108 | 0.0203 |
| F\_COL | F\_MER | 2.5448 | 0.0109 | 0.0203 |
| S\_DER | F\_DER | 2.4792 | 0.0132 | 0.0243 |
| F\_MRH | L\_USR | 2.4721 | 0.0134 | 0.0243 |
| F\_MKH | F\_NIM | 2.4247 | 0.0153 | 0.0274 |
| F\_MER | L\_USR | 2.3802 | 0.0173 | 0.0306 |
| S\_DER | S\_MIL | 2.3737 | 0.0176 | 0.0307 |
| F\_TOU | S\_FRH | 2.3175 | 0.0205 | 0.0350 |
| F\_COL | S\_MIL | 2.3160 | 0.0206 | 0.0350 |
| S\_FRH | S\_BUT | 2.3072 | 0.0210 | 0.0353 |
| S\_DER | F\_STN | 2.3011 | 0.0214 | 0.0355 |
| F\_MKH | F\_BUT | 2.2460 | 0.0247 | 0.0403 |
| F\_MER | S\_FRH | 2.2424 | 0.0249 | 0.0403 |
| S\_MIL | L\_USR | 2.2223 | 0.0263 | 0.0421 |
| S\_DER | F\_MRH | 2.1767 | 0.0295 | 0.0467 |
| F\_MKH | S\_FRH | 2.1339 | 0.0329 | 0.0514 |
| F\_BUT | L\_USR | 2.1018 | 0.0356 | 0.0550 |
| S\_MIL | F\_BUT | 2.0595 | 0.0394 | 0.0602 |
| F\_TOU | F\_BUT | 1.9219 | 0.0546 | 0.0825 |
| F\_TOU | L\_USR | 1.8511 | 0.0642 | 0.0959 |
| F\_MIL | F\_MRH | 1.7714 | 0.0765 | 0.1131 |
| F\_MIL | S\_MIL | 1.7463 | 0.0808 | 0.1169 |
| S\_MIL | F\_STN | 1.7458 | 0.0808 | 0.1169 |
| F\_MRH | S\_MIL | 1.7340 | 0.0829 | 0.1187 |
| F\_NIM | F\_STN | 1.6828 | 0.0924 | 0.1309 |
| F\_FRH | S\_DER | 1.6697 | 0.0950 | 0.1312 |
| F\_STN | L\_USR | 1.6679 | 0.0953 | 0.1312 |
| F\_MRH | F\_STN | 1.6671 | 0.0955 | 0.1312 |
| F\_MIL | F\_COL | 1.6313 | 0.1028 | 0.1398 |
| F\_COL | F\_TOU | 1.5766 | 0.1149 | 0.1547 |
| F\_MIL | S\_DER | 1.4998 | 0.1337 | 0.1783 |
| F\_NIM | F\_MER | 1.4075 | 0.1593 | 0.2103 |
| F\_MRH | S\_FRH | 1.3826 | 0.1668 | 0.2181 |
| F\_MER | F\_STN | 1.3571 | 0.1748 | 0.2264 |
| F\_COL | F\_MKH | 1.3062 | 0.1915 | 0.2445 |
| F\_MRH | F\_MER | 1.3034 | 0.1924 | 0.2445 |
| F\_MKH | F\_DER | 1.2651 | 0.2058 | 0.2592 |
| F\_COL | F\_DER | 1.2378 | 0.2158 | 0.2693 |
| F\_TOU | F\_MER | 1.2137 | 0.2249 | 0.2781 |
| F\_MRH | F\_TOU | 1.1348 | 0.2564 | 0.3141 |
| F\_FRH | F\_MIL | 1.0988 | 0.2718 | 0.3300 |
| F\_MKH | S\_MIL | 1.0733 | 0.2831 | 0.3407 |
| F\_COL | F\_NIM | 1.0212 | 0.3072 | 0.3665 |
| F\_BUT | F\_STN | 1.0124 | 0.3114 | 0.3683 |
| F\_NIM | S\_MIL | 0.9862 | 0.3240 | 0.3799 |
| F\_MRH | F\_MKH | 0.9442 | 0.3451 | 0.4011 |
| F\_DER | F\_STN | 0.8775 | 0.3802 | 0.4382 |
| F\_TOU | F\_NIM | 0.8558 | 0.3921 | 0.4481 |
| F\_COL | F\_MRH | 0.8391 | 0.4014 | 0.4549 |
| F\_BUT | S\_FRH | 0.7339 | 0.4630 | 0.5181 |
| F\_NIM | F\_DER | 0.7310 | 0.4648 | 0.5181 |
| F\_STN | S\_FRH | 0.6749 | 0.4998 | 0.5526 |
| S\_MIL | S\_FRH | 0.5755 | 0.5650 | 0.6179 |
| F\_MRH | F\_NIM | 0.5648 | 0.5722 | 0.6179 |
| S\_DER | F\_COL | 0.5644 | 0.5725 | 0.6179 |
| F\_MKH | F\_TOU | 0.4981 | 0.6184 | 0.6622 |
| F\_TOU | S\_MIL | 0.4701 | 0.6383 | 0.6782 |
| S\_FRH | L\_USR | 0.4292 | 0.6678 | 0.7040 |
| F\_TOU | F\_DER | 0.3595 | 0.7192 | 0.7510 |
| F\_MER | S\_MIL | 0.3539 | 0.7234 | 0.7510 |
| F\_DER | S\_MIL | 0.3217 | 0.7477 | 0.7704 |
| F\_MKH | F\_STN | 0.3003 | 0.7639 | 0.7811 |
| F\_STN | F\_TOU | 0.1999 | 0.8416 | 0.8542 |
| F\_DER | S\_FRH | 0.1105 | 0.9120 | 0.9188 |
| F\_MER | F\_DER | 0.0626 | 0.9501 | 0.9501 |

Table S8: Comparison of mean +/- standard deviation, minimum, and maximum values of the nucleotide diversity (pi) across loci among individuals grouped by run within tributary.

| pop | mean | std | max | min |
| --- | --- | --- | --- | --- |
| F\_FRH | 0.001058 | 0.001272 | 0.013546 | 0 |
| F\_MIL | 0.001056 | 0.001298 | 0.015935 | 0 |
| S\_FRH | 0.001048 | 0.001392 | 0.014501 | 0 |
| S\_DER | 0.001046 | 0.001275 | 0.015410 | 0 |
| F\_COL | 0.001045 | 0.001275 | 0.014935 | 0 |
| S\_MIL | 0.001043 | 0.001315 | 0.016022 | 0 |
| F\_MRH | 0.001042 | 0.001311 | 0.015478 | 0 |
| F\_DER | 0.001038 | 0.001294 | 0.014056 | 0 |
| F\_MKH | 0.001035 | 0.001276 | 0.015955 | 0 |
| F\_MER | 0.001035 | 0.001274 | 0.014661 | 0 |
| F\_NIM | 0.001034 | 0.001271 | 0.014977 | 0 |
| F\_TOU | 0.001033 | 0.001275 | 0.014795 | 0 |
| L\_USR | 0.001029 | 0.001289 | 0.014423 | 0 |
| F\_STN | 0.001028 | 0.001281 | 0.014589 | 0 |
| F\_BUT | 0.001026 | 0.001292 | 0.014613 | 0 |
| S\_BUT | 0.001012 | 0.001308 | 0.015957 | 0 |
| W\_USR | 0.000790 | 0.001212 | 0.013255 | 0 |

#### 4.7 Fixed loci

**Fixed loci** are loci that are not polymorphic for a given set of individuals, i.e. all individuals are homozygous for the same allele. The **global allele frequency/counts** is the allele frequency or count for a given locus across all individuals.

Differences in sample size may create bias for this comparison, as sample sizes that are (too) low can result in not all alleles being present in a population being sampled in a sufficiently representative manner, which can lead loci to appear fixed even though they do have rare alleles that were simply not sampled. Sample sizes are quite variable across run types, but for individuals grouped by run type withing tributaries, the sample sizes do become much more comparable.

**Comparison of fixed loci within tributaries within runs**

Identify number of fixed loci within individuals grouped by tributary and run type.

Number of fixed alleles for individuals of each tributary grouped by run type (using rarefied allele counts).

| GRP | n |
| --- | --- |
| F\_COL | 3835 |
| F\_MIL | 3574 |
| F\_DER | 4898 |
| F\_BUT | 4702 |
| F\_FRH | 4385 |
| F\_NIM | 4369 |
| F\_MKH | 3788 |
| F\_STN | 3609 |
| F\_TOU | 3655 |
| F\_MRH | 3504 |
| F\_MER | 4191 |
| L\_USR | 6416 |
| W\_USR | 3853 |
| S\_MIL | 4910 |
| S\_DER | 5179 |
| S\_BUT | 6210 |
| S\_FRH | 4423 |

Identify loci that are fixed in more than on set of individuals grouped by tributary and run type: The set size (horizontal green bars) indicates the total number of loci fixed in a given location, the intersection size (vertical orange bars) indicates the number of loci fixed only in a single location (single blue dot) or in two, three, or four locations (indicated by blue dots connected by line).

Calculate allelic richness across all individuals in a data set.

```
# calculate global allelic richness
setPop(gen) <- ~OVERALL

# calculate allele counts using rarefaction
dat <- genind2hierfstat(gen)

df <- allelic.richness(dat,
                       diploid = TRUE)

ar <- as.data.frame(df$Ar) %>%
  rownames_to_column("LOCUS") %>%
  select(-3)

colnames(ar) <- c("LOCUS", "OVERALL")

write_delim(ar, "results/rarefied.allelecount", delim = "\t")
```

Determine the global rarefied allele counts for loci that are fixed in at least one group.

Distribution of global rarefied allele counts for loci fixed across individuals grouped by tributary within runs.

Table S9: Comparison of mean, median, 25th and 75th quantile of global diversity at fixed loci for individuals grouped by run/tributary.

| GRP | mean | median | Q1 | Q3 |
| --- | --- | --- | --- | --- |
| F\_COL | 1.20 | 1.13 | 1.07 | 1.30 |
| F\_MIL | 1.19 | 1.13 | 1.07 | 1.29 |
| F\_DER | 1.30 | 1.19 | 1.07 | 1.46 |
| F\_BUT | 1.28 | 1.19 | 1.07 | 1.44 |
| F\_FRH | 1.26 | 1.14 | 1.07 | 1.40 |
| F\_NIM | 1.25 | 1.14 | 1.07 | 1.39 |
| F\_MKH | 1.20 | 1.13 | 1.07 | 1.29 |
| F\_STN | 1.18 | 1.13 | 1.07 | 1.25 |
| F\_TOU | 1.19 | 1.13 | 1.07 | 1.29 |
| F\_MRH | 1.19 | 1.13 | 1.07 | 1.29 |
| F\_MER | 1.24 | 1.13 | 1.07 | 1.36 |
| L\_USR | 1.46 | 1.35 | 1.12 | 1.79 |
| W\_USR | 1.31 | 1.19 | 1.07 | 1.48 |
| S\_MIL | 1.36 | 1.24 | 1.07 | 1.59 |
| S\_DER | 1.38 | 1.25 | 1.07 | 1.63 |
| S\_BUT | 1.41 | 1.30 | 1.07 | 1.70 |
| S\_FRH | 1.28 | 1.19 | 1.07 | 1.43 |

Compare the distribution of fixed loci across the genome.

Proportion of loci on chromosome of fixed loci per chromosome for individuals grouped by run type.

| CHR | total | F\_COL | F\_MIL | F\_DER | F\_BUT | F\_FRH | F\_NIM | F\_MKH | F\_STN | F\_TOU | F\_MRH | F\_MER | L\_USR | W\_USR | S\_MIL | S\_DER | S\_BUT | S\_FRH |
| --- | --- | --- | --- | --- | --- | --- | --- | --- | --- | --- | --- | --- | --- | --- | --- | --- | --- | --- |
| 1 | 8167 | 2.8 | 2.4 | 3.2 | 3.3 | 2.9 | 3.1 | 2.9 | 2.5 | 2.5 | 2.6 | 3.1 | 4.7 | 2.9 | 3.6 | 3.7 | 4.2 | 3.0 |
| 2 | 4869 | 2.4 | 2.1 | 3.1 | 2.9 | 2.9 | 2.6 | 2.1 | 2.3 | 2.2 | 2.4 | 2.5 | 4.1 | 2.5 | 3.1 | 3.2 | 4.0 | 3.1 |
| 3 | 5253 | 2.7 | 2.5 | 3.6 | 3.4 | 3.2 | 3.1 | 2.7 | 2.6 | 2.6 | 2.3 | 3.0 | 4.6 | 2.7 | 3.7 | 3.6 | 4.5 | 3.0 |
| 4 | 5657 | 2.4 | 2.2 | 3.3 | 2.9 | 2.7 | 2.7 | 2.4 | 2.3 | 2.2 | 2.2 | 2.5 | 3.5 | 2.4 | 3.1 | 3.0 | 3.7 | 2.6 |
| 5 | 6268 | 2.8 | 2.6 | 3.6 | 3.5 | 3.1 | 3.3 | 2.8 | 2.6 | 2.6 | 2.4 | 3.0 | 4.4 | 2.6 | 3.4 | 3.7 | 4.1 | 3.1 |
| 6 | 6408 | 2.5 | 2.4 | 3.4 | 3.1 | 2.9 | 2.9 | 2.7 | 2.2 | 2.4 | 2.4 | 2.8 | 4.3 | 2.5 | 3.2 | 3.3 | 3.8 | 3.0 |
| 7 | 5707 | 2.5 | 2.2 | 3.2 | 3.0 | 2.8 | 2.7 | 2.3 | 2.2 | 2.5 | 2.1 | 2.6 | 4.2 | 2.4 | 2.9 | 3.4 | 3.7 | 2.6 |
| 8 | 6666 | 2.3 | 2.1 | 2.9 | 3.0 | 2.7 | 3.0 | 2.5 | 2.5 | 2.3 | 2.3 | 2.7 | 4.3 | 2.4 | 3.0 | 3.4 | 4.3 | 3.1 |
| 9 | 6000 | 2.8 | 2.7 | 3.3 | 3.3 | 3.2 | 3.2 | 2.6 | 2.7 | 2.6 | 2.2 | 2.8 | 4.2 | 2.6 | 3.3 | 3.4 | 4.0 | 3.1 |
| 10 | 5351 | 2.5 | 2.2 | 3.1 | 2.9 | 2.7 | 2.6 | 2.1 | 2.1 | 2.2 | 2.1 | 2.8 | 3.9 | 2.4 | 3.3 | 3.5 | 3.8 | 2.7 |
| 11 | 3095 | 2.8 | 2.3 | 3.1 | 3.3 | 3.2 | 2.8 | 2.7 | 2.4 | 2.5 | 2.5 | 2.6 | 4.9 | 2.6 | 3.2 | 3.4 | 4.4 | 3.1 |
| 12 | 5620 | 2.0 | 1.8 | 2.5 | 2.4 | 2.2 | 2.2 | 1.8 | 1.7 | 1.9 | 1.9 | 2.1 | 3.3 | 2.1 | 2.6 | 2.6 | 3.2 | 2.3 |
| 13 | 5979 | 2.3 | 2.2 | 3.1 | 3.4 | 2.7 | 2.9 | 2.6 | 2.3 | 2.4 | 2.2 | 2.8 | 4.1 | 2.6 | 3.1 | 3.2 | 4.2 | 2.9 |
| 14 | 4284 | 2.5 | 2.4 | 3.5 | 3.2 | 2.7 | 2.6 | 2.3 | 2.5 | 2.7 | 2.3 | 2.9 | 4.7 | 2.5 | 3.5 | 3.5 | 4.4 | 2.8 |
| 15 | 3245 | 1.9 | 2.3 | 2.9 | 3.0 | 2.9 | 2.7 | 1.8 | 2.0 | 2.0 | 2.2 | 2.3 | 3.9 | 2.4 | 2.9 | 3.5 | 3.9 | 3.0 |
| 16 | 5093 | 2.4 | 2.5 | 3.3 | 3.0 | 2.9 | 2.7 | 2.5 | 2.4 | 2.4 | 2.4 | 2.7 | 4.3 | 2.8 | 3.4 | 3.4 | 3.9 | 2.7 |
| 17 | 2070 | 2.2 | 2.2 | 2.7 | 3.0 | 2.6 | 2.3 | 2.1 | 2.1 | 2.1 | 2.3 | 2.7 | 4.2 | 2.4 | 2.8 | 3.0 | 3.9 | 2.6 |
| 18 | 2958 | 2.5 | 2.2 | 3.0 | 2.8 | 2.7 | 2.9 | 2.3 | 2.6 | 2.3 | 2.3 | 2.8 | 4.1 | 2.5 | 3.2 | 3.5 | 4.2 | 2.6 |
| 19 | 5071 | 2.5 | 2.5 | 3.3 | 3.2 | 2.9 | 2.9 | 2.6 | 2.5 | 2.4 | 2.4 | 2.8 | 4.1 | 2.7 | 3.7 | 3.8 | 4.8 | 2.9 |
| 20 | 3797 | 2.5 | 2.4 | 3.5 | 2.9 | 2.9 | 2.8 | 2.4 | 2.3 | 2.6 | 2.5 | 2.9 | 4.6 | 2.3 | 3.3 | 3.6 | 4.2 | 3.1 |
| 21 | 2937 | 2.8 | 2.9 | 3.4 | 2.9 | 3.5 | 2.8 | 2.7 | 2.8 | 2.7 | 2.6 | 3.0 | 4.3 | 2.9 | 3.5 | 4.0 | 4.1 | 2.9 |
| 22 | 3447 | 2.5 | 2.3 | 3.4 | 3.0 | 2.9 | 3.0 | 2.5 | 2.2 | 2.5 | 2.3 | 2.5 | 4.9 | 2.5 | 3.3 | 3.5 | 4.1 | 3.0 |
| 23 | 1756 | 2.7 | 2.1 | 3.2 | 2.9 | 2.7 | 2.5 | 2.5 | 2.1 | 2.2 | 2.0 | 2.5 | 3.8 | 2.3 | 3.2 | 3.2 | 3.5 | 2.9 |
| 24 | 2314 | 2.1 | 2.0 | 3.2 | 3.0 | 2.7 | 2.5 | 2.5 | 2.2 | 2.0 | 2.0 | 2.5 | 3.9 | 2.4 | 3.1 | 3.2 | 4.3 | 3.0 |
| 25 | 2980 | 2.4 | 2.4 | 3.1 | 3.0 | 2.6 | 3.1 | 2.2 | 2.0 | 2.3 | 2.2 | 2.9 | 4.3 | 2.3 | 3.2 | 3.8 | 4.3 | 2.6 |
| 26 | 4047 | 2.7 | 2.5 | 3.5 | 3.1 | 3.1 | 3.1 | 2.9 | 2.6 | 2.3 | 2.4 | 2.7 | 4.8 | 2.5 | 3.2 | 3.6 | 4.3 | 3.0 |
| 27 | 1997 | 2.8 | 2.3 | 3.0 | 2.8 | 3.0 | 3.1 | 2.6 | 2.5 | 2.6 | 2.4 | 3.2 | 3.3 | 2.5 | 3.2 | 2.9 | 3.5 | 3.1 |
| 28 | 3291 | 2.2 | 2.1 | 3.1 | 2.9 | 2.4 | 2.8 | 2.3 | 2.5 | 2.4 | 2.1 | 2.4 | 4.0 | 2.6 | 2.9 | 3.1 | 3.4 | 2.8 |
| 29 | 2272 | 3.0 | 2.6 | 3.5 | 3.7 | 3.3 | 3.2 | 2.8 | 2.6 | 3.0 | 2.3 | 2.9 | 3.9 | 2.8 | 3.4 | 3.5 | 4.6 | 3.2 |
| 30 | 3383 | 3.0 | 3.0 | 3.4 | 3.2 | 3.5 | 3.3 | 3.0 | 2.7 | 2.7 | 2.5 | 3.6 | 4.8 | 2.9 | 3.4 | 3.8 | 4.9 | 3.2 |
| 31 | 2772 | 2.3 | 2.2 | 2.8 | 2.8 | 2.5 | 2.8 | 2.4 | 2.2 | 1.9 | 1.8 | 2.7 | 3.5 | 2.3 | 2.6 | 2.8 | 3.7 | 2.7 |
| 32 | 1274 | 1.3 | 1.6 | 2.0 | 2.0 | 2.1 | 1.6 | 1.6 | 1.7 | 1.6 | 1.6 | 1.9 | 3.1 | 1.8 | 2.4 | 2.2 | 3.5 | 2.4 |
| 33 | 3255 | 2.1 | 2.1 | 2.4 | 2.8 | 2.4 | 2.3 | 2.2 | 2.0 | 2.0 | 1.9 | 2.2 | 3.8 | 1.8 | 2.7 | 2.9 | 3.6 | 2.4 |
| 34 | 1094 | 2.0 | 1.6 | 2.2 | 2.2 | 2.4 | 2.9 | 1.9 | 1.8 | 2.3 | 2.2 | 2.4 | 2.8 | 2.3 | 2.7 | 2.8 | 3.0 | 2.8 |

Determine the null distribution of the proportion of loci expected to be fixed per chromosome if fixed loci are randomly distributed among chromosomes. Maintain the total number of loci fixed for a run/tributary group and the total number of loci per chromosome in the data by shuffeling chromosome designations across loci. Then calculate the proportion of loci fixed on a chromosome for each run/tributary group and determining if the observed value falls is higher than the 5th and lower than the 95th percentile; determine significance as observed value lying outside observed distribution.

```
obs <- tidy_ar %>%
   mutate(FIXED = ifelse(AR == 1, TRUE, FALSE)) %>%
   mutate(GRP = ordered(GRP, levels = trib_runs)) %>%
   left_join(contigs) %>%
    filter(!is.na(CHROMOSOME)) %>%
    select(GRP, LOCUS, FIXED, CHROMOSOME)

reps <- 1000

sim <- list()

for(i in 1:reps){

    sim[[i]] <- obs %>%
        group_by(GRP) %>%
        mutate(SIM = sample(CHROMOSOME, replace = FALSE)) %>%
        ungroup() %>%
        filter(FIXED == TRUE) %>%
        count(GRP, SIM) %>%
        left_join(n_chrom, by = c("SIM" = "CHROMOSOME")) %>%
        mutate(percent = (n/total)*100)
}

null_dist <- ldply(sim, data.frame) %>%
    group_by(GRP, SIM) %>%
    summarize(min = min(percent),
              max = max(percent)) %>%
    ungroup() %>%
    rename(CHROMOSOME = SIM) %>%
    left_join(pattern) %>%
    select(GRP, CHROMOSOME, total, n, percent, min, max) %>%
    mutate(sign = ifelse(percent > min & percent < max, "expected", "significant"))
```

Compare observed and simulated distributions.

Table S10: Chromosomes with significantly higher/lower than expected (outside simulated null distribution) number of fixed loci for each run/tributary group.

| GRP | CHROMOSOME | total | n | percent | min | max |
| --- | --- | --- | --- | --- | --- | --- |
| F\_COL | 9 | 6000 | 169 | 2.82 | 1.80 | 2.82 |
| F\_MIL | 9 | 6000 | 161 | 2.68 | 1.62 | 2.65 |
| F\_DER | 19 | 5071 | 166 | 3.27 | 3.29 | 4.56 |
| F\_DER | 33 | 3255 | 77 | 2.37 | 2.46 | 4.36 |
| F\_FRH | 9 | 6000 | 191 | 3.18 | 2.12 | 3.15 |
| F\_NIM | 9 | 6000 | 190 | 3.17 | 1.98 | 3.17 |
| F\_NIM | 33 | 3255 | 74 | 2.27 | 2.27 | 3.66 |
| F\_STN | 9 | 6000 | 161 | 2.68 | 1.65 | 2.63 |
| L\_USR | 19 | 5071 | 208 | 4.10 | 4.30 | 5.74 |
| W\_USR | 33 | 3255 | 59 | 1.81 | 1.87 | 3.41 |

Table S11: Mean +/- sd, minimum, and maximum percent of loci per chromosome that are fixed across a group of individuals with the same run type for a given tributary.

| GRP | mean | sd | min | max |
| --- | --- | --- | --- | --- |
| F\_COL | 2.45 | 0.35 | 1.26 | 3.04 |
| F\_MIL | 2.30 | 0.30 | 1.57 | 3.02 |
| F\_DER | 3.12 | 0.38 | 2.04 | 3.59 |
| F\_BUT | 2.99 | 0.34 | 1.96 | 3.65 |
| F\_FRH | 2.82 | 0.32 | 2.12 | 3.47 |
| F\_NIM | 2.79 | 0.35 | 1.57 | 3.31 |
| F\_MKH | 2.42 | 0.33 | 1.65 | 2.99 |
| F\_STN | 2.31 | 0.28 | 1.73 | 2.83 |
| F\_TOU | 2.35 | 0.29 | 1.57 | 2.99 |
| F\_MRH | 2.24 | 0.22 | 1.57 | 2.58 |
| F\_MER | 2.70 | 0.32 | 1.88 | 3.58 |
| L\_USR | 4.10 | 0.51 | 2.83 | 4.91 |
| W\_USR | 2.46 | 0.25 | 1.81 | 2.90 |
| S\_MIL | 3.15 | 0.32 | 2.43 | 3.69 |
| S\_DER | 3.33 | 0.38 | 2.20 | 3.98 |
| S\_BUT | 4.00 | 0.43 | 3.02 | 4.94 |
| S\_FRH | 2.86 | 0.24 | 2.35 | 3.21 |

#### 4.8 Singletons

**Singletons** are alleles that only occur in a single individual. This could be a locus that is only variable in one individual or a locus that has multiple alleles though one (or more) of those alleles are extremely rare occurring in only one individual.

```
# group all individuals in single group
setPop(gen) <- ~OVERALL

# calculate basic stats
dat <- genind2hierfstat(gen)
stats <- basic.stats(dat)

# extract & format allele frequencies per locus into single df
f <- stats$pop.freq

freq <- list()

for(l in names(f)){

freq[[l]] <- as.data.frame(f[[l]]) %>%
  filter(Var2 == 1) %>%
  rename(ALLELE = x,
         FRQ = Freq) %>%
  select(ALLELE, FRQ)

}

freq <- ldply(freq, data.frame) %>%
  rename(LOCUS = `.id`)

# identify singletons
Nind <- length(indNames(gen))

min <- round(1/(2*Nind), digits = 4)

singletons <- freq %>%
  filter(FRQ <= min) %>%
  mutate(ALLELE = as.integer(ALLELE))
```

Total number of alleles across all loci in the data set is 29873, the total number of singletons is 363 (1.2% of loci) exhibit at least one singleton.

Number of loci with x singletons.

| singletons | loci |
| --- | --- |
| 1 | 347 |
| 2 | 8 |

Compare distribution of loci with singletons across the genome to random null distribution.

Chromosomes with significantly higher/lower than expected (outside simulated null distribution) number of loci with observed singletons.

| CHROMOSOME | min | max | n |
| --- | --- | --- | --- |

Determine the number of singletons individuals in each group carry.

Comparison of number of singletons per individual (individual circles); for individuals grouped by tributary and run type.

Table S12: Mean, median, 25th and 27th quantile of the number of singletons observer per individual for individuals grouped by run/tributary.

| RUN\_LOC | mean | median | Q1 | Q3 |
| --- | --- | --- | --- | --- |
| F\_COL | 1.55 | 1 | 1.00 | 2.00 |
| F\_MIL | 1.38 | 1 | 1.00 | 2.00 |
| F\_DER | 1.67 | 1 | 1.00 | 2.00 |
| F\_BUT | 1.33 | 1 | 1.00 | 1.00 |
| F\_FRH | 1.71 | 1 | 1.00 | 2.00 |
| F\_NIM | 2.21 | 1 | 1.00 | 3.75 |
| F\_MKH | 2.00 | 2 | 1.25 | 2.75 |
| F\_STN | 1.00 | 1 | 1.00 | 1.00 |
| F\_TOU | 1.36 | 1 | 1.00 | 1.75 |
| F\_MRH | 1.71 | 2 | 1.00 | 2.00 |
| F\_MER | 1.47 | 1 | 1.00 | 2.00 |
| L\_USR | 2.23 | 2 | 1.00 | 2.00 |
| W\_USR | 1.78 | 1 | 1.00 | 2.00 |
| S\_MIL | 1.82 | 1 | 1.00 | 2.00 |
| S\_DER | 3.00 | 2 | 1.00 | 2.75 |
| S\_BUT | 4.00 | 3 | 2.50 | 4.50 |
| S\_FRH | 1.25 | 1 | 1.00 | 1.25 |

#### 4.9 Loci variable in only one group of individuals (“private polymorphisms”)

Identify loci that are polymorphic in one set of individuals (i.e. \(N(alleles) > 1\)) but that locus is *not* variable in *any other* set of individuals (i.e. fixed across all other sets of loci). The polymorphisms must occur in at least one individual of a group (i.e. singletons) but could be variable in multiple individuals, the allele other groups are fixed for may also occur in the group it is variable at, i.e. these are *not* necessarily private alleles, rather they are loci exclusively polymorphic in a single group.

**Identify loci only variable among individuals from the same tributary by run type (private polymorphisms)**

```
tidy_ar <- read_delim("results/gendiv.locstats", delim = "\t") %>%
  select(GRP, LOCUS, N_ALLELES) %>%
  filter(GRP %in% trib_runs)

pops <- unique(tidy_ar$GRP)

# df for results
results <- setNames(data.frame(matrix(ncol = 2, nrow = 0)), 
                    c("LOCUS", "POLYMORPHIC")) %>%
  mutate(LOCUS = as.character(LOCUS),
         POLYMORPHIC = as.character(POLYMORPHIC))

# loop over to identify loci polymorphic in one pop but fixed in all others
for(p in pops){

# find loci polymorphic in pop
polymorphic <- tidy_ar %>%
  filter(N_ALLELES > 1 & GRP == p)

# find loci fixed in all other pops
fixed <- tidy_ar %>%
  filter(!GRP == p) %>%
  group_by(LOCUS) %>%
  count(N_ALLELES == 1) %>%
  filter(`N_ALLELES == 1` == TRUE) %>%
  filter(n == length(pops)-1) %>%
  mutate(POLYMORPHIC = p) %>%
  select(LOCUS, POLYMORPHIC)

results <- bind_rows(results, fixed)

}

pp <- results %>%
    count(LOCUS) %>%
    filter(n == 1)

results <- results %>%
    filter(LOCUS %in% pp$LOCUS)
```

Determine differences in number of loci with private polymorphisms among run/tributary groups.

Number of private polymorphisms identified in individuals of the same run type within a tributary.

Number of loci polymorphic in individuals of the same run type within a tributary that are not variable in individuals of any others group (based on observed allele counts).

| POLYMORPHIC | n |
| --- | --- |
| S\_DER | 153 |
| S\_BUT | 97 |
| F\_COL | 89 |
| L\_USR | 84 |
| F\_FRH | 83 |
| F\_NIM | 82 |
| S\_MIL | 70 |
| W\_USR | 70 |
| F\_MKH | 57 |
| F\_MER | 55 |
| F\_TOU | 54 |
| F\_MRH | 49 |
| F\_MIL | 47 |
| F\_STN | 46 |
| F\_DER | 44 |
| F\_BUT | 33 |
| S\_FRH | 18 |

Compare the distribution of exclusively polymorphic loci across the genome.

Determine the percent of loci on a chromosome that are only polymorphic for a set of individuals from the same tributary of the same run type.

Percent loci on a chromosome that are only polymorphic for a set of individuals from the same tributary of the same run type.

| CHROMOSOME | total | F\_COL | F\_MIL | F\_DER | F\_BUT | F\_FRH | F\_NIM | F\_MKH | F\_STN | F\_TOU | F\_MRH | F\_MER | L\_USR | W\_USR | S\_MIL | S\_DER | S\_BUT | S\_FRH |
| --- | --- | --- | --- | --- | --- | --- | --- | --- | --- | --- | --- | --- | --- | --- | --- | --- | --- | --- |
| 1 | 8167 | 0.10 | 0.04 | 0.04 | 0.04 | 0.10 | 0.07 | 0.09 | 0.04 | 0.02 | 0.04 | 0.04 | 0.02 | 0.05 | 0.07 | 0.07 | 0.11 | 0.04 |
| 2 | 4869 | 0.06 | 0.04 | 0.02 | 0.02 | 0.04 | 0.08 | 0.02 | NA | 0.04 | NA | 0.02 | 0.02 | NA | NA | 0.12 | 0.02 | NA |
| 3 | 5253 | 0.15 | 0.02 | 0.04 | NA | 0.04 | 0.02 | 0.04 | 0.06 | 0.02 | 0.02 | 0.04 | 0.06 | 0.04 | 0.06 | 0.13 | 0.06 | NA |
| 4 | 5657 | 0.05 | 0.02 | 0.04 | 0.02 | 0.02 | 0.04 | 0.02 | 0.04 | 0.07 | 0.05 | 0.11 | 0.05 | 0.04 | 0.02 | 0.12 | 0.05 | NA |
| 5 | 6268 | 0.11 | NA | 0.03 | 0.03 | 0.10 | 0.08 | 0.03 | 0.10 | NA | 0.02 | 0.02 | 0.10 | 0.05 | NA | 0.16 | 0.11 | 0.02 |
| 6 | 6408 | 0.02 | 0.03 | 0.02 | 0.05 | 0.06 | 0.06 | 0.05 | 0.02 | 0.02 | 0.02 | 0.05 | 0.05 | 0.05 | 0.08 | 0.09 | 0.08 | 0.02 |
| 7 | 5707 | 0.12 | 0.07 | NA | NA | 0.07 | 0.04 | 0.05 | 0.07 | 0.02 | 0.02 | 0.02 | 0.07 | 0.04 | 0.07 | 0.18 | 0.07 | 0.04 |
| 8 | 6666 | 0.05 | 0.05 | 0.03 | 0.05 | 0.02 | 0.05 | 0.03 | NA | 0.03 | 0.02 | 0.02 | NA | 0.08 | 0.05 | 0.02 | 0.05 | 0.03 |
| 9 | 6000 | 0.12 | 0.03 | 0.03 | 0.03 | 0.10 | 0.05 | NA | 0.07 | 0.05 | 0.03 | 0.03 | 0.05 | 0.02 | 0.07 | 0.13 | 0.05 | NA |
| 10 | 5351 | 0.02 | NA | 0.04 | 0.04 | 0.06 | 0.02 | 0.02 | 0.02 | 0.06 | 0.04 | NA | 0.09 | 0.09 | NA | 0.11 | 0.07 | NA |
| 11 | 3095 | 0.03 | 0.03 | NA | 0.03 | 0.06 | 0.06 | 0.03 | NA | 0.10 | 0.10 | NA | 0.03 | NA | 0.03 | NA | 0.06 | 0.03 |
| 12 | 5620 | 0.05 | 0.02 | 0.04 | NA | 0.07 | 0.05 | 0.02 | 0.02 | 0.04 | 0.02 | 0.05 | 0.04 | 0.02 | 0.05 | 0.07 | 0.11 | NA |
| 13 | 5979 | 0.08 | 0.02 | NA | NA | 0.05 | 0.02 | 0.05 | 0.03 | 0.03 | NA | 0.03 | 0.07 | 0.03 | 0.05 | 0.07 | 0.03 | 0.02 |
| 14 | 4284 | 0.09 | 0.05 | NA | 0.02 | 0.09 | 0.05 | 0.12 | 0.02 | 0.02 | 0.02 | 0.02 | 0.05 | 0.05 | 0.05 | 0.07 | 0.02 | NA |
| 15 | 3245 | NA | NA | 0.03 | 0.03 | NA | 0.06 | 0.03 | 0.06 | 0.06 | 0.03 | 0.06 | 0.03 | NA | NA | 0.12 | 0.03 | 0.03 |
| 16 | 5093 | 0.02 | 0.06 | 0.06 | NA | 0.02 | 0.14 | 0.04 | 0.02 | 0.08 | 0.04 | NA | 0.06 | 0.02 | NA | 0.08 | 0.04 | 0.02 |
| 17 | 2070 | 0.10 | 0.05 | NA | NA | NA | 0.10 | NA | NA | 0.10 | NA | 0.05 | NA | NA | NA | 0.14 | 0.10 | 0.05 |
| 18 | 2958 | 0.03 | NA | 0.10 | NA | 0.03 | NA | NA | 0.07 | NA | 0.14 | NA | 0.03 | 0.03 | 0.07 | 0.14 | 0.10 | NA |
| 19 | 5071 | 0.06 | 0.12 | 0.04 | 0.04 | NA | 0.12 | 0.08 | NA | NA | 0.06 | 0.04 | 0.04 | 0.04 | 0.02 | 0.04 | 0.04 | NA |
| 20 | 3797 | 0.03 | NA | NA | NA | 0.03 | NA | NA | NA | 0.05 | 0.05 | 0.03 | 0.08 | 0.03 | 0.05 | 0.16 | 0.08 | NA |
| 21 | 2937 | 0.03 | 0.07 | 0.03 | NA | 0.03 | 0.07 | 0.03 | 0.03 | 0.03 | 0.03 | 0.03 | 0.10 | 0.17 | 0.03 | 0.03 | NA | NA |
| 22 | 3447 | 0.09 | 0.03 | 0.03 | NA | 0.03 | NA | NA | 0.03 | 0.03 | NA | 0.03 | 0.09 | 0.12 | 0.09 | 0.03 | 0.03 | NA |
| 23 | 1756 | 0.06 | NA | NA | NA | 0.06 | NA | 0.06 | NA | NA | 0.06 | NA | 0.11 | 0.17 | NA | 0.17 | NA | NA |
| 24 | 2314 | 0.04 | 0.04 | 0.04 | NA | 0.04 | 0.13 | NA | 0.04 | 0.04 | NA | 0.09 | 0.09 | NA | NA | 0.04 | 0.09 | NA |
| 25 | 2980 | NA | 0.03 | NA | NA | NA | NA | 0.10 | NA | 0.03 | NA | 0.10 | 0.07 | 0.03 | 0.07 | 0.13 | 0.07 | NA |
| 26 | 4047 | 0.05 | 0.05 | 0.02 | NA | 0.05 | 0.10 | 0.02 | 0.02 | 0.05 | 0.07 | NA | 0.02 | 0.05 | 0.05 | 0.27 | 0.07 | NA |
| 27 | 1997 | NA | 0.05 | NA | NA | 0.15 | 0.05 | 0.10 | NA | 0.05 | 0.10 | 0.05 | 0.05 | 0.05 | 0.05 | 0.10 | 0.05 | 0.05 |
| 28 | 3291 | 0.15 | 0.06 | 0.03 | NA | 0.09 | NA | NA | 0.06 | 0.03 | 0.06 | NA | 0.03 | NA | 0.06 | 0.03 | 0.12 | NA |
| 29 | 2272 | NA | NA | 0.04 | NA | 0.04 | NA | 0.04 | 0.09 | NA | 0.04 | 0.13 | 0.04 | NA | 0.04 | 0.09 | 0.09 | NA |
| 30 | 3383 | 0.03 | NA | 0.15 | NA | 0.06 | 0.03 | 0.03 | NA | 0.03 | NA | 0.09 | 0.06 | 0.06 | 0.03 | 0.06 | 0.09 | 0.03 |
| 31 | 2772 | NA | 0.04 | 0.04 | 0.14 | 0.14 | NA | 0.07 | 0.04 | NA | 0.04 | 0.07 | NA | 0.04 | 0.07 | 0.04 | NA | NA |
| 32 | 1274 | 0.08 | NA | NA | 0.08 | NA | NA | 0.08 | NA | NA | NA | NA | NA | 0.08 | NA | 0.08 | 0.08 | NA |
| 33 | 3255 | NA | NA | NA | 0.03 | NA | 0.03 | NA | 0.06 | 0.03 | NA | NA | 0.06 | 0.06 | 0.06 | 0.12 | 0.12 | NA |
| 34 | 1094 | NA | 0.09 | 0.09 | NA | 0.18 | 0.09 | 0.09 | NA | NA | NA | NA | NA | 0.18 | NA | 0.09 | NA | NA |

Generate null distribution to determine if some groups exhibit more/less than the expected number of loci with exclusive polymorphisms for sets of individuals.

Chromosomes with significantly higher/lower than expected (outside simulated null distribution) number of private polymorhisms for each run/tributary group

| GRP | CHROMOSOME | total | n | percent | min | max | sign |
| --- | --- | --- | --- | --- | --- | --- | --- |
| F\_COL | 6 | 6408 | 1 | 0.0156 | 0.0156 | 0.1873 | lower |
| F\_COL | 10 | 5351 | 1 | 0.0187 | 0.0187 | 0.2056 | lower |
| F\_COL | 11 | 3095 | 1 | 0.0323 | 0.0323 | 0.2585 | lower |
| F\_COL | 16 | 5093 | 1 | 0.0196 | 0.0196 | 0.1767 | lower |
| F\_COL | 18 | 2958 | 1 | 0.0338 | 0.0338 | 0.2366 | lower |
| F\_COL | 20 | 3797 | 1 | 0.0263 | 0.0263 | 0.2107 | lower |
| F\_COL | 21 | 2937 | 1 | 0.0340 | 0.0340 | 0.2383 | lower |
| F\_COL | 23 | 1756 | 1 | 0.0569 | 0.0569 | 0.2847 | lower |
| F\_COL | 24 | 2314 | 1 | 0.0432 | 0.0432 | 0.2593 | lower |
| F\_COL | 30 | 3383 | 1 | 0.0296 | 0.0296 | 0.2956 | lower |
| F\_COL | 32 | 1274 | 1 | 0.0785 | 0.0785 | 0.3140 | lower |
| F\_MIL | 3 | 5253 | 1 | 0.0190 | 0.0190 | 0.1333 | lower |
| F\_MIL | 4 | 5657 | 1 | 0.0177 | 0.0177 | 0.1237 | lower |
| F\_MIL | 11 | 3095 | 1 | 0.0323 | 0.0323 | 0.1616 | lower |
| F\_MIL | 12 | 5620 | 1 | 0.0178 | 0.0178 | 0.1246 | lower |
| F\_MIL | 13 | 5979 | 1 | 0.0167 | 0.0167 | 0.1338 | lower |
| F\_MIL | 17 | 2070 | 1 | 0.0483 | 0.0483 | 0.1449 | lower |
| F\_MIL | 22 | 3447 | 1 | 0.0290 | 0.0290 | 0.1451 | lower |
| F\_MIL | 24 | 2314 | 1 | 0.0432 | 0.0432 | 0.2161 | lower |
| F\_MIL | 25 | 2980 | 1 | 0.0336 | 0.0336 | 0.1678 | lower |
| F\_MIL | 27 | 1997 | 1 | 0.0501 | 0.0501 | 0.2003 | lower |
| F\_MIL | 31 | 2772 | 1 | 0.0361 | 0.0361 | 0.1804 | lower |
| F\_MIL | 34 | 1094 | 1 | 0.0914 | 0.0914 | 0.3656 | lower |
| F\_DER | 2 | 4869 | 1 | 0.0205 | 0.0205 | 0.1438 | lower |
| F\_DER | 6 | 6408 | 1 | 0.0156 | 0.0156 | 0.1092 | lower |
| F\_DER | 15 | 3245 | 1 | 0.0308 | 0.0308 | 0.1541 | lower |
| F\_DER | 21 | 2937 | 1 | 0.0340 | 0.0340 | 0.1702 | lower |
| F\_DER | 22 | 3447 | 1 | 0.0290 | 0.0290 | 0.1741 | lower |
| F\_DER | 24 | 2314 | 1 | 0.0432 | 0.0432 | 0.2161 | lower |
| F\_DER | 26 | 4047 | 1 | 0.0247 | 0.0247 | 0.1235 | lower |
| F\_DER | 28 | 3291 | 1 | 0.0304 | 0.0304 | 0.1519 | lower |
| F\_DER | 29 | 2272 | 1 | 0.0440 | 0.0440 | 0.2201 | lower |
| F\_DER | 30 | 3383 | 5 | 0.1478 | 0.0296 | 0.1478 | higher |
| F\_DER | 31 | 2772 | 1 | 0.0361 | 0.0361 | 0.1804 | lower |
| F\_DER | 34 | 1094 | 1 | 0.0914 | 0.0914 | 0.2742 | lower |
| F\_BUT | 2 | 4869 | 1 | 0.0205 | 0.0205 | 0.1027 | lower |
| F\_BUT | 4 | 5657 | 1 | 0.0177 | 0.0177 | 0.0884 | lower |
| F\_BUT | 11 | 3095 | 1 | 0.0323 | 0.0323 | 0.1292 | lower |
| F\_BUT | 14 | 4284 | 1 | 0.0233 | 0.0233 | 0.1167 | lower |
| F\_BUT | 15 | 3245 | 1 | 0.0308 | 0.0308 | 0.1541 | lower |
| F\_BUT | 31 | 2772 | 4 | 0.1443 | 0.0361 | 0.1443 | higher |
| F\_BUT | 32 | 1274 | 1 | 0.0785 | 0.0785 | 0.2355 | lower |
| F\_BUT | 33 | 3255 | 1 | 0.0307 | 0.0307 | 0.1229 | lower |
| F\_FRH | 4 | 5657 | 1 | 0.0177 | 0.0177 | 0.1414 | lower |
| F\_FRH | 8 | 6666 | 1 | 0.0150 | 0.0150 | 0.1650 | lower |
| F\_FRH | 16 | 5093 | 1 | 0.0196 | 0.0196 | 0.1767 | lower |
| F\_FRH | 18 | 2958 | 1 | 0.0338 | 0.0338 | 0.2028 | lower |
| F\_FRH | 20 | 3797 | 1 | 0.0263 | 0.0263 | 0.2107 | lower |
| F\_FRH | 21 | 2937 | 1 | 0.0340 | 0.0340 | 0.2383 | lower |
| F\_FRH | 22 | 3447 | 1 | 0.0290 | 0.0290 | 0.2031 | lower |
| F\_FRH | 23 | 1756 | 1 | 0.0569 | 0.0569 | 0.2847 | lower |
| F\_FRH | 24 | 2314 | 1 | 0.0432 | 0.0432 | 0.2593 | lower |
| F\_FRH | 29 | 2272 | 1 | 0.0440 | 0.0440 | 0.2641 | lower |
| F\_NIM | 3 | 5253 | 1 | 0.0190 | 0.0190 | 0.1523 | lower |
| F\_NIM | 10 | 5351 | 1 | 0.0187 | 0.0187 | 0.1869 | lower |
| F\_NIM | 13 | 5979 | 1 | 0.0167 | 0.0167 | 0.1505 | lower |
| F\_NIM | 27 | 1997 | 1 | 0.0501 | 0.0501 | 0.2504 | lower |
| F\_NIM | 30 | 3383 | 1 | 0.0296 | 0.0296 | 0.2069 | lower |
| F\_NIM | 33 | 3255 | 1 | 0.0307 | 0.0307 | 0.2151 | lower |
| F\_NIM | 34 | 1094 | 1 | 0.0914 | 0.0914 | 0.3656 | lower |
| F\_MKH | 2 | 4869 | 1 | 0.0205 | 0.0205 | 0.1643 | lower |
| F\_MKH | 4 | 5657 | 1 | 0.0177 | 0.0177 | 0.1237 | lower |
| F\_MKH | 10 | 5351 | 1 | 0.0187 | 0.0187 | 0.1308 | lower |
| F\_MKH | 11 | 3095 | 1 | 0.0323 | 0.0323 | 0.1939 | lower |
| F\_MKH | 12 | 5620 | 1 | 0.0178 | 0.0178 | 0.1246 | lower |
| F\_MKH | 15 | 3245 | 1 | 0.0308 | 0.0308 | 0.2157 | lower |
| F\_MKH | 21 | 2937 | 1 | 0.0340 | 0.0340 | 0.1702 | lower |
| F\_MKH | 23 | 1756 | 1 | 0.0569 | 0.0569 | 0.1708 | lower |
| F\_MKH | 26 | 4047 | 1 | 0.0247 | 0.0247 | 0.1483 | lower |
| F\_MKH | 29 | 2272 | 1 | 0.0440 | 0.0440 | 0.2201 | lower |
| F\_MKH | 30 | 3383 | 1 | 0.0296 | 0.0296 | 0.1774 | lower |
| F\_MKH | 32 | 1274 | 1 | 0.0785 | 0.0785 | 0.2355 | lower |
| F\_MKH | 34 | 1094 | 1 | 0.0914 | 0.0914 | 0.2742 | lower |
| F\_STN | 6 | 6408 | 1 | 0.0156 | 0.0156 | 0.1092 | lower |
| F\_STN | 10 | 5351 | 1 | 0.0187 | 0.0187 | 0.1308 | lower |
| F\_STN | 12 | 5620 | 1 | 0.0178 | 0.0178 | 0.1246 | lower |
| F\_STN | 14 | 4284 | 1 | 0.0233 | 0.0233 | 0.1401 | lower |
| F\_STN | 16 | 5093 | 1 | 0.0196 | 0.0196 | 0.1571 | lower |
| F\_STN | 21 | 2937 | 1 | 0.0340 | 0.0340 | 0.1702 | lower |
| F\_STN | 22 | 3447 | 1 | 0.0290 | 0.0290 | 0.2031 | lower |
| F\_STN | 24 | 2314 | 1 | 0.0432 | 0.0432 | 0.2593 | lower |
| F\_STN | 26 | 4047 | 1 | 0.0247 | 0.0247 | 0.1730 | lower |
| F\_STN | 31 | 2772 | 1 | 0.0361 | 0.0361 | 0.2165 | lower |
| F\_TOU | 3 | 5253 | 1 | 0.0190 | 0.0190 | 0.1713 | lower |
| F\_TOU | 6 | 6408 | 1 | 0.0156 | 0.0156 | 0.1248 | lower |
| F\_TOU | 7 | 5707 | 1 | 0.0175 | 0.0175 | 0.1227 | lower |
| F\_TOU | 14 | 4284 | 1 | 0.0233 | 0.0233 | 0.1401 | lower |
| F\_TOU | 21 | 2937 | 1 | 0.0340 | 0.0340 | 0.2383 | lower |
| F\_TOU | 22 | 3447 | 1 | 0.0290 | 0.0290 | 0.1451 | lower |
| F\_TOU | 24 | 2314 | 1 | 0.0432 | 0.0432 | 0.3025 | lower |
| F\_TOU | 25 | 2980 | 1 | 0.0336 | 0.0336 | 0.2013 | lower |
| F\_TOU | 27 | 1997 | 1 | 0.0501 | 0.0501 | 0.2504 | lower |
| F\_TOU | 28 | 3291 | 1 | 0.0304 | 0.0304 | 0.1519 | lower |
| F\_TOU | 30 | 3383 | 1 | 0.0296 | 0.0296 | 0.1774 | lower |
| F\_TOU | 33 | 3255 | 1 | 0.0307 | 0.0307 | 0.1843 | lower |
| F\_MRH | 3 | 5253 | 1 | 0.0190 | 0.0190 | 0.1713 | lower |
| F\_MRH | 5 | 6268 | 1 | 0.0160 | 0.0160 | 0.1276 | lower |
| F\_MRH | 6 | 6408 | 1 | 0.0156 | 0.0156 | 0.1248 | lower |
| F\_MRH | 7 | 5707 | 1 | 0.0175 | 0.0175 | 0.1227 | lower |
| F\_MRH | 8 | 6666 | 1 | 0.0150 | 0.0150 | 0.1350 | lower |
| F\_MRH | 12 | 5620 | 1 | 0.0178 | 0.0178 | 0.1068 | lower |
| F\_MRH | 14 | 4284 | 1 | 0.0233 | 0.0233 | 0.1401 | lower |
| F\_MRH | 15 | 3245 | 1 | 0.0308 | 0.0308 | 0.1541 | lower |
| F\_MRH | 21 | 2937 | 1 | 0.0340 | 0.0340 | 0.2043 | lower |
| F\_MRH | 23 | 1756 | 1 | 0.0569 | 0.0569 | 0.2278 | lower |
| F\_MRH | 29 | 2272 | 1 | 0.0440 | 0.0440 | 0.2201 | lower |
| F\_MRH | 31 | 2772 | 1 | 0.0361 | 0.0361 | 0.1443 | lower |
| F\_MER | 2 | 4869 | 1 | 0.0205 | 0.0205 | 0.1438 | lower |
| F\_MER | 5 | 6268 | 1 | 0.0160 | 0.0160 | 0.1117 | lower |
| F\_MER | 7 | 5707 | 1 | 0.0175 | 0.0175 | 0.1402 | lower |
| F\_MER | 8 | 6666 | 1 | 0.0150 | 0.0150 | 0.1200 | lower |
| F\_MER | 14 | 4284 | 1 | 0.0233 | 0.0233 | 0.1401 | lower |
| F\_MER | 17 | 2070 | 1 | 0.0483 | 0.0483 | 0.2415 | lower |
| F\_MER | 20 | 3797 | 1 | 0.0263 | 0.0263 | 0.1580 | lower |
| F\_MER | 21 | 2937 | 1 | 0.0340 | 0.0340 | 0.1702 | lower |
| F\_MER | 22 | 3447 | 1 | 0.0290 | 0.0290 | 0.1451 | lower |
| F\_MER | 27 | 1997 | 1 | 0.0501 | 0.0501 | 0.2003 | lower |
| L\_USR | 2 | 4869 | 1 | 0.0205 | 0.0205 | 0.1643 | lower |
| L\_USR | 11 | 3095 | 1 | 0.0323 | 0.0323 | 0.2262 | lower |
| L\_USR | 15 | 3245 | 1 | 0.0308 | 0.0308 | 0.2465 | lower |
| L\_USR | 18 | 2958 | 1 | 0.0338 | 0.0338 | 0.2366 | lower |
| L\_USR | 26 | 4047 | 1 | 0.0247 | 0.0247 | 0.2224 | lower |
| L\_USR | 27 | 1997 | 1 | 0.0501 | 0.0501 | 0.2504 | lower |
| L\_USR | 28 | 3291 | 1 | 0.0304 | 0.0304 | 0.1823 | lower |
| L\_USR | 29 | 2272 | 1 | 0.0440 | 0.0440 | 0.2201 | lower |
| W\_USR | 9 | 6000 | 1 | 0.0167 | 0.0167 | 0.1500 | lower |
| W\_USR | 12 | 5620 | 1 | 0.0178 | 0.0178 | 0.1423 | lower |
| W\_USR | 16 | 5093 | 1 | 0.0196 | 0.0196 | 0.1571 | lower |
| W\_USR | 18 | 2958 | 1 | 0.0338 | 0.0338 | 0.2028 | lower |
| W\_USR | 20 | 3797 | 1 | 0.0263 | 0.0263 | 0.1844 | lower |
| W\_USR | 21 | 2937 | 5 | 0.1702 | 0.0340 | 0.1702 | higher |
| W\_USR | 25 | 2980 | 1 | 0.0336 | 0.0336 | 0.2013 | lower |
| W\_USR | 27 | 1997 | 1 | 0.0501 | 0.0501 | 0.2003 | lower |
| W\_USR | 31 | 2772 | 1 | 0.0361 | 0.0361 | 0.2165 | lower |
| W\_USR | 32 | 1274 | 1 | 0.0785 | 0.0785 | 0.3140 | lower |
| S\_MIL | 4 | 5657 | 1 | 0.0177 | 0.0177 | 0.1591 | lower |
| S\_MIL | 11 | 3095 | 1 | 0.0323 | 0.0323 | 0.2262 | lower |
| S\_MIL | 19 | 5071 | 1 | 0.0197 | 0.0197 | 0.1578 | lower |
| S\_MIL | 21 | 2937 | 1 | 0.0340 | 0.0340 | 0.1702 | lower |
| S\_MIL | 27 | 1997 | 1 | 0.0501 | 0.0501 | 0.2003 | lower |
| S\_MIL | 29 | 2272 | 1 | 0.0440 | 0.0440 | 0.2201 | lower |
| S\_MIL | 30 | 3383 | 1 | 0.0296 | 0.0296 | 0.2069 | lower |
| S\_DER | 8 | 6666 | 1 | 0.0150 | 0.0150 | 0.2700 | lower |
| S\_DER | 21 | 2937 | 1 | 0.0340 | 0.0340 | 0.2724 | lower |
| S\_DER | 22 | 3447 | 1 | 0.0290 | 0.0290 | 0.3191 | lower |
| S\_DER | 24 | 2314 | 1 | 0.0432 | 0.0432 | 0.3889 | lower |
| S\_DER | 28 | 3291 | 1 | 0.0304 | 0.0304 | 0.2735 | lower |
| S\_DER | 31 | 2772 | 1 | 0.0361 | 0.0361 | 0.2886 | lower |
| S\_DER | 32 | 1274 | 1 | 0.0785 | 0.0785 | 0.4710 | lower |
| S\_DER | 34 | 1094 | 1 | 0.0914 | 0.0914 | 0.4570 | lower |
| S\_BUT | 2 | 4869 | 1 | 0.0205 | 0.0205 | 0.2259 | lower |
| S\_BUT | 14 | 4284 | 1 | 0.0233 | 0.0233 | 0.2101 | lower |
| S\_BUT | 15 | 3245 | 1 | 0.0308 | 0.0308 | 0.2773 | lower |
| S\_BUT | 22 | 3447 | 1 | 0.0290 | 0.0290 | 0.2031 | lower |
| S\_BUT | 27 | 1997 | 1 | 0.0501 | 0.0501 | 0.2504 | lower |
| S\_BUT | 32 | 1274 | 1 | 0.0785 | 0.0785 | 0.3140 | lower |
| S\_FRH | 5 | 6268 | 1 | 0.0160 | 0.0160 | 0.0638 | lower |
| S\_FRH | 6 | 6408 | 1 | 0.0156 | 0.0156 | 0.0780 | lower |
| S\_FRH | 11 | 3095 | 1 | 0.0323 | 0.0323 | 0.0969 | lower |
| S\_FRH | 13 | 5979 | 1 | 0.0167 | 0.0167 | 0.1004 | lower |
| S\_FRH | 15 | 3245 | 1 | 0.0308 | 0.0308 | 0.0924 | lower |
| S\_FRH | 16 | 5093 | 1 | 0.0196 | 0.0196 | 0.0785 | lower |
| S\_FRH | 17 | 2070 | 1 | 0.0483 | 0.0483 | 0.1449 | lower |
| S\_FRH | 27 | 1997 | 1 | 0.0501 | 0.0501 | 0.1502 | lower |
| S\_FRH | 30 | 3383 | 1 | 0.0296 | 0.0296 | 0.1182 | lower |

Table S13: Comparison across run/tributary group of the mean +/- sd, minimum, and maximum percent of loci on a chromosome that are only polymorphic for a set of individuals from the same tributary of the same run type.

| POLYMORPHIC | mean | sd | min | max |
| --- | --- | --- | --- | --- |
| F\_COL | 0.07 | 0.04 | 0.02 | 0.15 |
| F\_MIL | 0.05 | 0.02 | 0.02 | 0.12 |
| F\_DER | 0.04 | 0.03 | 0.02 | 0.15 |
| F\_BUT | 0.04 | 0.03 | 0.02 | 0.14 |
| F\_FRH | 0.07 | 0.04 | 0.02 | 0.18 |
| F\_NIM | 0.06 | 0.03 | 0.02 | 0.14 |
| F\_MKH | 0.05 | 0.03 | 0.02 | 0.12 |
| F\_STN | 0.05 | 0.02 | 0.02 | 0.10 |
| F\_TOU | 0.04 | 0.02 | 0.02 | 0.10 |
| F\_MRH | 0.05 | 0.03 | 0.02 | 0.14 |
| F\_MER | 0.05 | 0.03 | 0.02 | 0.13 |
| L\_USR | 0.06 | 0.03 | 0.02 | 0.11 |
| W\_USR | 0.06 | 0.05 | 0.02 | 0.18 |
| S\_MIL | 0.05 | 0.02 | 0.02 | 0.09 |
| S\_DER | 0.10 | 0.05 | 0.02 | 0.27 |
| S\_BUT | 0.07 | 0.03 | 0.02 | 0.12 |
| S\_FRH | 0.03 | 0.01 | 0.02 | 0.05 |

Table S14: Number of chromosomes with significantly more/less loci with private polymorphisms than expected.

| GRP | sign | n |
| --- | --- | --- |
| F\_MKH | lower | 13 |
| F\_MIL | lower | 12 |
| F\_TOU | lower | 12 |
| F\_MRH | lower | 12 |
| F\_COL | lower | 11 |
| F\_DER | lower | 11 |
| F\_FRH | lower | 10 |
| F\_STN | lower | 10 |
| F\_MER | lower | 10 |
| W\_USR | lower | 9 |
| S\_FRH | lower | 9 |
| L\_USR | lower | 8 |
| S\_DER | lower | 8 |
| F\_BUT | lower | 7 |
| F\_NIM | lower | 7 |
| S\_MIL | lower | 7 |
| S\_BUT | lower | 6 |
| F\_DER | higher | 1 |
| F\_BUT | higher | 1 |
| W\_USR | higher | 1 |

Table S15: Number of run/trib groups a chromosome has been flagged as having significantly more/less loci with private polymorphisms than expected.

| CHROMOSOME | sign | n |
| --- | --- | --- |
| 21 | lower | 10 |
| 27 | lower | 9 |
| 22 | lower | 8 |
| 11 | lower | 7 |
| 15 | lower | 7 |
| 24 | lower | 7 |
| 2 | lower | 6 |
| 6 | lower | 6 |
| 14 | lower | 6 |
| 29 | lower | 6 |
| 30 | lower | 6 |
| 31 | lower | 6 |
| 32 | lower | 6 |
| 4 | lower | 5 |
| 12 | lower | 5 |
| 16 | lower | 5 |
| 34 | lower | 5 |
| 3 | lower | 4 |
| 8 | lower | 4 |
| 10 | lower | 4 |
| 18 | lower | 4 |
| 20 | lower | 4 |
| 23 | lower | 4 |
| 26 | lower | 4 |
| 28 | lower | 4 |
| 5 | lower | 3 |
| 7 | lower | 3 |
| 13 | lower | 3 |
| 17 | lower | 3 |
| 25 | lower | 3 |
| 33 | lower | 3 |
| 9 | lower | 1 |
| 19 | lower | 1 |
| 21 | higher | 1 |
| 30 | higher | 1 |
| 31 | higher | 1 |

#### 4.10 Private alleles

**Private alleles** are alleles that only occur within a given set of individuals; this can be a locus that is only variable in one set of individuals (all other individuals are fixed for the alternate allele(s)) or it could be a locus with multiple alleles, that are variable in other sets of individuals (i.e. heterozygotes & homozygotes occur) but one (or more) of those alleles only exists within a given set of individuals. Not all individuals of the group will have the private allele, i.e. they may be more or less common.

**Comparison of private alleles by tributary within run type**

Number of private alles identified in individuals of the same run type within a tributary.

Number of private alleles occurring in each set of individuals grouped by run type within tributaries.

| population | n |
| --- | --- |
| S\_DER | 267 |
| S\_BUT | 176 |
| F\_COL | 175 |
| F\_NIM | 161 |
| F\_FRH | 159 |
| S\_MIL | 145 |
| F\_MKH | 136 |
| L\_USR | 129 |
| W\_USR | 122 |
| F\_MER | 113 |
| F\_TOU | 104 |
| F\_MIL | 100 |
| F\_STN | 90 |
| F\_MRH | 87 |
| F\_DER | 82 |
| F\_BUT | 67 |
| S\_FRH | 34 |

There is a relationship between sample size and the number of private alleles found making it different to compare levels overall (many private alleles are rare so for smaller sample sizes they might not be recovered); though it is important to note that all run/tributary groups do exhibit private allele, i.e. harbor unique genetic material.

Table S11: Summary stats of number of individuals carrying private alleles (individuals grouped by run type within tributaries.

| population | mean | sd | max | min |
| --- | --- | --- | --- | --- |
| W\_USR | 4.07 | 5.22 | 25 | 1 |
| S\_BUT | 1.34 | 1.11 | 10 | 1 |
| S\_DER | 1.17 | 0.53 | 5 | 1 |
| F\_COL | 1.17 | 0.51 | 4 | 1 |
| L\_USR | 1.11 | 0.42 | 4 | 1 |
| F\_MIL | 1.07 | 0.29 | 3 | 1 |
| F\_NIM | 1.07 | 0.28 | 3 | 1 |
| S\_MIL | 1.17 | 0.43 | 3 | 1 |
| F\_BUT | 1.01 | 0.12 | 2 | 1 |
| F\_DER | 1.02 | 0.16 | 2 | 1 |
| F\_FRH | 1.09 | 0.29 | 2 | 1 |
| F\_MER | 1.03 | 0.16 | 2 | 1 |
| F\_MKH | 1.04 | 0.21 | 2 | 1 |
| F\_MRH | 1.03 | 0.18 | 2 | 1 |
| F\_STN | 1.03 | 0.18 | 2 | 1 |
| F\_TOU | 1.11 | 0.31 | 2 | 1 |
| S\_FRH | 1.03 | 0.17 | 2 | 1 |

Compare the distribution of loci with private alleles across the genome.

Determine the % loci of a given chromosome that are only polymorphic for a set of individuals from the same tributary of the same run type.

| CHROMOSOME | total | F\_COL | F\_MIL | F\_DER | F\_BUT | F\_FRH | F\_NIM | F\_MKH | F\_STN | F\_TOU | F\_MRH | F\_MER | L\_USR | W\_USR | S\_MIL | S\_DER | S\_BUT | S\_FRH |
| --- | --- | --- | --- | --- | --- | --- | --- | --- | --- | --- | --- | --- | --- | --- | --- | --- | --- | --- |
| 1 | 8167 | 0.12 | 0.09 | 0.09 | 0.05 | 0.16 | 0.23 | 0.12 | 0.06 | 0.02 | 0.06 | 0.07 | 0.04 | 0.07 | 0.10 | 0.17 | 0.17 | 0.04 |
| 2 | 4869 | 0.08 | 0.06 | 0.02 | 0.04 | 0.10 | 0.12 | 0.06 | NA | 0.08 | 0.06 | 0.06 | 0.02 | NA | 0.02 | 0.14 | 0.12 | NA |
| 3 | 5253 | 0.23 | 0.08 | 0.04 | NA | 0.08 | 0.10 | 0.11 | 0.06 | 0.08 | 0.04 | 0.08 | 0.08 | 0.06 | 0.13 | 0.19 | 0.17 | NA |
| 4 | 5657 | 0.11 | 0.05 | 0.05 | 0.05 | 0.04 | 0.09 | 0.02 | 0.05 | 0.09 | 0.09 | 0.14 | 0.14 | 0.16 | 0.05 | 0.19 | 0.09 | 0.02 |
| 5 | 6268 | 0.22 | 0.02 | 0.05 | 0.03 | 0.11 | 0.11 | 0.05 | 0.11 | 0.02 | 0.02 | 0.03 | 0.10 | 0.16 | 0.06 | 0.18 | 0.14 | 0.02 |
| 6 | 6408 | 0.05 | 0.05 | 0.06 | 0.06 | 0.11 | 0.14 | 0.16 | 0.05 | 0.03 | 0.02 | 0.09 | 0.06 | 0.08 | 0.11 | 0.17 | 0.14 | 0.03 |
| 7 | 5707 | 0.14 | 0.09 | 0.02 | NA | 0.07 | 0.07 | 0.07 | 0.09 | 0.04 | 0.04 | 0.04 | 0.09 | 0.11 | 0.09 | 0.25 | 0.11 | 0.07 |
| 8 | 6666 | 0.09 | 0.09 | 0.05 | 0.08 | 0.09 | 0.11 | 0.18 | 0.11 | 0.09 | 0.03 | 0.06 | 0.03 | 0.14 | 0.12 | 0.11 | 0.11 | 0.06 |
| 9 | 6000 | 0.23 | 0.07 | 0.05 | 0.08 | 0.15 | 0.05 | 0.05 | 0.12 | 0.07 | 0.03 | 0.08 | 0.07 | 0.03 | 0.13 | 0.17 | 0.08 | NA |
| 10 | 5351 | 0.09 | 0.04 | 0.04 | 0.07 | 0.09 | 0.06 | 0.11 | 0.02 | 0.06 | 0.07 | 0.02 | 0.15 | 0.13 | 0.06 | 0.19 | 0.09 | 0.04 |
| 11 | 3095 | 0.03 | 0.03 | 0.03 | 0.13 | 0.06 | 0.13 | 0.13 | 0.06 | 0.16 | 0.13 | 0.03 | 0.03 | 0.03 | 0.03 | 0.13 | 0.10 | 0.10 |
| 12 | 5620 | 0.12 | 0.04 | 0.07 | 0.02 | 0.12 | 0.12 | 0.07 | 0.05 | 0.07 | 0.07 | 0.09 | 0.04 | 0.11 | 0.09 | 0.14 | 0.14 | 0.02 |
| 13 | 5979 | 0.13 | 0.03 | NA | NA | 0.12 | 0.05 | 0.07 | 0.07 | 0.03 | NA | 0.08 | 0.08 | 0.03 | 0.12 | 0.12 | 0.05 | 0.02 |
| 14 | 4284 | 0.16 | 0.09 | NA | 0.05 | 0.14 | 0.07 | 0.16 | 0.05 | 0.09 | 0.05 | 0.05 | 0.07 | 0.07 | 0.05 | 0.16 | 0.05 | NA |
| 15 | 3245 | 0.06 | NA | 0.03 | 0.03 | 0.06 | 0.06 | 0.18 | 0.09 | 0.12 | 0.06 | 0.06 | 0.06 | NA | NA | 0.25 | 0.06 | 0.06 |
| 16 | 5093 | 0.12 | 0.08 | 0.14 | NA | 0.06 | 0.18 | 0.08 | 0.02 | 0.16 | 0.04 | 0.04 | 0.10 | 0.04 | NA | 0.12 | 0.08 | 0.02 |
| 17 | 2070 | 0.10 | 0.05 | NA | NA | NA | 0.14 | NA | NA | 0.10 | NA | 0.05 | 0.05 | NA | 0.14 | 0.14 | 0.14 | 0.05 |
| 18 | 2958 | 0.10 | NA | 0.14 | 0.07 | 0.14 | NA | NA | 0.07 | 0.07 | 0.17 | 0.03 | 0.14 | 0.07 | 0.14 | 0.14 | 0.20 | NA |
| 19 | 5071 | 0.10 | 0.20 | 0.08 | 0.08 | 0.10 | 0.20 | 0.12 | 0.04 | 0.02 | 0.14 | 0.10 | 0.08 | 0.12 | 0.04 | 0.16 | 0.06 | NA |
| 20 | 3797 | 0.08 | 0.08 | 0.05 | 0.03 | 0.03 | NA | 0.03 | 0.03 | 0.11 | 0.13 | 0.08 | 0.13 | 0.05 | 0.05 | 0.26 | 0.08 | NA |
| 21 | 2937 | 0.10 | 0.07 | 0.07 | 0.03 | 0.10 | 0.10 | 0.03 | 0.10 | 0.10 | 0.07 | 0.14 | 0.10 | 0.17 | 0.10 | 0.07 | 0.03 | NA |
| 22 | 3447 | 0.15 | 0.06 | 0.03 | 0.03 | 0.12 | 0.03 | 0.03 | 0.06 | 0.06 | 0.03 | 0.03 | 0.15 | 0.17 | 0.12 | 0.15 | 0.09 | NA |
| 23 | 1756 | 0.06 | NA | 0.06 | NA | 0.17 | 0.23 | 0.11 | 0.11 | NA | 0.11 | 0.06 | 0.11 | 0.23 | NA | 0.23 | 0.06 | NA |
| 24 | 2314 | 0.26 | 0.04 | 0.13 | 0.04 | 0.09 | 0.17 | 0.09 | 0.09 | 0.09 | NA | 0.13 | 0.13 | 0.04 | 0.09 | 0.09 | 0.22 | NA |
| 25 | 2980 | NA | 0.17 | 0.03 | NA | 0.07 | 0.10 | 0.13 | NA | 0.10 | 0.03 | 0.20 | 0.13 | 0.03 | 0.17 | 0.23 | 0.13 | NA |
| 26 | 4047 | 0.17 | 0.07 | 0.10 | 0.05 | 0.07 | 0.15 | 0.07 | 0.05 | 0.07 | 0.10 | 0.05 | 0.05 | 0.07 | 0.07 | 0.35 | 0.15 | NA |
| 27 | 1997 | 0.05 | 0.10 | NA | NA | 0.15 | 0.05 | 0.10 | 0.05 | 0.10 | 0.10 | 0.10 | 0.15 | 0.05 | 0.15 | 0.15 | 0.20 | 0.05 |
| 28 | 3291 | 0.21 | 0.12 | 0.03 | NA | 0.18 | NA | 0.03 | 0.06 | 0.03 | 0.06 | 0.06 | 0.03 | NA | 0.12 | 0.12 | 0.21 | NA |
| 29 | 2272 | NA | 0.04 | 0.04 | NA | 0.26 | 0.09 | 0.04 | 0.13 | 0.04 | 0.13 | 0.18 | 0.13 | 0.04 | 0.09 | 0.13 | 0.09 | NA |
| 30 | 3383 | 0.15 | 0.12 | 0.18 | NA | 0.06 | 0.09 | 0.06 | NA | 0.09 | NA | 0.09 | 0.09 | 0.06 | 0.06 | 0.09 | 0.12 | 0.06 |
| 31 | 2772 | NA | 0.04 | 0.04 | 0.18 | 0.22 | 0.04 | 0.22 | 0.04 | NA | 0.07 | 0.11 | 0.04 | 0.07 | 0.11 | 0.14 | 0.04 | NA |
| 32 | 1274 | 0.08 | NA | NA | 0.08 | NA | NA | 0.08 | NA | 0.08 | NA | NA | NA | 0.08 | 0.31 | 0.24 | 0.08 | 0.08 |
| 33 | 3255 | 0.06 | 0.03 | NA | 0.06 | NA | 0.09 | 0.09 | 0.06 | 0.06 | 0.06 | 0.03 | 0.09 | 0.06 | 0.22 | 0.40 | 0.22 | 0.03 |
| 34 | 1094 | NA | 0.09 | 0.09 | NA | 0.18 | 0.09 | 0.18 | NA | NA | NA | NA | NA | 0.18 | NA | 0.09 | 0.09 | NA |

Compare the mean proportion of loci per chromosome that are private polymorphisms across tributaries.

Table S13: Comparison across run/tributary group of the mean +/- sd, minimum, and maximum percent of loci on a chromosome that are only polymorphic for a set of individuals from the same tributary of the same run type.

| population | mean | sd | min | max |
| --- | --- | --- | --- | --- |
| F\_COL | 0.12 | 0.06 | 0.03 | 0.26 |
| F\_MIL | 0.07 | 0.04 | 0.02 | 0.20 |
| F\_DER | 0.06 | 0.04 | 0.02 | 0.18 |
| F\_BUT | 0.06 | 0.04 | 0.02 | 0.18 |
| F\_FRH | 0.11 | 0.05 | 0.03 | 0.26 |
| F\_NIM | 0.11 | 0.05 | 0.03 | 0.23 |
| F\_MKH | 0.10 | 0.05 | 0.02 | 0.22 |
| F\_STN | 0.07 | 0.03 | 0.02 | 0.13 |
| F\_TOU | 0.07 | 0.04 | 0.02 | 0.16 |
| F\_MRH | 0.07 | 0.04 | 0.02 | 0.17 |
| F\_MER | 0.08 | 0.04 | 0.02 | 0.20 |
| L\_USR | 0.09 | 0.04 | 0.02 | 0.15 |
| W\_USR | 0.09 | 0.05 | 0.03 | 0.23 |
| S\_MIL | 0.10 | 0.06 | 0.02 | 0.31 |
| S\_DER | 0.17 | 0.07 | 0.07 | 0.40 |
| S\_BUT | 0.11 | 0.05 | 0.03 | 0.22 |
| S\_FRH | 0.04 | 0.02 | 0.02 | 0.10 |

Generate a random null distribution to determine if loci with private alleles are non-randomly distributed across chromosomes.

Table S14: Chromosomes with significantly higher/lower than expected (outside simulated null distribution) number of fixed loci for each run/tributary group

| GRP | CHROMOSOME | total | n | percent | min | max | sign |
| --- | --- | --- | --- | --- | --- | --- | --- |
| F\_MIL | 34 | 1094 | 1 | 0.0914 | 0.0914 | 0.3656 | lower |
| F\_DER | 34 | 1094 | 1 | 0.0914 | 0.0914 | 0.2742 | lower |
| F\_NIM | 34 | 1094 | 1 | 0.0914 | 0.0914 | 0.3656 | lower |
| S\_DER | 34 | 1094 | 1 | 0.0914 | 0.0914 | 0.5484 | lower |
| S\_BUT | 34 | 1094 | 1 | 0.0914 | 0.0914 | 0.3656 | lower |
| F\_MIL | 33 | 3255 | 1 | 0.0307 | 0.0307 | 0.2765 | lower |
| F\_MER | 33 | 3255 | 1 | 0.0307 | 0.0307 | 0.2458 | lower |
| S\_FRH | 33 | 3255 | 1 | 0.0307 | 0.0307 | 0.1536 | lower |
| F\_COL | 32 | 1274 | 1 | 0.0785 | 0.0785 | 0.4710 | lower |
| F\_BUT | 32 | 1274 | 1 | 0.0785 | 0.0785 | 0.3140 | lower |
| F\_MKH | 32 | 1274 | 1 | 0.0785 | 0.0785 | 0.3925 | lower |
| F\_TOU | 32 | 1274 | 1 | 0.0785 | 0.0785 | 0.2355 | lower |
| W\_USR | 32 | 1274 | 1 | 0.0785 | 0.0785 | 0.4710 | lower |
| S\_BUT | 32 | 1274 | 1 | 0.0785 | 0.0785 | 0.4710 | lower |
| S\_FRH | 32 | 1274 | 1 | 0.0785 | 0.0785 | 0.2355 | lower |
| F\_MIL | 31 | 2772 | 1 | 0.0361 | 0.0361 | 0.2165 | lower |
| F\_DER | 31 | 2772 | 1 | 0.0361 | 0.0361 | 0.2165 | lower |
| F\_BUT | 31 | 2772 | 5 | 0.1804 | 0.0361 | 0.1804 | higher |
| F\_NIM | 31 | 2772 | 1 | 0.0361 | 0.0361 | 0.2886 | lower |
| F\_STN | 31 | 2772 | 1 | 0.0361 | 0.0361 | 0.2165 | lower |
| L\_USR | 31 | 2772 | 1 | 0.0361 | 0.0361 | 0.2525 | lower |
| S\_BUT | 31 | 2772 | 1 | 0.0361 | 0.0361 | 0.3247 | lower |
| F\_MIL | 29 | 2272 | 1 | 0.0440 | 0.0440 | 0.3081 | lower |
| F\_DER | 29 | 2272 | 1 | 0.0440 | 0.0440 | 0.3521 | lower |
| F\_MKH | 29 | 2272 | 1 | 0.0440 | 0.0440 | 0.3521 | lower |
| F\_TOU | 29 | 2272 | 1 | 0.0440 | 0.0440 | 0.3081 | lower |
| W\_USR | 29 | 2272 | 1 | 0.0440 | 0.0440 | 0.3081 | lower |
| F\_DER | 28 | 3291 | 1 | 0.0304 | 0.0304 | 0.2127 | lower |
| F\_MKH | 28 | 3291 | 1 | 0.0304 | 0.0304 | 0.2431 | lower |
| F\_TOU | 28 | 3291 | 1 | 0.0304 | 0.0304 | 0.1823 | lower |
| L\_USR | 28 | 3291 | 1 | 0.0304 | 0.0304 | 0.2735 | lower |
| F\_COL | 27 | 1997 | 1 | 0.0501 | 0.0501 | 0.4006 | lower |
| F\_NIM | 27 | 1997 | 1 | 0.0501 | 0.0501 | 0.4006 | lower |
| F\_STN | 27 | 1997 | 1 | 0.0501 | 0.0501 | 0.3505 | lower |
| W\_USR | 27 | 1997 | 1 | 0.0501 | 0.0501 | 0.4006 | lower |
| S\_FRH | 27 | 1997 | 1 | 0.0501 | 0.0501 | 0.1502 | lower |
| F\_DER | 25 | 2980 | 1 | 0.0336 | 0.0336 | 0.2349 | lower |
| F\_MRH | 25 | 2980 | 1 | 0.0336 | 0.0336 | 0.2349 | lower |
| W\_USR | 25 | 2980 | 1 | 0.0336 | 0.0336 | 0.2685 | lower |
| F\_MIL | 24 | 2314 | 1 | 0.0432 | 0.0432 | 0.3457 | lower |
| F\_BUT | 24 | 2314 | 1 | 0.0432 | 0.0432 | 0.2161 | lower |
| W\_USR | 24 | 2314 | 1 | 0.0432 | 0.0432 | 0.3025 | lower |
| F\_COL | 23 | 1756 | 1 | 0.0569 | 0.0569 | 0.5695 | lower |
| F\_DER | 23 | 1756 | 1 | 0.0569 | 0.0569 | 0.2847 | lower |
| F\_MER | 23 | 1756 | 1 | 0.0569 | 0.0569 | 0.3986 | lower |
| S\_BUT | 23 | 1756 | 1 | 0.0569 | 0.0569 | 0.3986 | lower |
| F\_DER | 22 | 3447 | 1 | 0.0290 | 0.0290 | 0.1741 | lower |
| F\_BUT | 22 | 3447 | 1 | 0.0290 | 0.0290 | 0.1741 | lower |
| F\_NIM | 22 | 3447 | 1 | 0.0290 | 0.0290 | 0.2611 | lower |
| F\_MKH | 22 | 3447 | 1 | 0.0290 | 0.0290 | 0.2611 | lower |
| F\_MRH | 22 | 3447 | 1 | 0.0290 | 0.0290 | 0.2321 | lower |
| F\_MER | 22 | 3447 | 1 | 0.0290 | 0.0290 | 0.2611 | lower |
| F\_BUT | 21 | 2937 | 1 | 0.0340 | 0.0340 | 0.2043 | lower |
| F\_MKH | 21 | 2937 | 1 | 0.0340 | 0.0340 | 0.3064 | lower |
| S\_BUT | 21 | 2937 | 1 | 0.0340 | 0.0340 | 0.3064 | lower |
| F\_BUT | 20 | 3797 | 1 | 0.0263 | 0.0263 | 0.1580 | lower |
| F\_FRH | 20 | 3797 | 1 | 0.0263 | 0.0263 | 0.2897 | lower |
| F\_MKH | 20 | 3797 | 1 | 0.0263 | 0.0263 | 0.2370 | lower |
| F\_STN | 20 | 3797 | 1 | 0.0263 | 0.0263 | 0.1844 | lower |
| F\_TOU | 19 | 5071 | 1 | 0.0197 | 0.0197 | 0.1972 | lower |
| F\_MER | 18 | 2958 | 1 | 0.0338 | 0.0338 | 0.2705 | lower |
| F\_MIL | 17 | 2070 | 1 | 0.0483 | 0.0483 | 0.2415 | lower |
| F\_MER | 17 | 2070 | 1 | 0.0483 | 0.0483 | 0.2415 | lower |
| L\_USR | 17 | 2070 | 1 | 0.0483 | 0.0483 | 0.3382 | lower |
| S\_FRH | 17 | 2070 | 1 | 0.0483 | 0.0483 | 0.1449 | lower |
| F\_DER | 16 | 5093 | 7 | 0.1374 | 0.0196 | 0.1374 | higher |
| F\_STN | 16 | 5093 | 1 | 0.0196 | 0.0196 | 0.1374 | lower |
| S\_FRH | 16 | 5093 | 1 | 0.0196 | 0.0196 | 0.1178 | lower |
| F\_DER | 15 | 3245 | 1 | 0.0308 | 0.0308 | 0.1541 | lower |
| F\_BUT | 15 | 3245 | 1 | 0.0308 | 0.0308 | 0.1849 | lower |
| S\_FRH | 13 | 5979 | 1 | 0.0167 | 0.0167 | 0.0836 | lower |
| F\_BUT | 12 | 5620 | 1 | 0.0178 | 0.0178 | 0.1246 | lower |
| S\_FRH | 12 | 5620 | 1 | 0.0178 | 0.0178 | 0.1068 | lower |
| F\_COL | 11 | 3095 | 1 | 0.0323 | 0.0323 | 0.3554 | lower |
| F\_MIL | 11 | 3095 | 1 | 0.0323 | 0.0323 | 0.2585 | lower |
| F\_DER | 11 | 3095 | 1 | 0.0323 | 0.0323 | 0.2262 | lower |
| F\_MER | 11 | 3095 | 1 | 0.0323 | 0.0323 | 0.2262 | lower |
| L\_USR | 11 | 3095 | 1 | 0.0323 | 0.0323 | 0.2585 | lower |
| W\_USR | 11 | 3095 | 1 | 0.0323 | 0.0323 | 0.2908 | lower |
| S\_MIL | 11 | 3095 | 1 | 0.0323 | 0.0323 | 0.3554 | lower |
| F\_STN | 10 | 5351 | 1 | 0.0187 | 0.0187 | 0.1682 | lower |
| F\_MER | 10 | 5351 | 1 | 0.0187 | 0.0187 | 0.2056 | lower |
| F\_DER | 7 | 5707 | 1 | 0.0175 | 0.0175 | 0.1577 | lower |
| F\_MRH | 6 | 6408 | 1 | 0.0156 | 0.0156 | 0.1561 | lower |
| F\_MIL | 5 | 6268 | 1 | 0.0160 | 0.0160 | 0.1914 | lower |
| F\_TOU | 5 | 6268 | 1 | 0.0160 | 0.0160 | 0.1755 | lower |
| F\_MRH | 5 | 6268 | 1 | 0.0160 | 0.0160 | 0.1755 | lower |
| S\_FRH | 5 | 6268 | 1 | 0.0160 | 0.0160 | 0.0957 | lower |
| F\_MKH | 4 | 5657 | 1 | 0.0177 | 0.0177 | 0.2475 | lower |
| S\_FRH | 4 | 5657 | 1 | 0.0177 | 0.0177 | 0.1061 | lower |
| F\_DER | 2 | 4869 | 1 | 0.0205 | 0.0205 | 0.1643 | lower |
| L\_USR | 2 | 4869 | 1 | 0.0205 | 0.0205 | 0.1848 | lower |
| S\_MIL | 2 | 4869 | 1 | 0.0205 | 0.0205 | 0.2054 | lower |
| F\_NIM | 1 | 8167 | 19 | 0.2326 | 0.0245 | 0.2204 | higher |

Table S15: Number of run/trib groups a chromosome has been flagged as having significantly more/less loci with exclusive polymorphisms than expected.

| CHROMOSOME | sign | n |
| --- | --- | --- |
| 11 | lower | 7 |
| 32 | lower | 7 |
| 22 | lower | 6 |
| 31 | lower | 6 |
| 27 | lower | 5 |
| 29 | lower | 5 |
| 34 | lower | 5 |
| 5 | lower | 4 |
| 17 | lower | 4 |
| 20 | lower | 4 |
| 23 | lower | 4 |
| 28 | lower | 4 |
| 2 | lower | 3 |
| 21 | lower | 3 |
| 24 | lower | 3 |
| 25 | lower | 3 |
| 33 | lower | 3 |
| 4 | lower | 2 |
| 10 | lower | 2 |
| 12 | lower | 2 |
| 15 | lower | 2 |
| 16 | lower | 2 |
| 1 | higher | 1 |
| 6 | lower | 1 |
| 7 | lower | 1 |
| 13 | lower | 1 |
| 16 | higher | 1 |
| 18 | lower | 1 |
| 19 | lower | 1 |
| 31 | higher | 1 |

Number of chromosomes with significantly more/less loci with exclusive polymorphisms than expected.

| GRP | sign | n |
| --- | --- | --- |
| F\_DER | lower | 11 |
| S\_FRH | lower | 9 |
| F\_MIL | lower | 8 |
| F\_BUT | lower | 7 |
| F\_MKH | lower | 7 |
| F\_MER | lower | 7 |
| W\_USR | lower | 6 |
| F\_STN | lower | 5 |
| F\_TOU | lower | 5 |
| L\_USR | lower | 5 |
| S\_BUT | lower | 5 |
| F\_COL | lower | 4 |
| F\_NIM | lower | 4 |
| F\_MRH | lower | 4 |
| S\_MIL | lower | 2 |
| F\_DER | higher | 1 |
| F\_BUT | higher | 1 |
| F\_FRH | lower | 1 |
| F\_NIM | higher | 1 |
| S\_DER | lower | 1 |

##### 4.10.1 Session Information

⊕Package versions used for this analysis (R 3.6.0).

```
##             package loadedversion
## ade4           ade4        1.7-16
## adegenet   adegenet         2.1.3
## ape             ape         5.4-1
## assigner   assigner         0.5.7
## coin           coin         1.3-1
## dplyr         dplyr         1.0.2
## forcats     forcats         0.5.0
## ggplot2     ggplot2         3.3.3
## glue           glue         1.4.2
## hierfstat hierfstat         0.5-7
## httr           httr         1.4.2
## Imap           Imap          1.32
## knitr         knitr          1.30
## magrittr   magrittr         2.0.1
## pegas         pegas          0.14
## plyr           plyr         1.8.6
## poppr         poppr         2.8.7
## purrr         purrr         0.3.4
## radiator   radiator         1.1.6
## readr         readr         1.4.0
## stringr     stringr         1.4.0
## survival   survival         3.2-7
## tibble       tibble         3.0.4
## tidyr         tidyr         1.1.2
## tidyverse tidyverse         1.3.0
## tint           tint         0.1.3
## tufte         tufte           0.9
## UpSetR       UpSetR         1.4.0
## vcfR           vcfR        1.12.0
```
